# Protein phosphatase 2C domain contributes to the pathobiological function of adenylyl cyclase in *Cryptococcus neoformans*

**DOI:** 10.64898/2025.12.10.693573

**Authors:** Soojin Yu, Seong-Ryong Yu, Myung Kyung Choi, Jae-Hyung Jin, Vikas Yadav, Hee Seong Choi, Eui-Seong Kim, Hyun-Woo Kim, Jae-Seok Jeong, Do-Kyun Kim, Joseph Heitman, Hyun-Soo Cho, Kyung-Tae Lee, Yong-Sun Bahn

**Author notes:** Present address: Aligned Genetics, Gunpo 15850, Republic of Korea. These authors contributed equally (Soojin Yu, Seong-Ryong Yu, Myung Kyung Choi, Jae-Hyung Jin, Vikas Yadav). **Correspondences and requests for materials should be addressed to YSB (****), KTL (****), HSC (****), JH (**).

## Abstract

Adenylyl cyclase produces cyclic adenosine monophosphate, a signalling molecule that controls fungal development and virulence. Fungal adenylyl cyclases are distinguished by the presence of a conserved protein phosphatase 2C domain, whose function remains unknown. Here we show that the protein phosphatase 2C domain of Cac1, the adenylyl cyclase of the human pathogen *Cryptococcus neoformans*, has an unusual structure but functions as a metal-dependent serine/threonine phosphatase. Deletion of this domain or the adenylyl cyclase catalytic domain demonstrated that the protein phosphatase 2C domain is required for full Cac1 activity, influencing melanin and capsule synthesis, sexual differentiation, titan cell formation, and cell wall integrity. Notably, loss of the protein phosphatase 2C domain induces type 2 helper T-cell-biased immunity and extensive pulmonary damage yet does not cause mortality in mice. Integrated transcriptomic and phosphoproteomic analyses further revealed shared and domain-specific signalling outputs. Together, these findings define a pivotal role for the adenylyl cyclase-linked protein phosphatase 2C domain in *C. neoformans* pathobiology.

## INTRODUCTION

Cyclic adenosine monophosphate (cAMP) is a fundamental second messenger that regulates diverse cellular processes across all domains of life. In bacteria, cAMP plays key roles in carbon catabolite repression, motility and chemotaxis, biofilm formation, and the expression of virulence factors^1^. In humans, cAMP regulates cellular metabolism, gene expression, cell proliferation and differentiation, immune responses, neuronal signalling, hormone release, and sensory transduction^2,3^. In fungi, the cAMP pathway functions as a central regulatory hub controlling growth, differentiation, morphogenetic development, stress adaptation, and virulence^4,5^. For example, in *Candida albicans*, an opportunistic human pathogen and member of the gastrointestinal microbiota, the cAMP pathway is essential for yeast-to-hyphae transition and virulence^6^. In *Cryptococcus neoformans*, the causative agent of life-threatening fungal meningoencephalitis, cAMP signalling governs the production of two major virulence factors, capsule and melanin, as well as sexual differentiation and stress responses^7,8^. In *Aspergillus fumigatus*, a filamentous fungus responsible for invasive aspergillosis, the pathway is implicated in conidiation, pigmentation, toxin production, and virulence^9,10^. In plant fungal pathogens such as *Magnaporthe oryzae* and *Fusarium graminearum*, cAMP signalling controls growth, differentiation, and pathogenicity^11,12^.

Adenylyl cyclases (ACs) are key enzymes that catalyse the conversion of adenosine triphosphate (ATP) to cAMP. In humans and other eukaryotes, ACs are primarily activated through G protein-coupled receptors (GPCRs) and other signalling molecules. Upon ligand binding, such as neurotransmitters, hormones, or amino acids, GPCRs undergo conformational changes that activate heterotrimeric G proteins (Gαβγ). The GTP-bound Gα subunit then modulates AC activity, either stimulating or inhibiting cAMP production^13^. In fungi, ACs can also be activated by small GTPase proteins such as Ras or by small molecules like bicarbonate, which directly bind to ACs and promote cAMP synthesis^14^. Although less common in prokaryotes, certain bacteria possess G-protein-like signalling mechanisms capable of modulating AC activity in response to environmental signals^15,16^. In eukaryotic cells, newly synthesised cAMP binds to the regulatory subunits of protein kinase A (PKA), relieving their inhibition of the catalytic subunits^17,18^. The liberated PKA catalytic subunits then phosphorylate downstream transcription factors and effector proteins to elicit cellular responses. To terminate signalling, cAMP is degraded by high– and low-affinity phosphodiesterases (PDEs), providing a negative feedback mechanism^19^. In bacteria such as *Escherichia coli*, cAMP binds to the catabolite activator protein (CAP), also known as cAMP receptor protein (CRP), enabling it to regulate global catabolic gene expression related to carbon metabolism^20,21^.

Compared to bacterial and human ACs, fungal ACs exhibit a complex multidomain architecture (Fig. 1). Bacterial ACs typically range from 269 to 950 amino acids (aa) in length and contain a single AC catalytic domain. While most bacteria encode only one AC, humans express multiple AC paralogs, each ranging from 1,077 to 1,610 aa and usually containing two catalytic domains. In contrast, fungal ACs are significantly larger, ranging from 1,690 to 2,026 aa, and are composed of multiple distinct domains. These include a Gα-binding (GB) domain, a Ras-associating (RA) domain, leucine-rich repeats (LRRs), a protein phosphatase 2C (PP2C) domain, and adenylate/guanylate cyclase catalytic (ACC) domain (Fig. 1). The presence of GB and RA domains suggests that fungal ACs are regulated by either heterotrimeric G-proteins or Ras-family small GTPases (or both). In some fungi, an additional cyclase-associated protein (CAP)-binding domain is present at the C-terminus, which mediates interaction with CAP proteins. These CAP proteins modulate cAMP induction in response to upstream signals—without affecting basal cAMP levels— and thereby govern most cAMP-dependent phenotypes^8,22^.

**Fig 1.**
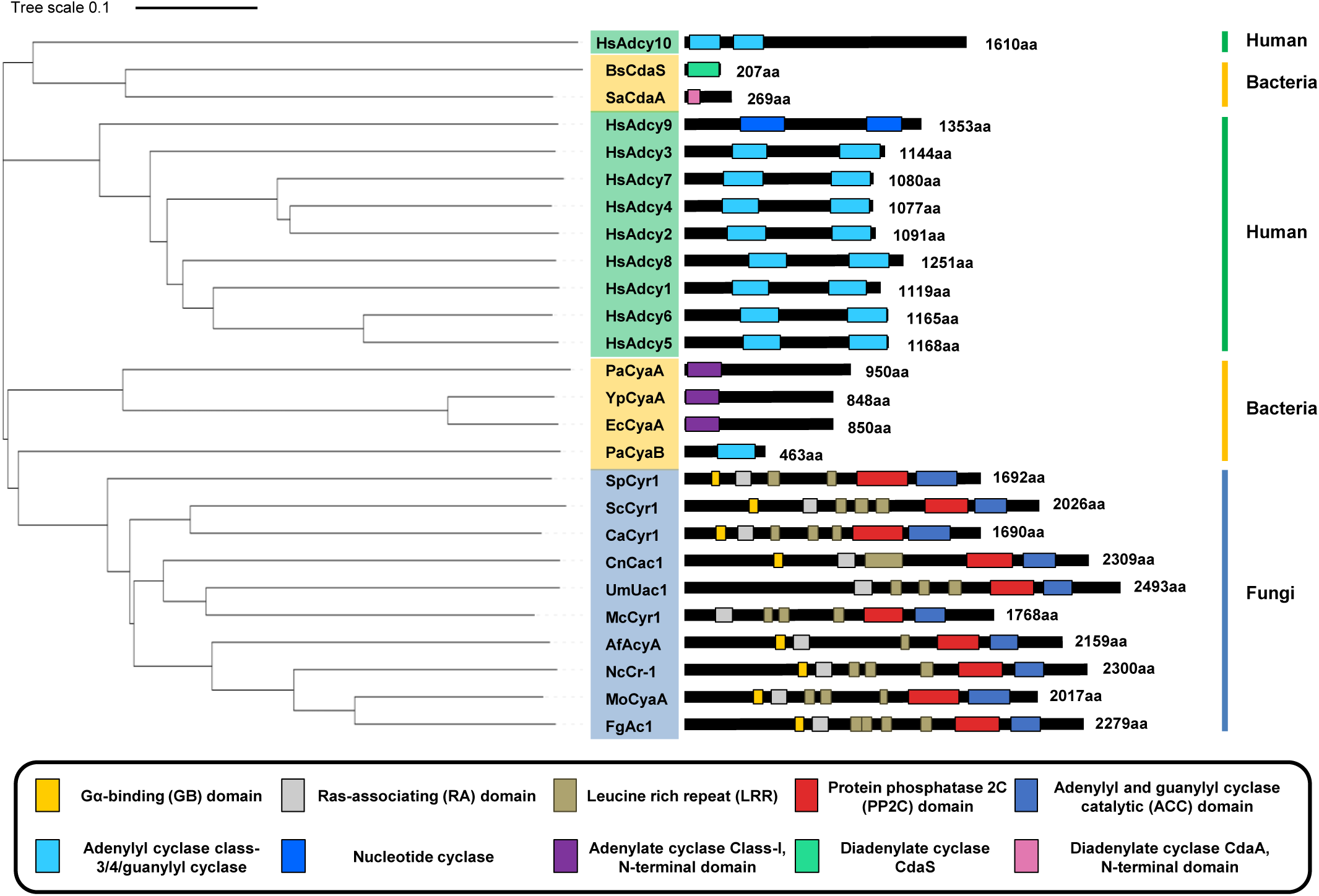
Domain architecture and phylogenetic relationships of Cac1 and orthologous adenylyl cyclases. A phylogenetic tree (left; scale bar = 0.1 substitutions per site) is shown alongside linear domain maps (right) for representative ACs from humans (green), bacteria (orange), and fungi (blue). For each protein, predicted domains are displayed on a schematic of the full-length sequence with the total length (aa) indicated at the right. Species abbreviations: Hs, *Homo sapiens*; BS, *Bacillus subtilis*; SA, *Staphylococcus aureus*; Pa, *Pseudomonas aeruginosa*; Yp, *Yersinia pestis*; Ec, *Escherichia coli*; Sp, *Schizosaccharomyces pombe*; Sc, *Saccharomyces cerevisiae*; Ca, *Candida albicans*; Cn, *Cryptococcus neoformans*; Um, *Ustilago maydis*; Mc, *Malassezia sp*.; Af, *Aspergillus fumigatus*; Nc, *Neurospora crassa*; Mo, *Magnaporthe oryzae*; Fg, *Fusarium graminearum*.

Unlike other AC domains, the unique presence of a PP2C domain in fungal ACs remains enigmatic, as this domain is absent in both bacterial and mammalian ACs. Our bioinformatic analysis indicates that all reported fungal ACs consistently harbour a PP2C domain adjacent to the ACC domain (Fig. 1), suggesting that this domain may serve an evolutionarily conserved role specific to the fungal kingdom. In *C. neoformans*, deletion of the AC gene *CAC1* markedly reduces the production of key virulence factors, including capsule and melanin, impairs sexual differentiation and titan cell formation, and eliminates virulence^7,8,23^. Moreover, in our prior genome-wide functional survey of 114 predicted phosphatases in *C. neoformans*, Cac1 was classified as a putative pathogenicity-related phosphatase due to the presence of its PP2C domain^24^. However, it remains unclear whether this domain possesses bona fide phosphatase activity or contributes directly to the pathobiological function of Cac1.

To address this question, we investigated the in vitro and in vivo roles of the Cac1-PP2C domain in *C. neoformans* using a combination of biochemical and molecular genetic approaches. We demonstrate that the PP2C domain in Cac1 possesses Ser/Thr-specific phosphatase activity and plays a critical role in regulating key pathobiological processes, including melanin and capsule production, sexual differentiation, titan cell development, cell wall integrity, and overall virulence. Notably, transcriptomic and phosphoproteomic analyses revealed that the PP2C domain modulates a broad network of downstream targets, including components of major signalling pathways such as mitogen-activated protein kinases (MAPKs) and calcineurin. Together, these findings provide insight into the function of the evolutionarily conserved PP2C domain in fungal ACs and suggest its potential as a fungal-specific therapeutic target.

## RESULTS

### The noncanonical Cac1-PP2C domain possesses Ser/Thr phosphatase activity

To investigate the structural basis of the PP2C domain function in Cac1, we modelled the *C. neoformans* Cac1 protein using AlphaFold3^25^. The resulting structure included five annotated Cac1 domains (876-2309 aa): RA, LRR, PP2C, ACC, and CAP-binding domains (Fig. 2A). The CAP-binding domain was included because the C-terminal region of *C. neoformans* Cac1 exhibits clear sequence homology to characterised CAP-binding domains. In contrast, the GB domain was excluded from the model due to extensive intrinsic disorder, which impeded reliable structural prediction^26^. The PP2C domain forms the structural core of Cac1 and is encased by a concave surface consisting of 22 tandem LRR domains. The most extensive interface between the LRR and PP2C domains occurs within the ascending loops of LRR4-9 and LRR15-22, as well as the C-terminal CAP through hydrogen bonding and ionic interactions. The RA domain is situated at the N-terminal side of the LRRs, interacting with the α2 helix and flap subdomain of the PP2C domain. In contrast, the ACC and CAP-binding domains are positioned on the β-sheet core of the PP2C domain and the convex surface of the LRR, respectively.

**Fig 2.**
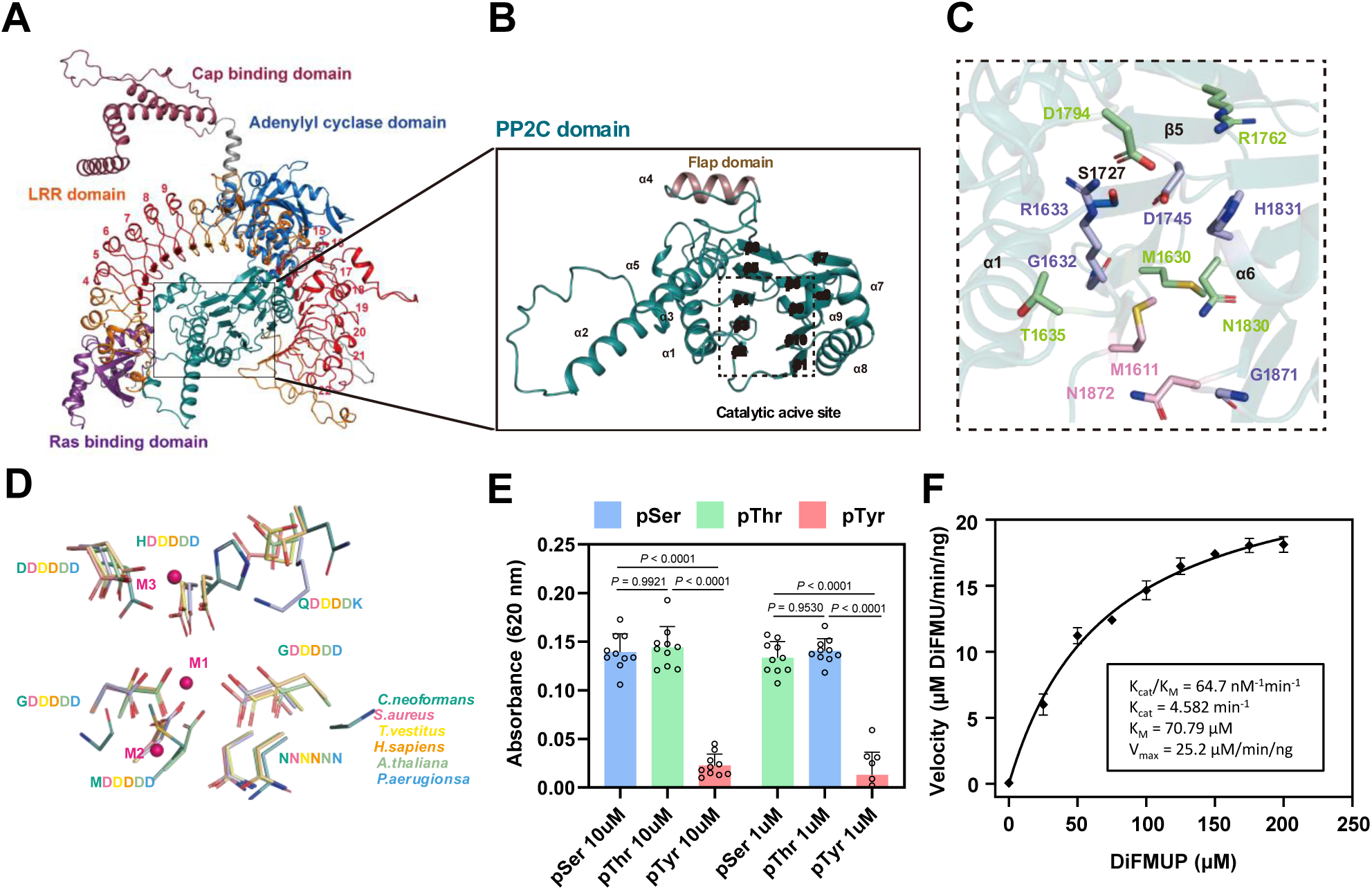
Structural modelling and biochemical characterisation of the PP2C domain of Cac1. (A)Structural model of the Cac1 using AlphaFold3. Each domain is differently coloured (purple: Ras binding domain, orange: LRR domain, green: PP2C domain, blue: adenylyl cyclase domain and maroon: Cap binding domain) and an undefined part is shown in grey. (B) Predicted overall structure of the phosphatase domain PP2C with annotation of secondary structure elements (α1-9, β1-10). The flap domain is coloured brown. The active site in beta sheet core are represented by dashed line boxes. (C) Active site of PP2C domain. Residues are shown as combination of cartoon and stick models. Potential Mn^2+^ (pink) and Mg^2+^ (green) binding sites residues predicted by IONCOM program. Residues capable of binding both metal ions are also represented (blue). (D) Metal binding site alignment of various PP2C like phosphate. The predicted three metal binding M1, M2 and M3 site are shown as sphere (magenta) and residues are shown as sticks (green: *C. neoformans*, light green: *A. thaliana*, yellow: *H. sapiens*, lime: *T. vestitus*, blue: *P. aeruginosa*, salmon: *S. aureus*). (E) Determination of PP2C phosphatase substrate specificity among Phospho-Ser (blue bar), Phosphor-Thr (green bar) and Phosphor-Tyr (pink bar) (10μM and 1μM). Malachite green assays reveal that PP2C phosphatase exhibits activity only on Phosphor-Ser, Phosphor-Thr. Absorbance was measured at a wavelength of 620 nm and shown in bar using Prism 9. Data are means ± SEM (n = 3 independent experiments). (F) Michaelis-Menten plots and kinetic parameters Kcat, Vmax, KM, and Kcat/KM for the hydrolysis of DiFMUP (0-200 μM) by PP2C at 37 °C. Plots were fit by non-linear regression analysis to the Michaelis-Menten model using Prism 9. Kinetic analyses were conducted for PP2C using 40 µg of purified recombinant enzyme. Data are means ± SEM (n = 3 independent experiments).

The Cac1-PP2C domain adopts the typical secondary structure conserved among PP2C phosphatases^27^, consisting of a central β-sandwich formed by two five-stranded antiparallel β-sheets flanked by two pairs of antiparallel α helices (Fig. 2B). To identify structural deviations, we aligned the Cac1-PP2C structures from multiple species, including *Arabidopsis thaliana* PP2C 77^28^ (PDB code: 3UJK), cyanobacterial *Thermosynechococcus vestitus* PP2C^29^ (PDB code: 5ItI), *Pseudomonas aeruginosa* PppA^30^ (PDB code: 6JKV), *Staphylococcus aureus*^31^ (PDB code: 5F1M), and human PP2Ca^32^ (PDB code: 4RAF) (Fig. S1A, 1B). The Cac1-PP2C domain, like its bacterial counterparts, lacks the three additional C-terminal α-helices characteristic of human PP2C^33^ (Fig. S1A). Moreover, the conformation of the flap subdomain in Cac1 differs from that of bacterial proteins^34^, while more closely resembling the structures found in human and *Arabidopsis* PP2C^35^.

Canonical PP2C phosphatases contain highly conserved Asp residues that coordinate metal ions essential for catalysis^28–32,36–38^. In contrast, the predicted active site of Cac1-PP2C shows substantial divergence in residue composition. At the M1 and M2 positions, the conserved Asp residues are replaced by G1632 and M1611, respectively, when aligned by α-carbon position. By contrast, at the M3 site, only one of the three canonical Asp residues is retained, with the remaining two substituted by H1831 and Q1835 (Fig. 2C, 2D). Because of the atypical active-site composition, we used IONCOM^39,40^ to predict potential metal-binding residues (Fig. 2C). Several candidates including G1632, R1633, D1745, H1831, and G1871 were predicted to bind both Mg² and Mn² and clustered within the PP2C active site. Together, these findings indicate that the Cac1 PP2C domain possesses a noncanonical structural scaffold distinct from other PP2C phosphatases.

To evaluate whether the noncanonical Cac1-PP2C domain retains phosphatase activity, we expressed and purified the recombinant protein with Mn² supplementation (Fig. S1B and S1C). We next assessed the phosphatase activity of the Cac1-PP2C domain using a malachite green phosphate assay^41^ with phosphorylated Ser, Thr, and Tyr substrates (Fig. 2E). At substrate concentrations of 10 μM and 1 μM, the Cac1-PP2C domain exhibited robust dephosphorylation of p-Ser and p-Thr but showed no activity toward p-Tyr, indicating that it functions as a Ser/Thr-specific phosphatase. Kinetic analysis using 6,8-difluoro-4-methylumbelliferyl phosphate^42^ (DiFMUP) revealed a Michaelis constant (Km) of 65.41 μM for the Cac1-PP2C domain (Fig. 2F), which is notably lower than the millimolar-range of Km values previously reported for human and bacterial PP2Cs using p-nitrophenyl phosphate (pNPP) as a substrate^43^. Taken together, these findings demonstrate that the Cac1 PP2C domain possesses a noncanonical structural scaffold distinct from other PP2C phosphatases, yet still functions as a metalloenzyme with Ser/Thr phosphatase activity in *C. neoformans*.

### The PP2C domain is required for Cac1 function in melanin biosynthesis

To elucidate the pathobiological role of the Cac1-PP2C domain, we constructed domain-specific deletion strains in *C. neoformans* by complementing a *cac1*Δ mutant with the *CAC1^PP2C^*^Δ^ or *CAC1^ACC^*^Δ^ allele, in which the PP2C domain (aa 1611-1874) or the ACC domain (aa 1934-2119) was selectively deleted and reintegrated into the native locus. This approach generated the *cac1*Δ::*CAC1^PP2C^*^Δ^ strain (hereafter the *CAC1^PP2C^*^Δ^ strain) and the *cac1*Δ::*CAC1^ACC^*^Δ^ strain (hereafter the *CAC1^ACC^*^Δ^ strain) (Fig. 3A). As a control, the *cac1*Δ mutant was complemented with the full-length wild-type *CAC1* allele (hereafter the *CAC1* strain). Each allele was verified by sequencing prior to integration at the native *CAC1* locus, and targeted insertion was confirmed by diagnostic PCR (Fig. S2B). *CAC1* transcript levels were comparable across the wild-type, *CAC1*, *CAC1^PP2C^*^Δ^, and *CAC1^ACC^*^Δ^ strains (Fig. S2C), and AlphaFold3-predicted structures of the truncated Cac1*^PP2C^*^Δ^ and Cac1*^ACC^*^Δ^ proteins retained the overall architecture of full-length Cac1 (Fig. S2D).

**Fig 3.**
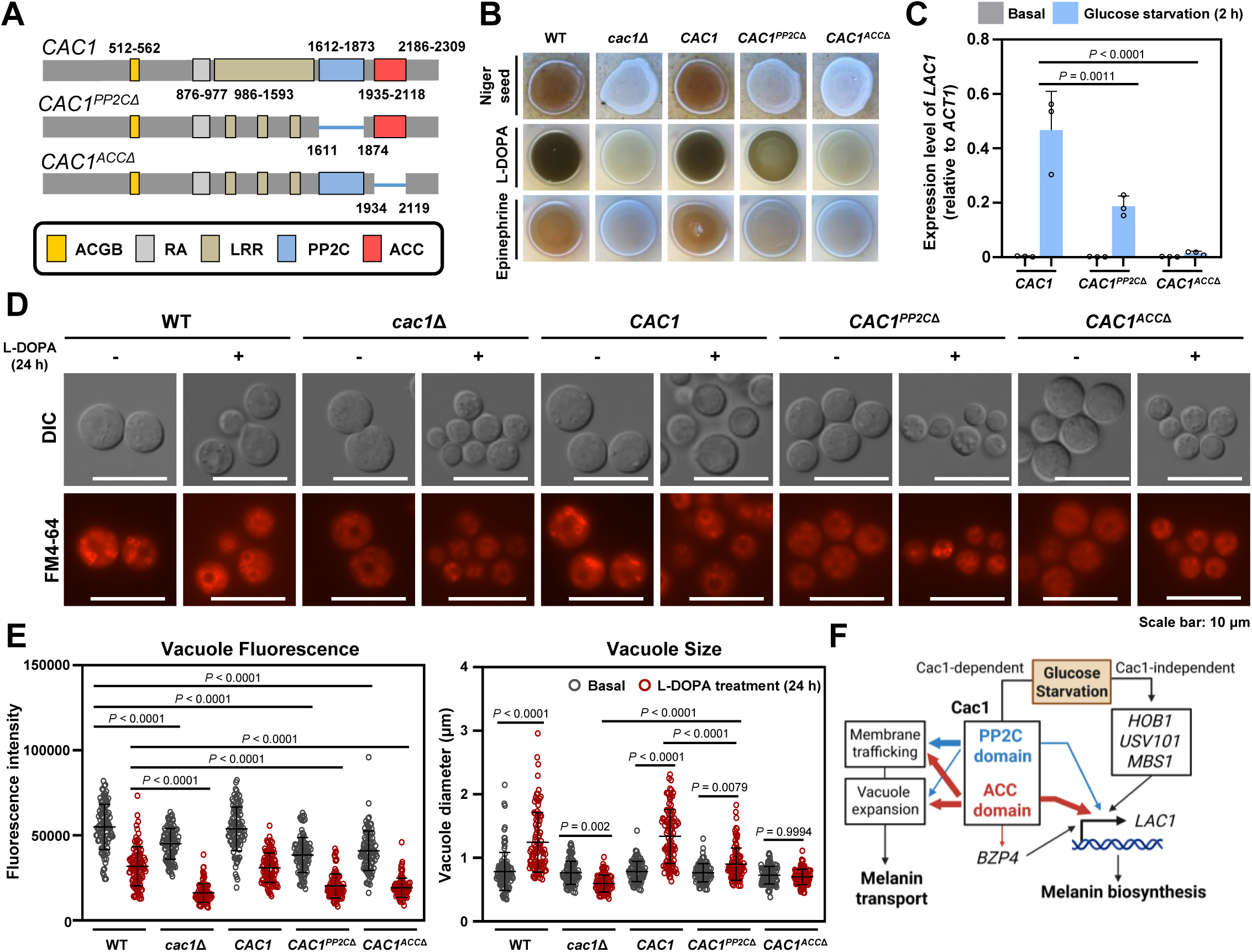
The PP2C domain is required for Cac1 function in melanin biosynthesis in *C. neoformans*. (A) Constructs used to generate *CAC1* strain consisting of the full-length wild-type *CAC1* allele, *CAC1^PP2C^*^Δ^ strain lacking the PP2C domain, and *CAC1^ACC^*^Δ^ strain lacking the ACC domain. Each allele was inserted at the native *CAC1* locus. (B) Melanisation assay. The indicated strains were spotted on Niger seed, L-DOPA, or epinephrine media under standard conditions. Representative colony pigmentation is shown. (C) *LAC1* transcript levels measured by qRT-PCR. Basal conditions are shown in grey, and glucose starvation is shown in blue. Three biological replicates were performed with three technical replicates each, and the expression level of each gene was normalised to *ACT1* (ΔCt). Error bars indicate SEM. Statistics were performed by one-way ANOVA with Tukey’s multiple comparisons. (D) Vacuole staining using FM4-64. Representative DIC and FM4-64 fluorescence images of cells grown in basal medium or after 24 h in L-DOPA. Scale bar, 10 μm. (E) Quantification of vacuolar phenotypes. Whole-cell FM4-64 fluorescence intensity was measured for 100 randomly selected cells, and vacuole diameter was measured for 100 vacuoles. Basal conditions are shown in grey, and L-DOPA treated conditions are shown in red. Error bars indicate SEM. Same statistics analysis was applied. (F) Proposed model for the Cac1-PP2C and Cac1-ACC domains in melanin biosynthesis under glucose starvation in *C. neoformans*. Both the PP2C (blue) and ACC (red) domains drive *LAC1* induction, with the PP2C domain contributing less than the ACC domain. Both domains additionally support membrane trafficking and vacuolar expansion required for melanin transport, although the PP2C domain plays a supportive role in vacuolar expansion. Arrow thickness indicates the relative magnitude of contribution. Created with BioRender.com (https://BioRender.com/nlretcs).

To further ensure that any phenotype observed in the domain-deletion strains reflects domain-specific functional loss rather than altered Cac1 stability, we generated strains expressing C-terminally 4xFLAG-tagged Cac1, Cac1*^PP2C^*^Δ^, and Cac1*^ACC^*^Δ^ (Fig. S2E), with correct integration confirmed by diagnostic PCR (Fig. S2F). FLAG tagging did not interfere with Cac1 function, as the tagged strains recapitulated the melanin phenotypes of their untagged counterparts (Fig. S2G). Western blotting confirmed that all three Cac1 variants – Cac1-4xFLAG (257 kDa), Cac1*^PP2C^*^Δ^-4xFLAG (227 kDa), and Cac1*^ACC^*^Δ^-4xFLAG (236 kDa) – were expressed at comparable levels and at their predicted molecular weights under basal conditions (Fig. S2H). Protein levels also remained stable under glucose starvation and cell wall stress (Fig. S2I). Together with the comparable transcript levels across strains, these results indicate that neither domain deletion compromises Cac1 protein abundance or stability, supporting that the phenotypes reported in this study reflect domain-specific functional contributions rather than differences in Cac1 abundance.

Cac1 governs the biosynthesis and intracellular trafficking of the polyphenol melanin pigment, which confers antioxidant and antiphagocytic properties and is essential for cryptococcal virulence^44,45^. As previously reported^7,24^, the *cac1*Δ mutant failed to produce melanin under various inducing conditions, including Niger seed, L-DOPA, and epinephrine media (Fig. 3B). While the *CAC1* strain restored melanin production to wild-type levels, the *CAC1^ACC^*^Δ^ strain failed to do so (Fig. 3C), confirming the essential role of the ACC domain. Notably, the *CAC1^PP2C^*^Δ^ strain partially rescued melanin synthesis (Fig. 3B), suggesting that the PP2C domain contributes to—but is not solely responsible for—this phenotype.

Melanin synthesis requires transcriptional induction of *LAC1*, which encodes the laccase, the key enzyme for eumelanin biosynthesis in *C. neoformans*. Consistent with prior findings^24^, *LAC1* expression was rapidly induced under carbon starvation in the wild-type and *CAC1* strains but was abolished in the *CAC1^ACC^*^Δ^ strain (Fig. 3C). Notably, *LAC1* expression was moderately but significantly reduced in the *CAC1^PP2C^*^Δ^ strain compared to the *CAC1* strain (*P* = 0.001; Fig. 3C), indicating that the PP2C domain is required for full induction of *LAC1* under carbon-limited conditions. In contrast, the expression levels of key melanin-regulating transcription factors that control *LAC1* induction—*HOB1*, *BZP4*, *USV101*, and *MBS1*^45^—were not significantly altered in the *CAC1^PP2C^*^Δ^ strain (*P* > 0.05 for all; Fig. S3). By comparison, *BZP4* induction alone was decreased in the *CAC1^ACC^*^Δ^ strain (*P* = 0.001), suggesting that the ACC domain is required for full *BZP4* induction (Fig. S3).

Melanin formation in *C. neoformans* depends on vesicle trafficking and vacuolar function^45,46^. Therefore, to examine vacuolar morphology, we used FM4-64, a lipophilic styryl dye that intercalates into the plasma membrane and subsequently labels the vacuolar membrane^47,48^. Cells were imaged at baseline and after 24 h of incubation in minimal medium supplemented with L-DOPA to induce melanisation. In the wild-type and *CAC1* strains, vacuoles appeared small, round, and distinctly labelled at baseline and became enlarged following L-DOPA treatment (Fig. 3D). By contrast, the *cac1*Δ mutant exhibited markedly reduced FM4-64 fluorescence under both basal and L-DOPA conditions compared to the wild-type (*P* < 0.0001 for both conditions). Furthermore, rather than expanding as observed in the wild-type, the vacuole diameter of the *cac1*Δ mutant was modestly reduced after L-DOPA treatment (*P* = 0.002 versus basal), indicating severe defects in vacuole formation and the structural remodelling required for melanisation (Fig. 3D, 3E). Like the *cac1*Δ mutant, the *CAC1^ACC^*^Δ^ strain exhibited markedly reduced FM4-64 fluorescence relative to the wild-type (*P* < 0.0001) and failed to undergo vacuolar expansion after L-DOPA treatment (*P* > 0.05 versus basal; Fig. 3D, 3E). The *CAC1^PP2C^*^Δ^ strain also shared this severe reduction in FM4-64 fluorescence (*P* < 0.0001 versus wild-type) but notably retained a partial capacity for vacuolar expansion, resulting in an intermediate post-treatment size that was significantly smaller than the wild-type (*P* < 0.0001) but still larger than the *cac1*Δ mutant (*P* < 0.0001) (Fig. 3D, 3E). Given that FM4-64 fluorescence is consistent with the dynamic flux of membrane and vesicle fusion events required to maintain vacuolar homeostasis^47,49^, while the vacuolar lumen expansion is consistent with accommodating elevated secretory demand under melanin-inducing conditions, these results indicate that the PP2C domain is required for maintaining steady-state membrane trafficking and contributes to vacuolar remodelling. Together with the reduced *LAC1* induction observed in the *CAC1^PP2C^*^Δ^ strain, these findings demonstrate that the Cac1-PP2C domain supports optimal melanin biosynthesis by coordinating both transcriptional regulation and vacuole-mediated secretory processes required for laccase delivery to the cell surface in *C. neoformans* (Fig. 3F).

### The PP2C domain is required for Cac1 function in serum-mediated capsule biosynthesis

Cac1 also plays a central role in the biosynthesis of capsule, another major virulence factor in *C. neoformans*^7^. The cryptococcal capsule, mainly composed of glucuronoxylomannan (GXM) and galactoxylomannan (GalXM), also contributes to virulence by providing antiphagocytic protection^50,51^. As previously reported^24^, the *cac1*Δ mutant exhibited severe defects in capsule production in Dulbecco’s Modified Eagle (DME), Littman’s (LIT), and foetal bovine serum (FBS) media, compared to the wild-type and *CAC1* strains (*P* < 0.0001 for all conditions; Fig. 4A). In both DME and LIT media, the *CAC1^PP2C^*^Δ^ strain exhibited capsule levels comparable to those of the wild-type and *CAC1* strain (*P* > 0.05 for all; Fig. 4A), suggesting that the PP2C domain is dispensable for capsule formation under these conditions. However, in FBS medium, the *CAC1^PP2C^*^Δ^ strain showed significant defects in capsule synthesis compared to the wild-type (*P* < 0.0001), although less severe than those of *CAC1^ACC^*^Δ^ (Fig. 4A), indicating that the PP2C domain contributes to serum-mediated capsule induction. This defect was not attributable to serum sensitivity, as both *cac1*Δ and *CAC1^PP2C^*^Δ^ strains were as serum-resistant as the wild type, whereas the *ena1*Δ strain, lacking the Ena1 P-type cation ATPase and previously reported to be serum-sensitive^52^, exhibited impaired growth (Fig. S4A, B).

**Fig 4.**
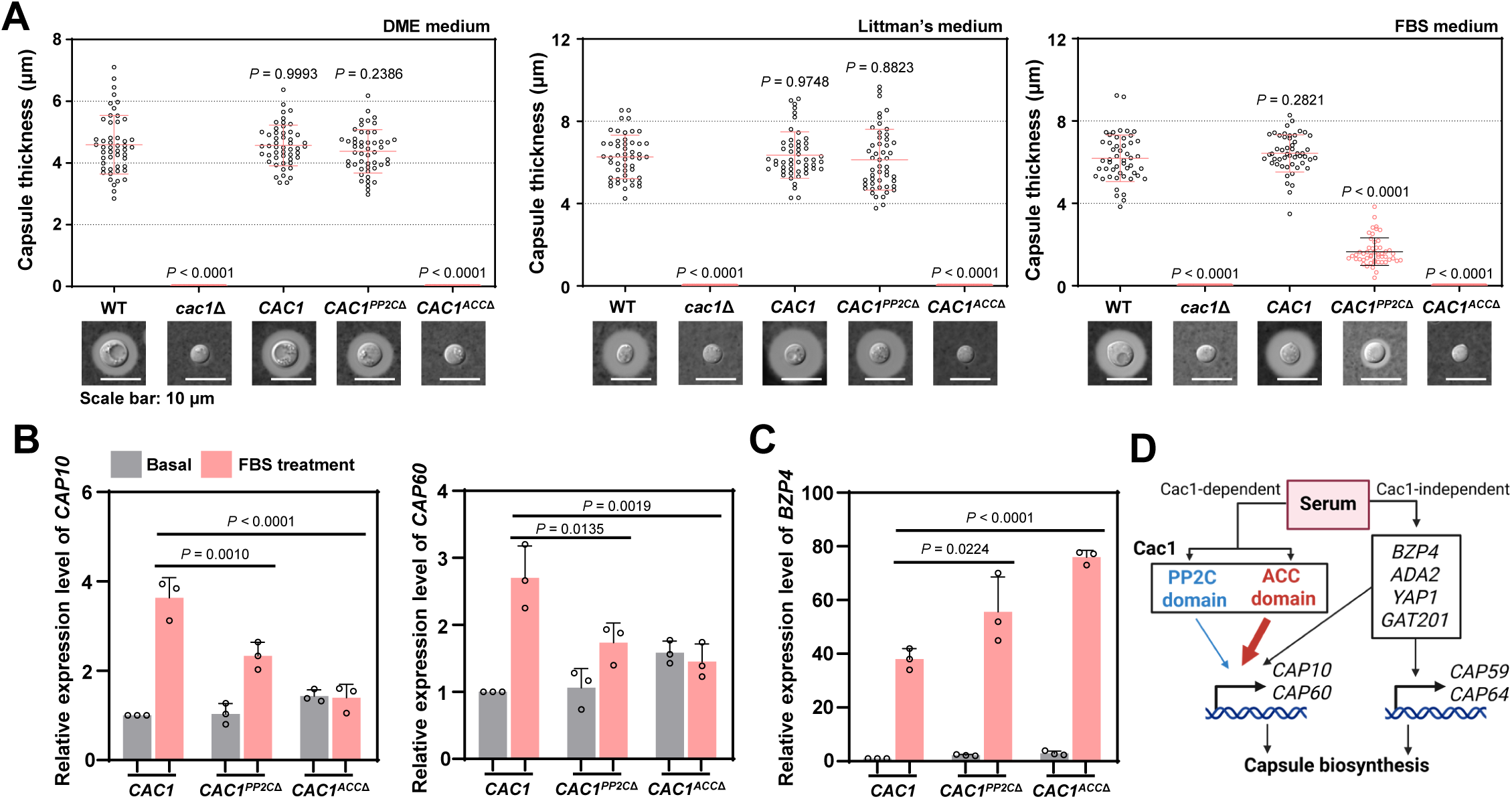
The PP2C domain is required for Cac1 function in serum-mediated capsule biosynthesis in *C. neoformans*. (A) Capsule analysis. Cells were spotted onto DME, LIT, or FBS media, and capsule thickness was measured after India-ink staining (n = 50 cells per condition). Representative images and quantification are shown. Scale bar, 10 μm. Error bars indicate SEM. Statistical significance was determined by one-way ANOVA with Dunnett’s multiple comparisons test*; P*-values denote comparisons between the wild-type and each strain. (B) qRT-PCR of *CAP10* and *CAP60* after 6h induction in 10% FBS. Basal conditions are shown in grey, and FBS treated conditions are shown in pink. Three biological replicates were performed with three technical replicates each, and relative transcript levels were calculated using the 2^−ΔΔCt^ method. Error bars indicate SEM. Statistical significance was determined by one-way ANOVA with Tukey’s multiple comparisons test. (C) qRT-PCR of *BZP4* after 2 h treatment in FBS. Error bars indicate SEM. Same statistics analysis was applied as panel B. (D) Proposed model for the Cac1-PP2C and Cac1-ACC domains in capsule biosynthesis under serum induction in *C. neoformans*. Both the PP2C (blue) and ACC (red) domains drive *CAP10* and *CAP60* induction, with the PP2C domain contributing less than the ACC domain. Cac1-independent transcription factors (*BZP4*, *ADA2*, *YAP1*, *GAT201*) regulate *CAP10, CAP59, CAP60,* and *CAP64* induction in parallel. Arrow thickness indicates the relative magnitude of contribution. Created with BioRender.com. (https://BioRender.com/nlretcs).

Given that *CAP59, CAP10*, *CAP60*, and *CAP64* are each essential for capsule production in *C. neoformans*^53–56^, we measured their expression during serum-mediated capsule induction. Following FBS treatment, both *CAC1^PP2C^*^Δ^ and *CAC1^ACC^*^Δ^ strains showed reduced *CAP10* and *CAP60* expression relative to the *CAC1* strain (*P* < 0.05 for all; Fig. 4B), whereas *CAP59* and *CAP64* levels remained comparable to the complemented strain (*P* > 0.05 for all; Fig. S4C). We next examined capsule-associated regulators (*BZP4, ADA2, GAT201, YAP1*)^57^. Notably, following FBS treatment, *BZP4* expression was induced to a significantly higher level in both domain deletion mutants compared to the *CAC1* strain (*P* < 0.05 for both; Fig. 4C), while *ADA2*, *GAT201*, and *YAP1* remained comparable to the complemented strain (*P* > 0.05 for all; Fig. S4D). These findings suggest that *BZP4* induction is largely Cac1-independent and that serum-specific capsule defects in *CAC1^PP2C^*^Δ^ and *CAC1^ACC^*^Δ^ strains may trigger compensatory upregulation of *BZP4*. Together, these results demonstrate that the Cac1-PP2C domain supports serum-responsive capsule biosynthesis in *C. neoformans* (Fig. 4D).

### The PP2C domain is required for Cac1 function in developmental processes

The key developmental processes of *C. neoformans* diverge between environmental niches and host tissues, with sexual reproduction predominating in the environment and titan cell formation prevailing within the host (Fig. 5A). Given that the cAMP signalling pathway plays a crucial role in orchestrating both processes^58,59^, we next asked whether the Cac1-PP2C domain contributes to these functions. As previously reported^7^, the *MAT*α *cac1*Δ mutant displayed severely impaired unilateral mating when crossed with *MAT***a** wild-type strain (KN99**a**), accompanied by a drastic reduction in basidium and basidiospore formation, although both structures were still observed (Fig. 5B). While the *CAC1* strain fully restored mating efficiency, the *CAC1^ACC^*^Δ^ strain failed to recover the phenotype (Fig. 5B), consistent with the essential role of the ACC domain. Notably, the *CAC1^PP2C^*^Δ^ strain showed partially restored mating efficiency (Fig. 5B), suggesting that the Cac1-PP2C domain contributes to the role of Cac1 in sexual reproduction.

**Fig 5.**
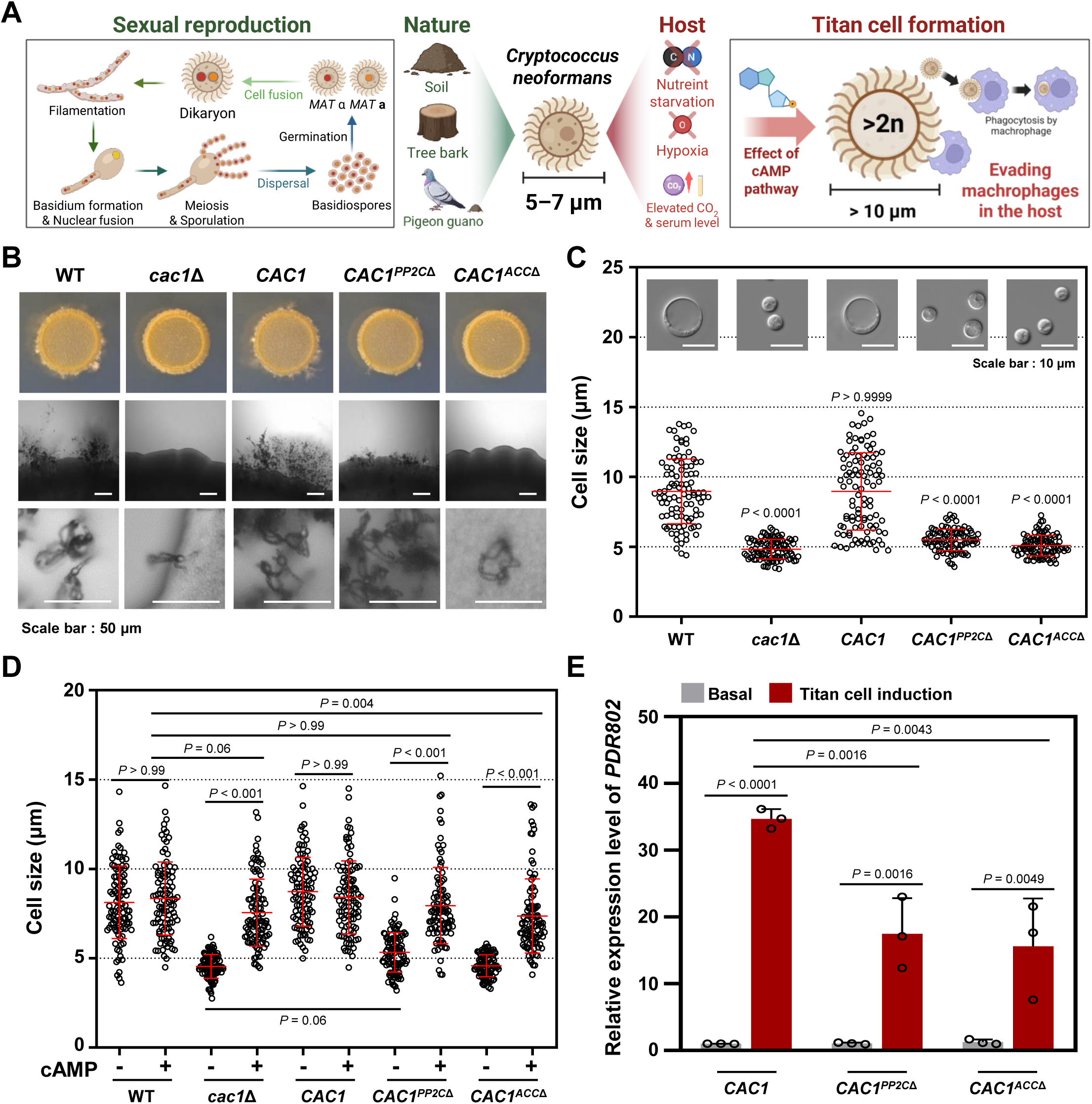
The PP2C domain is required for Cac1-dependent developmental processes in *C. neoformans*. (A) Schematic overview of niche-specific developmental processes in *C. neoformans*. Created with BioRender.com (https://BioRender.com/nlretcs). (B) Unilateral mating assay. Indicated strains were crossed with KN99**a** on MS medium and imaged by bright-field microscopy to visualise cells, filaments, and spores. Scale bar, 50 μm. (C) Titan cell formation in vitro. Cells were incubated under titan-inducing conditions and cell diameters were quantified (n = 100 cells per strain). Representative DIC images are shown above; scale bar, 10 μm. Error bars indicate SEM. Statistics were performed by one-way ANOVA with Dunnett’s multiple comparisons test. (D) Titan cell analysis with cAMP supplementation (10 mM) under the same conditions as in panel C. Error bars indicate SEM. Statistics were performed by one-way ANOVA with Sidak’s multiple comparisons test. (E) qRT-PCR of *PDR802* after titan cell induction. Basal conditions are shown in grey, and titan cell induction conditions are shown in red. Three biological replicates were performed with three technical replicates each, and relative transcript levels were calculated using the 2^−ΔΔCt^ method. Error bars indicate SEM. Error bars indicate SEM. Statistics were performed by one-way ANOVA with Tukey’s multiple comparisons test.

Titan cells are an enlarged morphological form characterised by a thickened cell wall, modified capsule, and increased ploidy, commonly observed during host infection^60^. These cells enhance resistance to phagocytosis by alveolar macrophages and support persistent infection^61^. To investigate whether the PP2C domain is required for titan cell formation, we induced titan cells under established in vitro conditions^62^. As previously shown, *CAC1* deletion completely abolished titan cell formation (*P* < 0.001 compared to the wild-type; Fig. 5C), reaffirming the essential role of the cAMP pathway in this process. The *CAC1* strain restored titan cell formation (*P* > 0.99), whereas the *CAC1^ACC^*^Δ^ strain did not (*P* < 0.001; Fig. 5C), indicating that ACC activity is indispensable. Importantly, the *CAC1^PP2C^*^Δ^ strain also failed to restore titan cell formation (*P* < 0.001; Fig. 5C), suggesting that the PP2C domain is critical for this developmental transition. As expected, exogenous cAMP supplementation restored titan cell formation to the wild-type level in the *cac1*Δ strain (*P* = 0.06; Fig. 5D). Strikingly, the *CAC1^PP2C^*^Δ^ strain was also fully rescued by exogenous cAMP (*P* > 0.99 versus the wild type; Fig. 5D), demonstrating that the titan cell defect caused by loss of the PP2C domain can be bypassed by restoring cAMP signalling. The *CAC1^ACC^*^Δ^ strain was also partially rescued, although titan cell formation in this strain remained significantly below wild-type levels (*P* = 0.004; Fig. 5D). Together, these data demonstrate that both domains operate within the cAMP signalling cascade to regulate this process.

The GPCR Gpr5 and the transcription factor Rim101 are key signalling components of the cAMP pathway that are essential for titan cell formation^23,62^. We confirmed that both *GPR5* and *RIM101* were induced under titan cell-inducing conditions (*P* = 0.002 and *P* < 0.001, respectively; Fig. S5A); however, their expression was unaffected by deletion of either the ACC or PP2C domains of Cac1 (*P* > 0.05 for all; Fig. S5A), suggesting that these domains are not required for the transcriptional activation of these signalling components. Pdr802 is another transcription factor involved in titan cell regulation; its deletion leads to dramatic titan cell expansion ^63^. We found that *PDR802* expression was significantly reduced under titan-inducing conditions in both the *CAC1^PP2C^*^Δ^ and *CAC1^ACC^*^Δ^ strains compared to the *CAC1* control strain (*P* = 0.001 and *P* < 0.001, respectively; Fig. 5E). To determine whether this downregulation reflects impaired cAMP signalling, we supplemented the titan-inducing medium with exogenous cAMP and measured *PDR802* expression. Notably, *PDR802* expression in both the *CAC1^PP2C^*^Δ^ and *CAC1^ACC^*^Δ^ strains after titan cell induction was restored to levels comparable to the complemented strain (*P* > 0.05 for both; Fig. S5B), indicating that *PDR802* expression is positively coupled to cAMP signalling and that its downregulation in the domain-deletion strains is a direct consequence of reduced cAMP signal output. Because Pdr802 acts as a negative regulator of titan cell formation, its reduced expression is not expected to drive the titan cell defect of the domain-deletion strains; rather it may partially counteract the compromised cAMP signal by relieving Pdr802-mediated repression, although the net titan cell phenotypes remain dominated by the upstream loss of cAMP signalling.

Together, these findings demonstrate that the Cac1-PP2C domain contributes to sexual reproduction and is essential for titan cell formation, two major developmental processes that enable *C. neoformans* to survive and proliferate in environmental and host niches, respectively.

### The role of the Cac1-PP2C domain in the virulence of *C. neoformans*

Given the critical role of the Cac1-PP2C domain in melanin, capsule, and titan cell formation, we next investigated its contribution to virulence using a murine model of systemic cryptococcosis. Mice were infected through intranasal inhalation with wild-type, *cac1*Δ, *CAC1*, *CAC1^PP2C^*^Δ^, and *CAC1^ACC^*^Δ^ strains. As previously reported^7^, mice infected with the *cac1*Δ mutant survived throughout the experimental period, while those infected with the wild-type or *CAC1* strain became moribund between 16 and 23 days post-infection (dpi), accompanied by significant weight loss (Fig. 6A, B). Consistent with the essential role of the ACC domain, the *CAC1^ACC^*^Δ^ strain was also completely avirulent (Fig. 6A). The *CAC1^PP2C^*^Δ^ strain also exhibited a similar avirulent phenotype (Fig. 6A), and no significant weight loss was observed during infection (Fig. 6B). When monitored over an extended period, all mice infected with the *CAC1^PP2C^*^Δ^ strain survived for at least 111 dpi (Fig. S6A).

**Fig 6.**
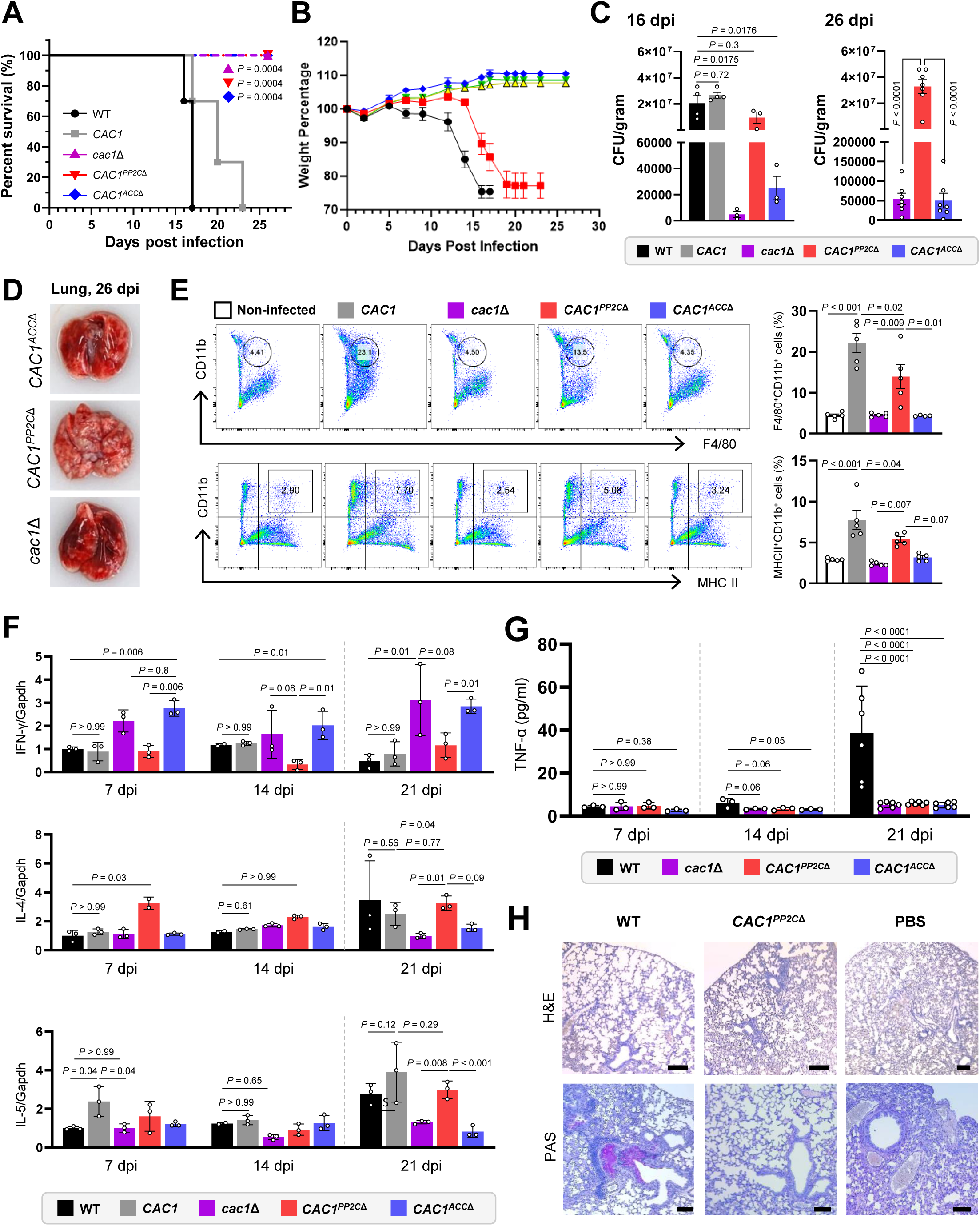
Survival analysis of *CAC1* mutants in a murine cryptococcosis model. BALB/c mice were infected intranasally with 5 × 10^5^ cells of *Cryptococcus neoformans* strains and sacrificed at indicated time points for each analysis. (A-B) Survival assay (n=7). (A) Survival curve and (B) body weight changes of infected mice with log-rank (Mantel–Cox) test, with corresponding *P*-values: WT vs +*CAC1* complemented strain (>0.9999); WT vs *cac1*Δ (0.0004); WT vs +*CAC1^PP2C^*^Δ^ (0.0004); WT vs +*CAC1^ACC^*^Δ^ (0.0004). (C) Fungal burden (colony-forming units) in lung tissue at days 16 and 26 post-infection. (D) Representative lung morphology of *CAC1* mutants at day 26 post-infection. (E) Flow cytometry analysis of pulmonary immune cells (n=5) 14 dpi. Macrophages and dendritic cells were identified by staining with anti-F4/80, anti-MHC II, and anti-CD11b antibodies. (F) Gene expression profiles in lung tissue at 7, 14, and 21 days post-infection (n=3). (G) Serum TNF-α levels in mice infected with *C. neoformans* strains. (H) Histopathological analysis of lung tissues at day 16 post-infection. Sections were stained with H&E (top panels) to evaluate tissue architecture and inflammation, and with PAS (bottom panels) to visualise mucus production and polysaccharide deposition. Representative images are shown; scale bars = 100 μm. Statistical analysis (panels C, E, F, and G) was performed using one-way ANOVA followed by Tukey’s multiple-comparison test. Data are displayed as mean ± SEM with individual data points plotted. Significance levels: *P* < 0.05 (*), *P* < 0.01 (**), *P* < 0.001 (***), *P* < 0.0001 (****); ns, not significant.

Despite their similar impact on host survival, the *CAC1^ACC^*^Δ^ and *CAC1^PP2C^*^Δ^ strains differed markedly in fungal burden. At 16 and 26 dpi, mice infected with the *CAC1^PP2C^*^Δ^ strain harboured fungal loads in the lungs comparable to those infected with the wild-type or *CAC1* strain, whereas mice infected with the *cac1*Δ or *CAC1^ACC^*^Δ^ strain had markedly reduced pulmonary fungal burdens (*P* = 0.0175 and *P* = 0.0176, respectively; Fig. 6C). However, by 111 dpi, fungal clearance had occurred in the *CAC1^PP2C^*^Δ^-infected mice, as very few fungal cells were recovered from either the lung or brain (Fig. S6B), suggesting an eventual resolution of infection.

Interestingly, histological examination at 26 dpi revealed multifocal pale inflammatory nodules and uneven alveolar collapse in the lungs of mice infected with the *CAC1^PP2C^*^Δ^ strain (Fig. 6D and Fig. S6E), suggesting a localised inflammatory response, although this pathological change did not lead to mortality. To further investigate the host immune response, we analysed immune cell populations and cytokine expression in lung tissues. At 14 dpi, *CAC1^PP2C^*^Δ^-infected mice exhibited elevated infiltration of F4/80^+^ CD11b^+^ macrophages and MHCII^+^ CD11b^+^ antigen-presenting cells, representing an intermediate phenotype between the *CAC1* strain and the non-infected, *cac1*Δ, and *CAC1^ACC^*^Δ^ strains (Fig. 6E).

Cytokine profiling revealed divergent immune polarisation among strains. At 7, 14, and 21 dpi, *cac1*Δ or *CAC1^ACC^*^Δ^ infections were associated with elevated pulmonary IFNγ levels—a hallmark Th1 response—relative to wild-type*, CAC1*, and *CAC1^PP2C^*^Δ^ strains (Fig. 6F). In contrast, IL-4 and IL-5—cytokines associated with Th2 responses—were elevated and appeared earlier in *CAC1^PP2C^*^Δ^-infected mice than in wild-type and *CAC1-*infected mice. Meanwhile, *cac1*Δ or *CAC1^ACC^*^Δ^ strains elicited minimal IL-4 and IL-5 expression (Fig. 6F). Further supporting a Th2-skewed pathology, lung tissue from *CAC1^PP2C^*^Δ^-infected mice exhibited progressive and persistent emphysema-like pulmonary destruction, evident as parenchymal disorganisation at 16 dpi (Fig. 6H) and residual architectural damage in long-term survivors at 111 dpi (Fig. S6C, D). However, systemic immune activation appeared limited; TNF-α levels in serum remained low in *CAC1^PP2C^*^Δ^-infected mice compared to those infected with the wild-type strain (Fig. 6G).

In summary, both the ACC and PP2C domains of Cac1 are required for full virulence of *C. neoformans*. Despite this shared loss of virulence, deletion of the PP2C domain results in distinct immunological outcomes, including a heightened Th2 immune response, emphysema-like pathology, and eventual fungal clearance without systemic dissemination or lethality.

### The Cac1-PP2C domain is required for cell wall remodelling in *C. neoformans*

Given that deletion of either the ACC or PP2C domain alters immune responses and cytokine profiles during infection, we hypothesise that impairment of the cAMP pathway could affect cell surface architecture and antigen exposure in *C. neoformans*. The cryptococcal cell surface structure consists of inner and outer cell wall layers as well as exopolysaccharide capsules. The inner wall contains chitin and chitosan adjacent to the plasma membrane and β-glucans, while the outer wall is composed primarily of α-glucans and β-glucans. The capsule is anchored to the outer wall layer^64^. To investigate whether the *cac1*Δ mutation or domain-specific deletions could alter cell wall composition, we used a panel of fluorescent stains—calcofluor white (CFW), Eosin Y (EY), FITC-conjugated wheat germ agglutinin (WGA), Concanavalin A (ConA), and Dectin-1—to label distinct cell wall components, including chitin (CFW, WGA), chitosan (Eosin Y), mannoproteins (ConA), and β-glucan (Dectin-1).

We found that *CAC1* deletion markedly reduced CFW and EY staining (*P* < 0.0001 for both; Fig. 7A and 7B), indicating a significant decrease in chitin and chitosan deposition at the cell surface. Notably, both *CAC1^ACC^*^Δ^ and *CAC1^PP2C^*^Δ^ strains exhibited similar reductions compared to the wild-type (*P* < 0.0001 for all; Fig. 7A and 7B), suggesting that both the ACC and PP2C domains are required for proper chitin and chitosan incorporation. Furthermore, ConA staining was significantly diminished in the *cac1*Δ, *CAC1^PP2C^*^Δ^, and *CAC1^ACC^*^Δ^ strains relative to wild-type and *CAC1* strains (*P* < 0.0001 for all; Fig. 7A and 7B), indicating that Cac1 also influences mannan and mannoprotein levels. In contrast, WGA staining was not significantly altered among the strains (*P* > 0.05 for all; Fig. 7A and 7B), suggesting that the levels of surface-exposed chito-oligomer remain unaffected by Cac1 or its domains. Similarly, staining with the soluble human Dectin-1 receptor revealed that β-1,3-glucans remained masked and undetectable in the wild-type as well as in all mutant strains, whereas the *mpk1*Δ mutant, which lacks the gene encoding the core MAPK of the cell wall integrity pathway, showed positive staining as previously reported^65^ (Fig. S7).

**Fig 7.**
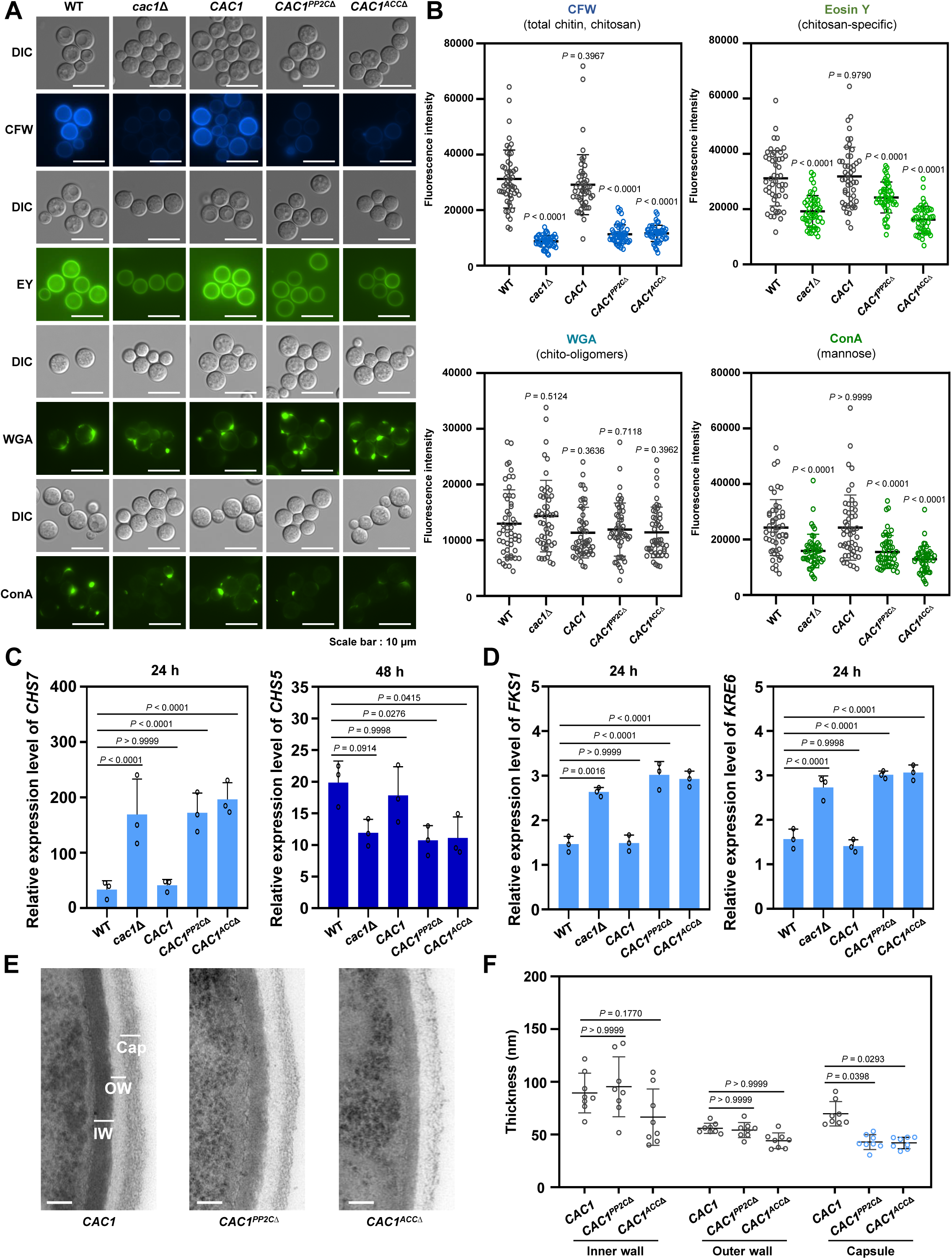
The PP2C domain is required for cell wall remodelling in *C. neoformans*. (A) Cell wall staining of the indicated strains with calcofluor white (CFW), Eosin Y (EY), wheat-germ agglutinin (WGA), and concanavalin A (ConA). Representative DIC and fluorescence images are shown. Scale bar, 10 μm. (B) Quantification of panel A. Fluorescence intensity per cell (n = 50 cells per stain). Error bars indicate SEM. Statistics were performed by one-way ANOVA with Dunnett’s multiple comparisons; *P*-values denote comparisons between the wild-type and each strain. (C) qRT-PCR of *CHS7* and *CHS5* after 24 h and 48 h of growth in YPD, respectively. Cultures were synchronised to OD_600nm_ = 0.2 before incubation. Three biological replicates were performed with three technical replicates each, and relative transcript levels were calculated using the 2^−ΔΔCt^ method. Error bars indicate SEM. Same statistics analysis was applied. (D) qRT-PCR of *FKS1* and *KRE6* after 24 h in YPD following synchronisation to OD_600nm_ = 0.2. Same statistics analysis was applied. (E) Transmission electron microscopy. Indicated strains were grown overnight at 30°C in YPD medium, synchronised to OD_600nm_ = 0.2, incubated to OD_600nm_ = 0.8, fixed with 2% paraformaldehyde and glutaraldehyde, and imaged by TEM. Inner wall (IW), outer wall (OW), and capsule (Cap) are indicated. Scale bar, 100 nm. (F) Thickness measurements of IW, OW, and capsule. For each cell layer, five positions were measured per cell (n = 8 cells) and averaged. Error bars indicate SEM. Same statistics analysis was applied.

We next addressed whether the observed reductions in chitin and chitosan were due to downregulation of genes involved in chitin synthesis and deacetylation in the *CAC1* and domain deletion strains. To evaluate this possibility, we quantified the transcript levels of chitin synthase genes (*CHS1*-*CHS8*) at 6, 24, and 48 h of incubation in YPD (yeast peptone dextrose) medium (Fig. 7C and Fig. S8A). At 24 h, *CHS7* expression was increased in *cac1*Δ, *CAC1^PP2C^*^Δ^, and *CAC1^ACC^*^Δ^ strains compared to the wild-type (*P* < 0.0001 for all; Fig. 7C), suggesting an early compensatory activation of the chitin biosynthetic pathway. By 48 h, *CHS5* expression was significantly reduced in the *CAC1^PP2C^*^Δ^ and *CAC1^ACC^*^Δ^ strains (*P* = 0.03 and *P* = 0.04, respectively), with a similar but non-significant decrease in the *cac1*Δ strain (*P* = 0.09; Fig. 7C). In contrast, other *CHS* genes showed no significant changes in expression at 6, 24, or 48 h (*P* > 0.05 for all; Fig. S8A). However, the modest *CHS5* decrease alone cannot fully explain the markedly reduced chitin and chitosan observed in *cac1*Δ, *CAC1^PP2C^*^Δ^, and *CAC1^ACC^*^Δ^ strains, suggesting the involvement of additional factors.

Beyond the chitin synthases, we examined other cell wall regulators in *C. neoformans*: *AGS1* (α-1,3-glucan synthase)^66^, *FKS1* (β-1,3-glucan synthase)^67^, and *KRE6*/*SKN1* (membrane proteins required for β-1,6-glucan biosynthesis)^68^. In line with *CHS7* expression, *FKS1* and *KRE6* expression levels increased in *cac1*Δ, *CAC1^PP2C^*^Δ^, and *CAC1^ACC^*^Δ^ strains at 24 h (*P* = 0.0016 for *FKS1* in *cac1*Δ, and *P* < 0.0001 for all others; Fig. 7D), supporting a compensatory activation to maintain cell wall integrity. In contrast, *AGS1* and *SKN1* remained comparable to wild-type levels at all time points (*P* > 0.05 for all; Fig. S8B). We also assessed expression of the chitin deacetylases *CDA1*, *CDA2*, and *CDA3* under the same conditions. Interestingly, while *CDA1* expression was slightly elevated in the *cac1*Δ strain at 48 h (*P* = 0.0168), transcription of these deacetylases otherwise remained at wild-type levels across the mutant strains (*P* > 0.05 for all others; Fig. S8C). Together, these data indicate that the diminished chitosan in the *cac1*Δ and domain deletion strains is not driven by a lack of deacetylase transcription. Rather than transcriptional repression, the concurrent loss of both chitin and chitosan suggests a defect in post-transcriptional or metabolic regulation.

To visualise whether the altered cell wall staining patterns were accompanied by morphological alterations, we examined cells grown under basal YPD conditions by transmission electron microscopy (TEM) and measured the thickness of the inner wall, outer wall, and capsule (Fig. 7E and 7F). Both the *CAC1^PP2C^*^Δ^ and *CAC1^ACC^*^Δ^ strains showed reduced capsule thickness relative to the *CAC1* strain, even in the absence of capsule-inducing conditions (*P* < 0.05 for both; Fig. 7E and 7F). In contrast, the thickness of the inner and outer wall layers did not differ significantly among the strains (*P* > 0.05 for both; Fig. 7E and 7F). Notably, the reduced chitin, chitosan, and mannan staining described above occurred without a corresponding change in inner and outer wall thickness, suggesting that the domain deletions alter the composition, organisation, or surface exposure of cell wall components rather than the overall dimensions of the wall. Together, these findings suggest that both the ACC and PP2C domains contribute to cell surface organisation in *C. neoformans*, while TEM further reveals a capsule defect that is already apparent under non-inducing conditions.

### PP2C– and ACC-specific transcriptional programs mediated by Cac1

Building on our finding that the Cac1-PP2C domain is essential for *C. neoformans* pathobiology, we performed RNA sequencing on wild-type, *cac1*Δ, *CAC1*, *CAC1^PP2C^*^Δ^ and *CAC1^ACC^*^Δ^ strains cultured in minimal medium (YNB medium) with or without 2% glucose (MMG and MM; DEGs: *P* < 0.05 and |log_2_FC| > 1) (Fig. 8A). Transcriptional perturbation was markedly greater in carbon-starved MM than in glucose-supplemented MMG: the *cac1*Δ strain displayed 1,259 DEGs in MM versus 248 in MMG; *CAC1^ACC^*^Δ^ showed 1,059 versus 211; and *CAC1^PP2C^*^Δ^ exhibited 376 versus 71. These results indicate that Cac1, especially its cAMP-generating activity, has heightened functional importance under nutrient limitation (Fig. 8B). Consistent with this, the principal component analysis (PCA) plot revealed more distinct separation of MM samples, with the *cac1*Δ and *CAC1^ACC^*^Δ^ strains clustering closely together (Fig. S9).

**Fig 8.**
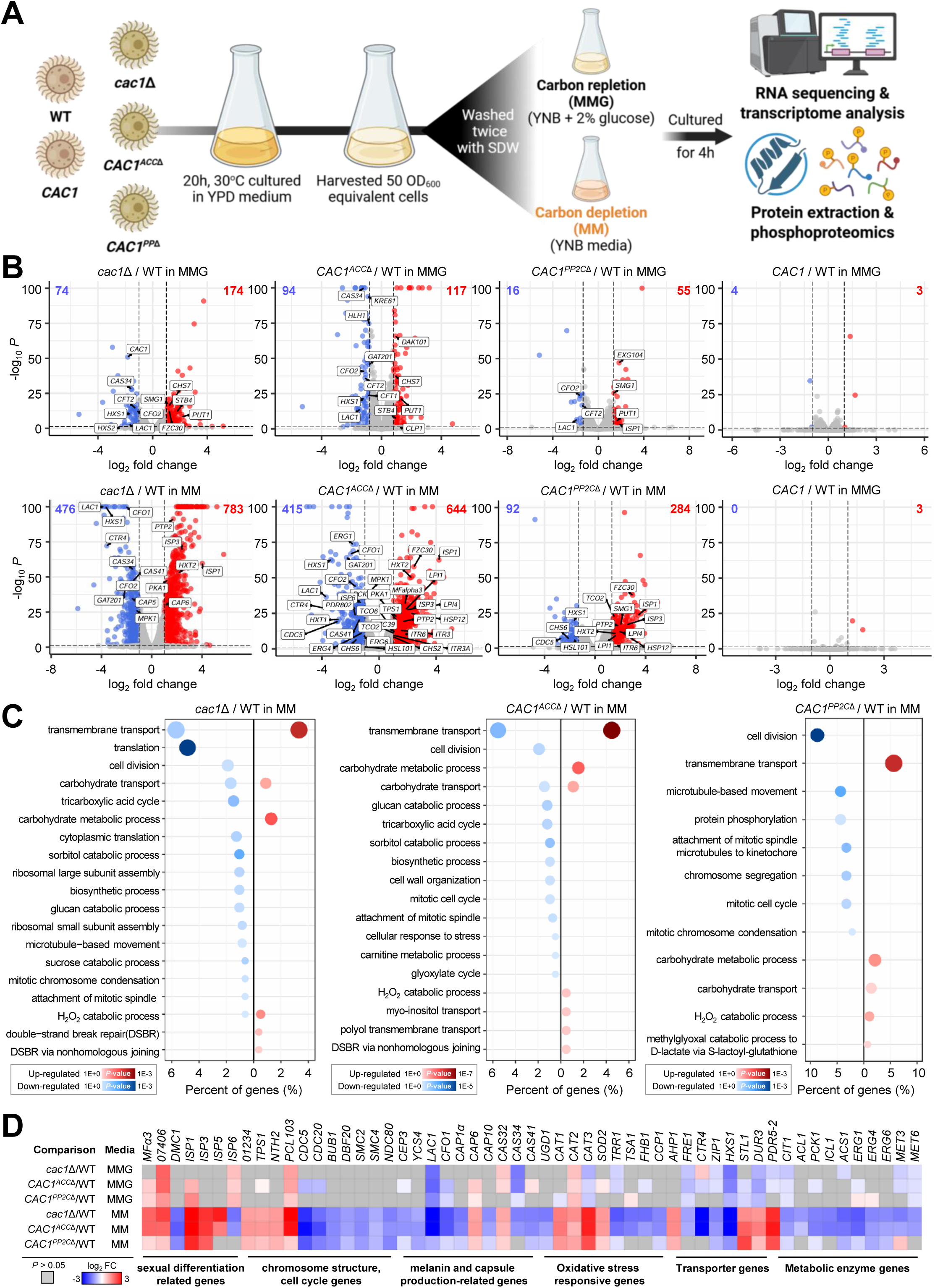
Transcriptomic profiling of *CAC1* domain mutants reveals domain-specific regulatory functions under carbon starvation and repletion conditions. (A) Schematic overview of the experimental design. Wild-type (WT), *cac1*Δ, and domain complementation strains (*CAC1^ACC^*^Δ^ and *CAC1^PP2C^*^Δ^) were cultured 20 hours in YPD, and harvested 50 OD600 equivalent cells. These cells were washed with PBS, sterilised distilled water twice. Cultures were then split into two conditions: MM (carbon depletion) and MM supplemented with 2% glucose (MMG; carbon repletion). Cultures were cultured for 4 hours, and RNA was extracted for RNA-seq and transcriptome analysis, and proteins were harvested for phosphoproteomic profiling. Created with BioRender.com (https://BioRender.com/nlretcs). (B) Volcano plot showing the number of significantly upregulated (red) and downregulated (blue) genes in each mutant strain relative to WT under MM or MMG conditions. Differential expression was defined as genes with *P* < 0.05 and |log_2_ fold change| > 1. selected genes of interest are labelled. (C) Biological process GO term enrichment analysis of significantly upregulated (red, right) and downregulated (blue, left) genes in mutants under MM conditions. Circle size reflects gene count. Cellular component, molecular functions, GO term enrichment analysis of MMG conditions are in supplementary figure 10. (D) Heatmap showing log_2_ fold changes of selected representative genes across all comparisons, chosen as markers of major and putative virulence-associated genes (melanin, iron uptake, capsule, and pheromone, developmental signalling). Genes were categorised based on function (e.g., melanin, capsule, mating, metabolism). Grey tiles indicate non-significant changes (*P* > 0.05), while red and blue represent up– and downregulation, respectively.

Comparison of domain-specific profiles showed that *CAC1^ACC^*^Δ^ largely phenocopied the full *cac1*Δ, implicating loss of ACC activity as the principal driver of broad transcriptional reprogramming. In MM, both *cac1*Δ and *CAC1^ACC^*^Δ^ exhibited up-regulation of genes involved in carbohydrate metabolic processes, oxidative stress responses, membrane components and transporters, and DNA repair, whereas down-regulated genes were strongly enriched for translation and ribosome biogenesis, certain carbon metabolism (tricarboxylic acid cycle, sorbitol catabolic process, glycolysis/gluconeogenesis, glyoxylate/pyruvate metabolism), pyridoxal phosphate binding, and amino acid biosynthesis. These patterns indicate that loss of cAMP production shifts cells into a stress-adapted, low-biosynthesis state under carbon limitation (Fig. 8C).

By contrast, *CAC1^PP2C^*^Δ^ produced a much narrower transcriptional impact. Although it showed modest overlap with the stress– and metabolism-associated up-regulation seen in *cac1*Δ and *CAC1^ACC^*^Δ^, its dominant and distinct signature was strong enrichment for down-regulated cell-cycle and cytoskeletal programs, including cell division, microtubule-based movement, kinetochore and microtubule cellular components, and microtubule motor activity, with KEGG enrichments for motor proteins and cell cycle (Fig. S10). Only a small set of genes (28 in MM, 7 in MMG) were uniquely affected in *CAC1^PP2C^*^Δ^, many of which are hypothetical or near the significance threshold (Fig. S11). Among these, several stress-related (*SRX1, ISP1*) and membrane-associated transporter genes imply that the PP2C domain contributes to redox balance and membrane remodelling under nutrient stress (Supplementary Table S3). Together, these domain-resolved results suggest that while the Cac1-ACC domain mediates the global nutrient-responsive metabolic and biosynthetic program, the Cac1-PP2C domain primarily fine-tunes mitotic and cytoskeletal circuits under nutrient stress.

Under MMG, the overall DEG burden decreased across all genotypes; nevertheless, *cac1*Δ and *CAC1^ACC^*^Δ^ still failed to fully induce carbohydrate-catabolic and oxidoreductase genes, while retaining modest enrichment of transporter/membrane terms, consistent with incomplete transcriptional adaptation to glucose in the absence of cAMP signalling. By contrast, *CAC1^PP2C^*^Δ^ showed minimal transcriptional changes in MMG.

We also found that development– and virulence-associated modules were differentially affected by the Cac1-ACC and Cac1-PP2C domains. In MM, spore/pheromone response genes (*ISP1*, *MF*α*3*) were up-regulated, whereas melanin biosynthesis (*LAC1*), iron-uptake genes (*CFO1/2*, *CFT1/2/4*), and putative capsule-associated genes (*CAS34*, *CAS41*) were down– regulated in *cac1*Δ and *CAC1^ACC^*^Δ^ strains (Fig. 8D). These genes have been functionally validated in previous studies and serve as representative markers illustrating how the Cac1-ACC domain modulates pathogenicity-related pathways^69–73^, thereby linking the ACC-dependent metabolic program to the virulence defects observed in the *cac1*Δ and *CAC1^ACC^*^Δ^ strains. The Cac1-PP2C domain also moderately influences virulence gene expression (Fig. 8D), suggesting that both domains contribute to pathogenic programs, albeit with differing breadth and emphasis.

### PP2C– and ACC-specific phosphoproteomic signatures of Cac1

Given the evident Ser/Thr-phosphatase activity of the Cac1-PP2C domain, we hypothesised that Cac1 influences the phosphorylation status of target proteins through two distinct mechanisms: i) direct dephosphorylation by the Cac1-PP2C domain and ii) cAMP production by the Cac1-ACC domain, potentially in coordination with the Cac1-PP2C domain, which activates PKA to phosphorylate downstream targets. To test this hypothesis, we performed phosphoproteomic profiling under the same experimental conditions as the transcriptome analysis (Fig. 8A, Supplementary Table S4).

Comparison of wild-type and *CAC1* strains to *cac1*Δ or domain-deletion mutants (P < 0.05 and |log_2_FC| > 1 cutoff) revealed widespread remodelling of the phosphoproteome, with the strongest effects observed under glucose-depleted minimal medium. In MM, the *cac1*Δ mutant exhibited 2,527 phosphosites with increased and 1,990 with decreased abundance, spanning **∼**1,646 unique proteins after subtracting background changes observed in *CAC1* and wild-type controls (Fig. 9A). This effect was markedly attenuated in glucose-replete MMG conditions (787 increased, 1,321 decreased sites; 1,039 proteins), suggesting that Cac1’s regulatory influence on phosphorylation is nutrient-sensitive and most pronounced during starvation.

**Fig 9.**
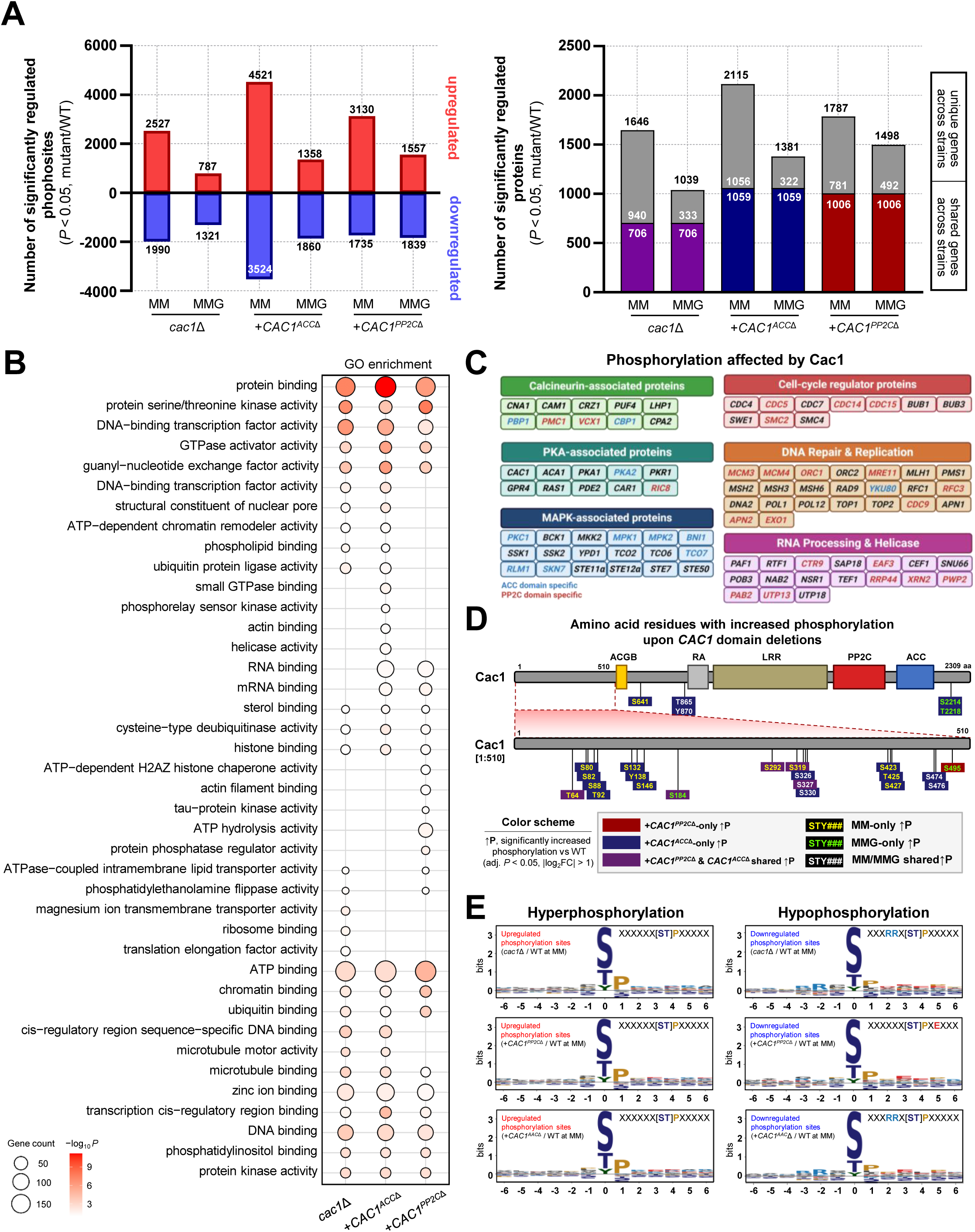
Phosphoproteomic analysis of Cac1 and its domain-specific functions. (A) Quantification of phosphosites (left) that are upregulated (red) or downregulated (blue) in the *cac1*Δ, *CAC1^ACC^*^Δ^, and *CAC1^PP2C^*^Δ^ mutants compared with the wild-type under MM or MMG. Corresponding unique proteins numbers are shown in the right panel. Each bar represents the number of significantly altered phosphosites (|log_2_FC| > 1, adjusted P < 0.05), and the overlaid labels indicate the total number of phosphosites and proteins exhibiting differential phosphorylation. (B) Biological process GO enrichment analysis of proteins with altered phosphorylation in *cac1*Δ and domain deletion cells under MM. (C) Functional categorisation of Cac1-regulated phosphorylation proteins and categorisation. PP2C domain–specific targets are indicated in red, and ACC domain–specific targets are shown in blue. (D) Schematic representation of the Cac1 protein structure showing the AC and PP2C phosphatase domains. Differentially phosphorylated sites mapped onto each domain are indicated. Sites unique to the *+CAC1^PP2C^*^Δ^ are shown in red, those unique to the *+CAC1^ACC^*^Δ^ are in blue, and sites shared by both are in grey. Media-specific phosphorylation events are also indicated (MM only, MMG only, shared). (E) Sequence motif analysis of upregulated (red) and downregulated (blue) phosphorylation sites identified in *cac1*Δ versus wild-type and in domain-deletion mutants (*+CAC1^PP2C^*^Δ^, *+CAC1^ACC^*^Δ^) under MM conditions. Consensus logos were generated from ±6 amino acid flanking sequences.

Analysis of the domain-deletion mutants revealed that loss of the Cac1-ACC domain largely recapitulated the phosphoproteomic profile of the *cac1*Δ mutant. The *CAC1^ACC^*^Δ^ strain displayed 4,521 up– and 3,524 down-regulated phosphosites in MM, affecting 2,115 proteins, with substantial overlap with the *cac1*Δ phosphoproteomic signature. Notably, deletion of the PP2C domain also led to extensive phosphoproteomic remodelling (3,130 up, 1,735 down in MM; 1,787 proteins) in contrast to its relatively limited contribution to transcriptional regulation. These findings highlight that the Cac1-PP2C domain acts predominantly through post-translational mechanisms rather than broad transcriptional control.

Principal component analysis further supported these domain-specific roles. In MMG, the phosphoproteomes of *CAC1^PP2C^*^Δ^, *CAC1^ACC^*^Δ^, and *cac1*Δ were distinctly separated from one another, indicating that each domain contributes uniquely to phosphorylation dynamics even under glucose-replete conditions. In contrast, under glucose-depleted MM conditions, *CAC1^ACC^*^Δ^ and *CAC1^PP2C^*^Δ^ clustered closely together, apart from both WT and *CAC1*, which themselves formed a separate group distinct from the domain-deleted and full knockout mutants (Fig. S12). These PCA patterns suggest that the ACC and PP2C domains function largely independently, with partially overlapping roles under nutrient-rich conditions, but act more coordinately during nutrient starvation.

Gene ontology (GO) analysis revealed that phosphoproteins altered in the *cac1*Δ mutant were enriched for functions related to protein binding, kinase and transcription factor activity, and chromatin association (Fig. 9B). Across all mutants, commonly affected categories included signal transduction and transcriptional regulation. Notably, the *CAC1^ACC^*^Δ^ mutant uniquely impacted proteins associated with microtubule, DNA, and phosphatidylinositol binding, consistent with increased phosphorylation of key MAPK cascade components (Pkc1, Bck1, Ssk1, Ssk2, Mpk1, Mpk2, Ste11α, Ste7) and PKA-related proteins (Pka1, Pka2, Cac1, Aca1, Tps1, Pde2). Additionally, phosphosite enrichment extended to the calcineurin–Crz1 pathway, with hyperphosphorylation detected on Cna1, Cam1, and Crz1, along with downstream Ca² regulators Pmc1 and Vcx1. These data indicate that Cac1 and its Cac1-ACC domain integrate multiple major signalling modules, including MAPK, PKA, and calcineurin–Crz1 pathways (Fig. 9C).

We next sought to identify phosphorylation events uniquely dependent on the Cac1-PP2C domain. Cross-strain comparisons revealed a distinct subset of phosphosites that were altered exclusively in *CAC1^PP2C^*^Δ^ or shared between *CAC1^PP2C^*^Δ^ and *cac1*Δ, suggesting PP2C-specific regulatory targets. These included key cell-cycle regulators (Smc2, Cdc14, Cdc5, Cdc15), DNA repair and replication proteins (Mcm3, Mcm4, Orc1, Mre11), and multiple RNA processing and helicase components (Ctr9, Eaf3, Rrp44, Xrn2, Pwp2, Pab2, and Utp13) (Fig. 9C). GO enrichment of these PP2C-dependent phosphoproteins highlighted functions associated with the nucleolus, DNA repair, RNA processing, and cell-cycle regulation. In addition, dozens of other cytosolic and signalling proteins displayed selective phosphorylation changes in *CAC1^PP2C^*^Δ^. Collectively, these results indicate that the Cac1-PP2C domain modulates a specialised regulatory layer—centred on chromatin, mRNA processing, and mitotic control—distinct from the canonical cAMP-PKA signalling output.

Interestingly, Cac1 itself exhibited extensive hyperphosphorylation across multiple sites in the domain-deletion backgrounds. These phosphosites clustered within the N-terminal region (aa 64–495) and were particularly prominent under MM conditions. Most sites were shared between *cac1*Δ and *CAC1^ACC^*^Δ^; however, phosphorylation at S495 was uniquely elevated in *CAC1^PP2C^*^Δ^, suggesting domain-specific feedback regulation in which the PP2C module suppresses aberrant self-phosphorylation of Cac1 (Fig. 9D). These findings imply that Cac1 undergoes autoregulatory or compensatory kinase activity when its domains are disrupted, with the Cac1-PP2C module functioning as a local negative regulator to maintain proper phosphorylation homeostasis.

To dissect the nature of kinase activity shifts, we performed de novo motif analysis on phosphosites that were increased in *cac1*Δ, *CAC1^ACC^*^Δ^, and *CAC1^PP2C^*^Δ^ (Fig. 9E). Across all mutants, phosphosites with increased abundance were significantly enriched for the [S/T]–P motif, consistent with enhanced activity of proline-directed kinases such as MAPKs and cyclin-dependent kinases (CDKs) when Cac1 function is compromised^74,75^. In contrast, phosphosites showing decreased abundance in *cac1*Δ and *CAC1^ACC^*^Δ^ were enriched for R–R–x–[S/T]–P motifs, characteristic of PKA substrates, indicating that loss of the ACC domain attenuates basophilic PKA signalling toward motifs enriched in basic residues at the −2/−3 positions relative to the phosphoacceptor site^76,77^. This pattern is consistent with the established role of the Cac1-ACC domain in generating cAMP to activate PKA. By comparison, in the *CAC1^PP2C^*^Δ^ strain, down-regulated phosphosites were enriched for the [S/T]–P–x–[E/D] motif, an extended MAPK consensus sequence, indicating proline-directed phosphorylation with a preference for acidic residues at +3. These data suggest that the Cac1-PP2C module is required to sustain phosphorylation of a subset of MAPK-associated, acidophilic substrates, likely by modulating downstream phosphoregulatory networks rather than by broadly reshaping the transcriptional program. These domain-specific motif patterns were recapitulated under glucose-replete conditions (MMG) (Fig. S13).

Collectively, these phosphoproteomic analyses demonstrate that Cac1 orchestrates nutrient-responsive phosphorylation through two coordinated mechanisms encoded within its domains. The Cac1-ACC domain broadly drives cAMP–PKA signalling and MAPK-related phosphorylation, whereas the Cac1-PP2C domain fine-tunes nuclear and cell-cycle-associated phosphorylation events. This domain bifurcation underscores the modular logic of the Cac1 architecture and highlights the plasticity of kinase-phosphatase networks during metabolic stress.

## DISCUSSION

Fungal ACs possess a distinctive multidomain architecture comprising GB, RA, LRR, PP2C, and ACC domains, unlike their human and bacterial counterparts that contain only the catalytic ACC domain. Depending on upstream regulators, fungal ACs may harbour both GB and RA domains or only one of them. While recent work has implicated the LRR domain in host adaptation and virulence attenuation in *C. neoformans*^78^, the role of the conserved PP2C domain has remained elusive. Here, we functionally characterise this domain in *C. neoformans* and reveal that the Cac1-PP2C domain functions as an active Ser/Thr phosphatase that modulates key developmental and virulence-associated processes, including melanin and capsule biosynthesis, sexual differentiation, and titan cell formation, through coordinated control with the ACC domain. Mechanistically, loss of the PP2C domain disrupts the balance between kinase and phosphatase activities, leading to dysregulated signalling outputs that reshape both fungal physiology and host immune responses. Notably, *CAC1^PP2C^*^Δ^ elicited a pronounced Th2-biased response with severe pulmonary damage yet failed to sustain systemic dissemination in mice, leading to complete avirulence and eventual fungal clearance (summarised in Fig. 10).

**Fig 10.**
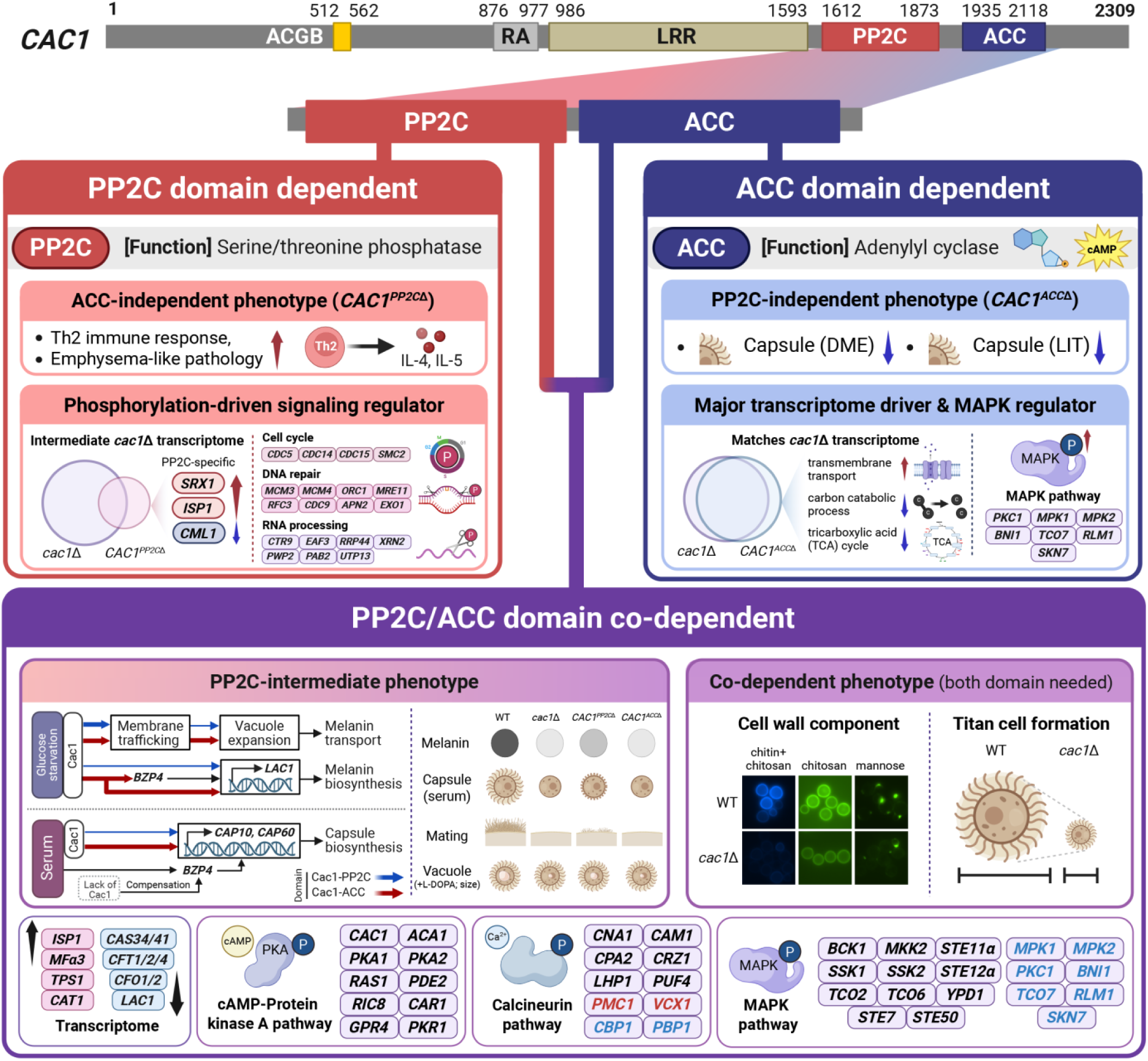
Domain-specific and co-dependent roles of the Cac1-PP2C and Cac1-ACC domains. Schematic summary of phenotypes and regulatory pathways controlled by the PP2C and ACC domains of Cac1. PP2C-dependent phenotypes and pathways are represented in red, whereas ACC-dependent functions are shown in blue. The phenotypes and pathways requiring coordinated activity of both domains are depicted in purple. Created with BioRender.com (https://BioRender.com/nlretcs).

In this study, we demonstrated that the Cac1-PP2C domain exhibits Ser/Thr-specific phosphatase activity despite its noncanonical structural configuration relative to other PP2C enzymes. Because the full structure of Cac1, including its PP2C domain, has not yet been experimentally resolved, current insights into its domain boundaries, interdomain linkages, and divergence from canonical PP2C and PPM family members rely on predictive modelling. Studies of bacterial PP2Cs have shown that a flexible flap domain can regulate substrate access to the catalytic site by toggling between open and closed conformations, and in some cases, serves as an allosteric inhibition site^35^. In contrast, structural predictions for Cac1 suggest that its flap domain is displaced from the active site and instead interacts with the LRR4-LRR10 regions. When modelled using bacterial PP2C templates, the flap domain aligns near the active site as expected; however, integrating this configuration into a model of the full LRR-containing Cac1 protein causes β-sheet disruptions and global misfolding, indicating that such a bacterial-like arrangement is structurally incompatible. Instead, the overall architecture of the Cac1-PP2C flap domain more closely resembles that of human or *Arabidopsis* PPM phosphatases. Definitive elucidation of its spatial organisation and regulatory role will require high-resolution structural studies.

The noncanonical active-site composition of Cac1-PP2C, in which several conserved metal-coordinating Asp residues are replaced by Gly, Met, His, and Gln, also raises the question of its metal dependence. In this study, Mn^2+^ was used throughout recombinant expression, purification, and enzymatic assays to ensure stable folding and reproducible activity, as the protein was largely insoluble in its absence. We therefore cannot exclude the possibility that other divalent cations, such as Mg^2+^, Ca^2+^, or Zn^2+^, may substitute for Mn^2+^ and support catalysis. Beyond catalysis, these structural features may also shape how the domain is regulated. The predicted displacement of the flap domain away from the catalytic centre suggests that access to the active site may be constrained by interdomain architecture rather than by intrinsic flap flexibility alone, while the noncanonical arrangement of metal-coordinating residues may alter metal-dependent catalytic tuning relative to classical PP2Cs. Together, these differences raise the possibility that Cac1-PP2C is regulated through mechanisms distinct from those of standalone PP2C phosphatases. Its divergence from human PP2Cs and the fungal-specific ACC-PP2C architecture further suggest that this domain may provide a basis for evaluating selective inhibition of fungal Cac1 signalling. A systematic investigation of these properties, encompassing kinetic characterisation with alternative divalent cations, structural determination of metal-binding modes, and consideration of physiologically relevant cation availability, represents an important direction for future work.

Our domain-specific deletion analyses revealed that the Cac1-PP2C domain shares substantial functional overlap with the Cac1-ACC domain, suggesting that the PP2C module contributes to autoregulation of AC activity. Consistent with this notion, deletion of the PP2C domain caused partial defects in melanin and capsule production, sexual reproduction, and titan cell formation—phenotypes that were almost completely abolished in the ACC-domain deletion mutant. Notably, the requirement for the PP2C domain appears to be context-dependent. The *CAC1^PP2C^*^Δ^ strain exhibited consistent melanin defects across all inducing conditions, whereas its contribution to capsule formation varied by medium. Capsule production in DME and LIT media remained comparable to wild-type levels, but a pronounced defect was observed under serum-induced conditions. These results suggest that capsule-inducing cues in DME and LIT media differ from those in FBS-containing medium, and that the latter requires the PP2C domain for full signalling. Together, these findings raise the possibility that an unidentified serum component promotes capsule formation via the Cac1-PP2C domain, which is dispensable for sensing capsule-inducing signals from DME or LIT media. Such AC-dependent serum sensing appears to be conserved across fungal pathogens. In *C. albicans*, for example, the yeast-to-hyphae transition is induced by serum in an AC-dependent manner^79^. Similarly, AC-driven signalling governs filamentation in *Candida tropicalis* and *Candida dubliniensis*, which suggests that AC-dependent serum sensing is likely a conserved feature within human fungal pathogens^80–82^. In future studies, the specific serum component(s) and the mechanism by which serum-derived signals engage the PP2C-dependent pathway remain to be elucidated.

The Cac1-PP2C domain was also required for key developmental processes in *C. neoformans*, though its contribution varied by pathway. While it played a partial role in sexual differentiation, it was indispensable for titan cell formation, a hallmark morphological trait linked to pathogenicity. Restoration of titan cell formation in the *CAC1^PP2C^*^Δ^ strain by exogenous cAMP indicates that the Cac1-PP2C domain functions within the cAMP signalling cascade, likely in coordination with the ACC domain. Furthermore, although the reduced induction of *PDR802* appeared to serve as a compensatory response to relieve inhibitory pressure under diminished signalling, this adjustment was still insufficient to restore titan cell formation in the absence of the Cac1-PP2C domain, highlighting its essential regulatory role in the developmental pathway.

Beyond canonical cAMP-dependent phenotypes, our study uncovered a role for Cac1 in maintaining cell wall architecture, particularly in regulating chitin/chitosan and mannan/mannoprotein composition—a function requiring both the ACC and PP2C domains. Expression profiling of chitin biosynthetic genes showed a biphasic response to cAMP pathway perturbation, characterised by early induction of *CHS7* followed by reduced *CHS5* expression. In *S. cerevisiae*, *CHS7* encodes an essential ER export chaperone for trafficking Chs3, the major chitin synthase^83^, whereas *CHS5* encodes a Golgi-associated factor required for Chs3 delivery from the trans-Golgi network to the plasma membrane^84^. Accordingly, *CHS7* upregulation likely reflects a compensatory response to impaired ER-to-Golgi trafficking under disrupted cAMP signalling, while *CHS5* downregulation suggests defective Golgi-mediated trafficking. This interpretation is supported by our vacuole staining assays, which revealed that both the PP2C and ACC domains are required for proper vesicle trafficking, as evidenced by diminished FM4-64 uptake and smaller vacuoles. Such trafficking defects have broad implications, given the central role of the secretory pathway in coordinating multiple interconnected processes^85^. For instance, laccase transport is essential for melanin synthesis, as melanin is produced within vesicles that must be properly trafficked and retained at the cell wall^46^. The same trafficking machinery also mediates the export of capsule polysaccharides from their intracellular synthesis sites to the cell surface^86^. Consequently, disruption of cAMP signalling leads to a chitin/chitosan– and mannan/mannoprotein-deficient cell wall that not only impairs anchoring and retention of melanin-containing vesicles and capsule polysaccharides but also exacerbates trafficking defects, thereby hindering the proper delivery of enzymes and substrates required for their synthesis.

Notably, deletion of the PP2C domain elicited a host immune response distinct from that caused by ACC-domain loss, inducing an early and sustained Th2 polarisation with elevated IL-4 and IL-5, prolonged fungal persistence in the lungs, and severe but nonlethal pulmonary injury. In wild-type infection, rapid clinical decline and mortality precluded assessment of prolonged pulmonary damage, whereas the *CAC1^PP2C^*^Δ^ strain allowed extended observation of chronic, emphysema-like tissue injury despite sustained lung colonisation. During the early phase of infection, *CAC1^PP2C^*^Δ^-infected mice showed increased infiltration of macrophages and antigen-presenting cells, with levels intermediate between those of the *CAC1* strain and the non-infected, *cac1*Δ, and *CAC1^ACC^*^Δ^ groups. This cellular response was accompanied by an early increase in Th2 cytokines, suggesting rapid establishment of a Th2-dominant immune environment that may promote chronic tissue remodelling rather than effective fungal clearance. Mechanistically, these immune outcomes likely arise from PP2C-dependent regulation of cell wall and capsule remodelling, which modulates key immunoregulatory signals at the fungal surface. Such surface imbalance may reprogram host recognition toward type-2 immunity, analogous to the polysaccharide remodelling in *Aspergillus fumigatus* that drives Th2-skewed pulmonary inflammation^87^. Similarly, *Cryptococcus gattii* infections in immunocompetent hosts are characterised by lung-restricted persistence and dampened Th1/Th17 polarisation^88^, supporting a model in which prolonged fungal residence and altered surface signalling favour a type-2–biased immune environment. However, whereas *A. fumigatus* infection produces eosinophil-rich, bronchocentric inflammation and *C. gattii* induces localised M2 macrophage–driven granulomatous lesions with limited tissue damage, *C. neoformans CAC1^PP2C^*^Δ^ infection results in diffuse parenchymal destruction and emphysema-like changes without systemic dissemination. Together, these findings highlight the PP2C domain of *C. neoformans* Cac1 as a key interface module that maintains fungal surface integrity and ensures balanced host immune interpretation.

Transcriptomic profiling highlights a dominant role of the Cac1-ACC domain during carbon limitation, confirming that cAMP synthesis is the principal driver of transcriptional remodelling under nutrient starvation. In carbon-poor environments, Cac1-mediated signalling shifts the cellular program from anabolic growth to stress adaptation, coordinating carbon utilisation, oxidative defence, and virulence-associated processes such as capsule and cell-wall synthesis. These observations reinforce the established model that cAMP–PKA signalling globally reprograms metabolism to sustain survival under host-like nutrient stress^89^. By contrast, the PP2C domain contributed little to the transcriptional landscape, implying that its major role lies beyond transcriptional regulation. The limited gene expression changes observed mainly in vesicle trafficking and cytoskeletal organisation align with the Cac1-PP2C deletion phenotypes. This supports the view that the PP2C module acts post-translationally to stabilise phosphorylation networks downstream of cAMP signalling, refining rather than initiating transcriptional responses.

The phenotypic complexity observed upon disruption of Cac1 domains is explained by its phosphoregulation of key nodes across multiple signalling and effector modules. At the signalling level, both Cac1-ACC and Cac1-PP2C domains modulate distinct yet intersecting cascades, including PKA, MAPK, and calcineurin. Targets such as Pka1/2, Aca1, and downstream effectors like Car1 link Cac1 to classical cAMP–PKA signalling that governs metabolic adaptation, capsule biosynthesis, and virulence gene expression^8,89–91^. ACC-dependent phosphorylation of MAPKs, including Mpk1/2 and upstream components, suggests that cAMP output interfaces with the MAPK network to influence cell wall integrity and mating^92,93^. The broad reach of the Cac1-ACC domain across these kinase modules explains why its deletion causes severe impairment across all major phenotypes, from capsule to titan cell formation. The PP2C-specific phosphorylation enrichment of the calcineurin-Crz1 pathway and Ca^2+^ regulators, including Crz1, Pmc1, and Vcx1, indicates that it contributes to Ca² –sensitive stress adaptation, vesicle transport, and ion homeostasis^94–96^. The dysregulation of this module likely explains the abnormal cell wall remodelling and altered cell surface composition observed in the *CAC1^PP2C^*^Δ^ strain, which in turn provokes a Th2-skewed immune response. Similarly, phosphoregulation of key cell-cycle proteins and DNA repair complexes aligns with the titan cell defect, as such extreme cell enlargement demands tightly regulated nuclear division and replication^60,97^. Furthermore, the PP2C domain modulates numerous RNA processing and helicase components (Rrp44, Xrn2, Pwp2, Utp13), suggesting a role in post-transcriptional buffering of stress-responsive and morphogenesis-related gene expression^98^. Disruption of RNA maturation and ribonucleoprotein assembly may underlie the observed defects in vesicle production and membrane trafficking, particularly given the close coupling between mRNA processing and secretory capacity in fungi^99^.

In conclusion, our findings demonstrate that Cac1 functions through a dual-domain architecture in which the ACC and PP2C modules exert complementary control over phosphorylation networks. The ACC domain amplifies nutrient-derived cAMP signals through PKA, MAPK, and calcineurin cascades to coordinate metabolic and virulence responses, while the Cac1-PP2C domain fine-tunes these outputs by stabilising phosphorylation balance across chromatin, replication, vesicle trafficking, and post-transcriptional pathways. This layered regulation enables *C. neoformans* to precisely modulate morphogenesis, capsule and melanin synthesis, titan cell formation, and immune evasion without compromising signalling robustness. Loss of the ACC domain abolishes, and loss of the PP2C domain miscalibrates, cAMP signalling, with either defect attenuating pathogenicity. Beyond intracellular regulation, the Cac1-PP2C domain also shapes host–pathogen outcomes by inducing localised Th2-biased inflammation while restricting systemic spread. Because mammalian adenylyl cyclases lack a comparable integrated PP2C domain, this domain may serve as a basis for selective antifungal targeting. Moreover, because its noncanonical active site diverges structurally from human PP2C phosphatases and its loss attenuates virulence factors such as melanin rather than compromising viability, this domain may be amenable to selective, anti-virulence inhibition that facilitates immune clearance while limiting resistance pressure under conventional antifungal therapies.

## METHODS

### Ethics statement

Animal care and experimental procedures were reviewed and approved by the Institutional Animal Care and Use Committee of the Experimental Animal Center at Jeonbuk National University (approval number JBNU 2023–228). All animal experiments were performed in full accordance with institutional and national ethical guidelines for animal research. Biosafety-related experiments were conducted within certified biosafety cabinets in the BL3 and ABL3 facilities of the Core Facility Center for Zoonosis Research (Core-FCZR).

### Expression and purification of the PP2C domain

The PP2C domain of the *C. neoformans* (strain H99) Cac1 (residue 1595-1884) was cloned into a pET vector containing an N-terminal 10xHis tag, MBP and a tobacco etch virus (TEV) protease cleavage site. The subcloned plasmid was transformed into *Escherichia coli* BL21 (DE3) and grown in LB medium. When the OD_600_ reached 0.6∼0.8 after 6 h, cells were induced with 0.5 mM IPTG (isopropyl 1-thio-ß-D-galactopyranoside) and 2 mM MnCl_2_ at 17 overnight. The cells were harvested by centrifugation at 3000 rpm for 20 min and then sonicated (3 seconds on, 5 seconds off, at 60% amplitude, 5 min) in lysis buffer consisting of 25 mM Tris-HCl pH 8.0, 300 mM NaCl, 10 % glycerol, 30 mM imidazole, 2 mM MnCl_2_. After centrifugation at 13000 rpm for 2 h using a high-speed centrifuge, the supernatant was filtered through a 0.45 μm, 47 mm membrane filter (1215676; GVS Life Sciences, Italy). His-tagged proteins were then purified using nickel-affinity chromatography (His60 Ni Superflow Resin) and eluted with buffer consisting of 25 mM Tris-HCl pH 8.0, 300 mM NaCl, 10% glycerol, 300 mM imidazole, 2 mM MnCl_2_. The PP2C domain was then concentrated using Vivaspin 20 centrifuge concentrator with a molecular-weight cutoff of 30 kDa and further purified by size exclusion chromatography on Superdex 200 Increase gel filtration column (Cytiva, USA) with SEC buffer containing 20 mM HEPES-NaOH (pH 8.0), 300 mM NaCl, 2 mM MnCl_2_. Proteins were mixed with glycerol to a final concentration of 20%, divided into 200-μL aliquots, and stored frozen until use.

### Phospho-amino acid dephosphorylating and catalytic activity assay

Dephosphorylation of phospho-amino acid was assessed using Malachite Green Phosphate Assay Kit (Lot#307BI06A11; Sigma Aldrich, USA) with 10 μM and 1 μM concentration of phospho-serine (O-Phospho-L-serine; Sigma Aldrich, USA), phospho-threonine (O-Phospho-L-threonine; Bachem, Switzerland) and phospho-tyrosine (O-Phospho-L-tyrosine; Cayman chemical Company, USA) as a substrate. For dephosphorylation catalytic activity assay, the enzymatic assay was performed using EnzChek™ Phosphatase Assay Kit (Invitrogen, USA) which contains Molecular Probes’ patented DiFMUP as a substrate. For phospho-amino acid and dephosphorylating assay, the standard curve was measured at the 620 nm using a CLARIOstar Plus plate reader (BMG Labtech, Germany) based on quantification of the green complex formed between Malachite green and molybdate with different free orthophosphate concentrations from 0-40 μM. The amount of dephosphorylated free phosphate was measured with 10 μM recombinant proteins at RT for 30 min in a 20 μl of the working reagent to 80 μl of sample protein solution. The assay was ended by the addition of the working reagent. For dephosphorylation catalytic activity assay, the standard curve was obtained from the excitation/emission at 360 nm/460 nm wavelength plate reader CLARIOstar Plus with different DiFMU concentration from 0-100 μM. Assays with 0-200 μM DiFMUP were conducted using 40 μg purified recombinant PP2C proteins at RT at 3-min intervals (incubate protected from light) in a 50 μl reaction mixture buffer and 50 μl of sample protein solution.

### Strain construction

Domain deletion strains of *CAC1* were constructed (Supplementary Table S1) by complementing the *cac1*Δ mutant with PCR-amplified fragments containing the 5′ flanking region (including the native *CAC1* promoter) and the 3′ flanking region (including the native *CAC1* terminator) of either wild-type *CAC1* or the domain-deleted alleles (*CAC1^PP2C^*^Δ^ and *CAC1^ACC^*^Δ^). Primers used for amplification are listed in Supplementary Table S2. The amplified PCR products were cloned into the pNEO vector carrying the neomycin resistance gene (*NEO^R^*) to construct pNEO_CAC1, pNEO_CAC1*^PP2C^*^Δ^, and pNEO_CAC1*^ACC^*^Δ^ using Gibson Assembly Master Mix (New England Biolabs, USA). All constructs were sequence-verified. Each cassette was then inserted into the *cac1*Δ mutant strain by biolistic transformation, as previously described^100^. Targeted integration of the wild-type and domain-deleted *CAC1* alleles into the native locus was confirmed through diagnostic PCR.

### Protein extraction and western blot analysis

The wild-type, *CAC1*-4xFLAG, *CAC1^PP2C^*^Δ^-4xFLAG, and *CAC1^ACC^*^Δ^-4xFLAG strains were cultured in liquid YPD for 16 h at 30°C. The OD_600_ was adjusted to 0.2, and the cells were grown until they reached an OD_600_ of 0.8. For the basal condition, the cells were flash-frozen in liquid nitrogen at this stage. For the nutrient starvation condition, the cells were washed and resuspended in YNB medium without glucose and further incubated at 30°C for 2 h. For the cell wall stress condition, calcofluor white was added to a final concentration of 250 μg/ml, and the cells were further incubated at 30°C for 2 h. After each stress treatment, the cells were pelleted and flash-frozen in liquid nitrogen. For protein extraction, the frozen cells were resuspended with 0.75 mL of cell lysis buffer (50 mM Tris-HCl pH 7.5, 1% sodium deoxycholate, 5 mM sodium pyrophosphate, 0.1 μM sodium orthovanadate, 50 mM NaF, 0.1% SDS, 1% Triton X-100, 0.5 μM phenylmethylsulfonyl fluoride, and 1x protease inhibitor cocktail solution [protease inhibitor cocktail set IV, Calbiochem]) and were disrupted by bead-beating. The samples were then centrifuged at 13,000 rpm for 20 min at 4°C, and the supernatants were transferred to new tubes. Protein content was quantified using the Pierce BCA Protein Assay Kit (ThermoFisher Scientific) with bovine serum albumin (ThermoFisher Scientific) as the standard. Diluted protein samples were mixed with BCA working reagent alongside the bovine serum albumin (BSA) standards and incubated at 37°C for 30 min. After quantification, the protein concentration was adjusted to 1.5 μg/μL, 4x NuPAGE™ LDS sample buffer (ThermoFisher Scientific) was added, and the samples were stored at –20°C. To immunoblot large Cac1-4xFLAG, Cac1^PP2CΔ^-4xFLAG, and Cac1^ACCΔ^-4xFLAG proteins, samples were heated at 70°C for 5 min and resolved on 5% Tris-acetate gels at 150 V for 75 min. To immunoblot β-actin, samples were re-heated at 95°C for 10 min and resolved on 10% Tris-glycine gels, run at 50 V until the ladder began to separate and then at 100 V until the target size reached the bottom half of the gel. Because β-actin was run on a separate gel due to the large size discrepancy among the proteins, membranes were stained with Coomassie Brilliant Blue to visualize the protein loading. The following primary antibodies were used: monoclonal anti-FLAG antibody (Sigma-Aldrich, F1804, 1:2000) and β-actin antibody (Santa Cruz Biotechnology, sc-47778, 1:1000). The secondary reagent, mouse IgG Fc-binding protein conjugated to horseradish peroxidase (HRP; Santa Cruz Biotechnology, sc-525409), was used for detecting both FLAG (1:4000) and β-actin (1:2000).

### In vitro virulence factor analysis

To assess melanin production, each indicated strain was cultured in liquid YPD for 16 h at 30°C, washed twice with sterile phosphate-buffered saline (PBS), and spotted onto Niger seed (40 g/L), dopamine, or epinephrine media (1 g L-asparagine, 3 g KH_2_PO_4_, 0.25 g MgSO_4_, 1 mg thiamine, 5 μg biotin, and 100 mg L-DOPA or epinephrine hydrochloride per litre) containing 0.1% glucose. The cells were then incubated at 37°C for 1 to 3 days and photographed. For the capsule production assay, each strain was cultured in liquid YPD at 30°C for 16 h, washed twice with sterile PBS, and spotted onto Dulbecco’s modified Eagle’s (DME; Gibco, USA) agar medium, Littman’s medium^101^, and 10% foetal bovine serum (FBS) agar medium. The cells were then incubated at 37°C for two days, stained with India ink (BactiDrop; Remel, USA) and observed using a brightfield microscope (ECLIPSE Ni; Nikon, Japan). Cell body diameter and total diameter were measured for 50 random cells, and capsule thickness was calculated by subtracting cell body diameter from total diameter. Statistical significance was determined using one-way ANOVA with Bonferroni’s multiple comparison test in Prism 10.4 (GraphPad, USA).

### Vacuole staining assay

Vacuoles were visualised using FM4-64 (Invitrogen, USA) as previously described^45^, with minor modifications. Specifically, each indicated strain was cultured in liquid YPD for 16 h at 30°C. 1 ml of sample was set aside as the basal sample, and the rest of the cells were washed twice with sterile PBS and resuspended in minimal medium supplemented with L-DOPA (15 mM glucose, 10 mM MgSO_4_, 29.4 mM KH_2_PO_4_, 13 mM glycine, 3 μM thiamine, and 1 mM L-DOPA with pH adjusted to 5.5) at the OD_600_ of 0.2. The cells were then cultured at 30°C for 24 h, and 1 ml of sample was set aside as the 24 h sample. To stain the cells with FM4-64, the cells were spun down and resuspended with 100 μl of 5 μg/ml FM4-64 dye diluted in ice-cold Hanks’ balanced salt solution (HBSS; Gibco, USA), and kept on ice for 30 min. The cells were pelleted, washed three times with HBSS, and resuspended with 100 μl of HBSS. To visualise the cells, 5 μl of cells and 5 μl of mounting glue (Biomeda, USA) were thoroughly mixed, fixed for 30 min in the dark, and observed by DIC and fluorescence microscope (ECLIPSE Ni; Nikon, Japan). For quantification, fluorescence of 100 random cells was measured using ImageJ, and diameters of 100 random vacuoles were measured using Nikon software.

### Mating efficiency and titan cell formation assay

To evaluate unilateral mating efficiency, indicated strains and *MAT***a** wild-type (KN99**a**) strain were cultured in liquid YPD for 16 h at 30°C and washed twice with sterile PBS. Equal concentrations (10^7^ cells/ml) of indicated cells and *MAT***a** KN99**a** cells were mixed, spotted onto MS (Murashige and Skoog) medium (pH 5.8), and incubated at room temperature in the dark for 7-14 days. Filament and spore formation were observed using a brightfield microscope. For in vitro induction of titan cell formation, indicated cells were cultured in liquid YPD for 16 h at 30°C and washed twice with minimal media (15 mM D-glucose, 10 mM MgSO_4_, 29.4 mM KH_2_PO_4_, 13 mM glycine, 3 μM thiamine with pH adjusted to 5.5). Then, 10^6^ cells were inoculated in 1 ml of minimal media in 1.5 ml centrifuge tube and incubated at 30°C with shaking for 72 h. For cAMP supplementation assay, 10 mM of cAMP (Sigma-Aldrich, USA) was added. The cells were then spun down, observed using brightfield microscope, and cell diameter was measured for 100 random cells for quantification.

### Gene expression analysis using quantitative RT-PCR

To prepare samples for RNA extraction of melanin-regulating genes, cells were cultured in 50 ml of YPD for 16 h at 30°C. Then, OD_600_ was adjusted to 0.2, and when cells reached OD_600_ of 0.8, 10 ml of sample was spun down, frozen in liquid nitrogen, and lyophilized as basal sample. The remaining cells were washed three times with sterile PBS, resuspended in liquid yeast nitrogen base (YNB) medium without glucose, and incubated at 30°C for 2 h. The cells were then spun down, frozen in liquid nitrogen, and lyophilized. For capsule-related genes, the basal sample was prepared in the same manner, and the remaining cells were washed three times with sterile PBS, resuspended in 10% FBS, and incubated for 2 or 6 h. The cells were then spun down, frozen in liquid nitrogen, and lyophilized. To assess titan cell-regulating genes, cells were prepared in the same manner as the titan cell formation assay at a density of 10^6^ cells/ml, with a final volume of 30 ml in an air-tight 50 ml conical tube. For cAMP supplementation, 10 mM of cAMP (Sigma-Aldrich, USA) was added. To prepare samples for RNA extraction of chitin synthesis-related genes, the cells were cultured in 50 ml of YPD for 16 h at 30°C, the OD_600_ was adjusted to 0.2, and incubated in YPD at 30°C. Samples were then collected 6, 24, or 48 h later. Each sample was spun down, frozen in liquid nitrogen, and lyophilized. For RNA extraction, lyophilized samples were homogenized with 3 mm glass beads, and total RNA was extracted using an RNA extraction kit (iNtRON Biotechnology, South Korea). RNA concentration was then measured, and 5,000 ng of total RNA was used to synthesize cDNA using reverse transcriptase (ThermoFisher Scientific, USA). qRT-PCR was then performed using CFX96 (Bio-Rad Laboratories, USA) using primers designed for each target gene (Supplementary Table S2) and TB Green (ThermoFisher Scientific, USA). Unless otherwise specified as “normalized to *ACT1*,” transcript levels were calculated by the ΔΔCt method with *ACT1* as the reference gene. For targets with very low basal expression – specified by “normalized to *ACT1*” – we report ΔCt = Ct(target) – Ct(*ACT1*) per sample because ΔΔCt produced unstable fold changes. Statistical analysis was performed by one-way ANOVA or Student’s *t*-test, which is specified in each figure legend. The relative gene expression was then visualized using Prism 10.4.

### Murine infection, fungal burden, and histopathology

Animal experiments were conducted at the Core Facility Center for Zoonosis Research (Jeonbuk National University, South Korea). Specific pathogen-free/viral antibody-free (SPF/VAF)-confirmed, inbred 6-week-old female BALB/cAnNCrlOri mice were purchased from ORIENT BIO INC (South Korea). Upon arrival, mice were randomly assigned to experimental groups and acclimatised for one week before infection. Mice were housed under controlled temperature and humidity conditions with a 12-h light/dark cycle and were provided standard laboratory chow and water *ad libitum*. Investigators were aware of the experimental group and infecting strain assigned to each animal during infection procedures. For infection, *C. neoformans* strains were cultured overnight in fresh YPD medium at 30°C with shaking, and yeast cells were counted and adjusted to the required concentration. For survival and long-term survival assays, mice were anaesthetised with isoflurane administered via a vaporizer and infected intranasally with 5 × 10^5^ cells per mouse, then monitored daily for clinical condition, body weight, and survival. A loss of ≥25% of the initial body weight was defined as the humane endpoint. Mice reaching the humane endpoint were deeply anaesthetised with a high concentration of isoflurane followed by cervical dislocation. Mice infected with the *CAC1^PP2C^*^Δ^ strain were monitored for up to 111 days post-infection (dpi). Survival differences were analysed using the log-rank (Mantel–Cox) test. For fungal burden and immune response analysis, mice were infected intranasally with 3 × 10^6^ cells per mouse and euthanised at the predetermined experimental endpoints by deep anaesthesia with a high concentration of isoflurane followed by cervical dislocation. For long-term assessment of infection, *CAC1^PP2C^*^Δ^-infected mice surviving 111 dpi were euthanised using the same procedure, and the lungs and brains were collected. Organs were weighed and homogenised, and serial dilutions of tissue homogenates were plated on YPD agar to determine colony-forming units. Statistical significance was assessed using one-way ANOVA with Tukey’s multiple-comparison test. GraphPad Prism 9.5.1 was used for statistical analysis. For histopathological analysis, lungs were collected at 16 and 26 dpi, as well as from *CAC1^PP2C^*^Δ^-infected long-term survivors at 111 dpi, and fixed in 3.7% formalin. Fixed tissues were dehydrated, cleared with xylene, and embedded in paraffin blocks. Sections (5 μm) were cut and stained with Grocott’s Methenamine Silver (GMS) Stain Kit (ab287884; Abcam, UK) according to the manufacturer’s instructions. Images were acquired using a Primostar 3 microscope (ZEISS, Germany) at 400× magnification.

### Immune cell and cytokine analyses

For immune cell analysis, immune cells were isolated from the lungs of infected mice at 14 days post-infection (dpi) and washed with PBS. Fcγ receptors were blocked with anti-CD16/CD32 antibody (clone 93, Cat. No. 14-0161-82; Invitrogen, Carlsbad, CA, USA) at 1 μL/test. Subsequently, the cells were stained with surface markers, including FITC-conjugated anti-CD3 (clone 17A2, Cat. No. 11-0032-82; Invitrogen), PE-Cy7-conjugated anti-CD4 (clone GK1.5, Cat. No. 25-0041-82; Invitrogen), PerCP-Cy5.5-conjugated anti-CD8 (clone 53-6.7, Cat. No. 45-0081-82; Invitrogen), APC-conjugated anti-CD19 (clone 1D3, Cat. No. 17-0193-82; Invitrogen), FITC-conjugated anti-F4/80 (clone BM8, Cat. No. 11-4801-82; Invitrogen), PE-conjugated anti-MHC class II (I-A/I-E) (clone M5/114.15.2, Cat. No. 12-5321-82; Invitrogen), and APC-conjugated anti-CD11b (clone M1/70, Cat. No. 17-0112-82; Invitrogen). Each surface-staining antibody was used at 1 µL/test. Stained cells were analysed using a BD FACSymphony A3 flow cytometer and FlowJo V10 software.

For pulmonary cytokine analysis, infected mice were euthanised at 7, 14, and 21 dpi using the procedure described above, and lung tissues were collected. Total RNA was isolated from lung tissues and subjected to quantitative PCR as described above. The mRNA expression levels of *IFNG*, *IL4*, and *IL5* were determined using gene-specific primers and normalised to *GAPDH*. The primer sequences are listed in Supplementary Table S2.

For serum cytokine analysis, serum was collected at 7, 14, and 21 dpi, and TNF-α levels were quantified using a commercially available ELISA kit (Cat. No. 88-7324; Invitrogen) according to the manufacturer’s instructions. Briefly, 96-well immunoplates (Cat. No. 32296; SPL, South Korea) were coated with the supplied capture antibody and incubated overnight at 4°C. After washing with PBS containing 0.05% Tween-20 (PBST), the plates were blocked with 1% BSA in PBS for 1 h at room temperature. Serum samples and recombinant TNF-α standards were added in duplicate and incubated for 2 h at room temperature. The plates were then incubated with the supplied biotinylated detection antibodies, followed by streptavidin-HRP conjugate. After washing, TMB substrate solution was added, and the reaction was stopped with 2 N H SO. Absorbance was measured at 450 nm using a SpectraMax microplate reader (Molecular Devices, USA).

### Quantitative measurement of cell wall components

To quantify the levels of cell wall components, we employed a series of fluorescent stains: 1) calcofluor white (CFW) that binds β-1, 4-linked polysaccharides such as chitin and chitosan^102^, 2) Eosin Y (EY), an anionic acidic dye that binds to positively charged polysaccharides like chitosan^103^, 3) FITC-conjugated wheat germ agglutinin (WGA) that recognises exposed chito-oligomers^104^, 4) Concanavalin A (ConA) that binds to α-D-mannopyranosyl and α-D-glucopyranosyl residues found in mannans and mannoproteins^105^, and 5) Dectin-1, a C-type lectin receptor recognising β-1,3 and β-1,6-glucans^106^. For cell wall staining, cells were incubated in 50 ml of liquid YPD for 72 h at 30°C, and two sets of 1 ml samples were spun down. For one set of 1 ml samples, the cells were washed twice with sterile PBS, resuspended in PBS, and stained with 25 μg/ml of CFW (Sigma-Aldrich, USA), 100 μg/ml of WGA (Invitrogen, USA), and 100 μg/ml of ConA (Invitrogen, USA) for 30 min in the dark. For the other set of 1 ml sample, the cells were washed once with McIlvaine buffer, resuspended in PBS, and stained with 1,200 μg/ml of Eosin Y for 5 min in the dark. After staining, the cells were washed two to three times with their corresponding buffer and resuspended. For Dectin-1 staining, soluble human Dectin-1a fused to an IgG1 Fc domain (Fc-hDectin-1a; InvivoGen, USA) was used. First, the cells were cultured for 16 h in liquid YPD at 30°C, and the cells were resuspended in PBS to OD_600_ of 1.0. For each strain, 100 μl of cells was pelleted and resuspended with 100 μl of PBS containing 15 μg/ml of Fc-hDectin-1a for 1 h at room temperature. Cells were then spun down and washed three times with PBS, resuspended in PBS containing 10 μg/ml of Alexa Fluor 488-conjugated goat anti-human IgG (Invitrogen, USA), and stained in the dark for 1 h at room temperature. The cells were then washed twice with PBS. All the stained and washed cells were then mounted onto glass slides and imaged using an ECLIPSE Ni fluorescence microscope (Nikon, Japan). The fluorescence of each stain was quantified by measuring the fluorescence intensity of 50 random cells using ImageJ/Fiji software.

### Transmission electron microscopy (TEM)

TEM was performed at Yonsei Biomedical Research Institute, Yonsei University College of Medicine. Cells were grown overnight at 30 in YPD broth and synchronized to OD_600_ = 0.2 in liquid YPD medium and then further incubated until they reached OD_600_ = 0.8. The harvested cells were fixed with 2% glutaraldehyde + 2% paraformaldehyde in 0.1 M phosphate buffer (pH 7.4) and sent to Yonsei Biomedical Research Institute, Yonsei University College of Medicine for TEM analysis. The fixed cells were rinsed in the fixation buffer and post-fixed in 1% OsO_4_ (0.1 M phosphate buffer) for 2 h. Samples were dehydrated through a graded ethanol series (50, 60, 70, 80, 90, 95, 100%, 10 min each), infiltrated with propylene oxide for 10 min, and embedded in Poly/Bed 812. Polymerization was carried out at 70 for 12 h in an EM oven. Blocks were trimmed and sectioned on an Ultramicrotome (UC7). Semithin sections (200 nm) were stained with toluidine blue for light-microscopy orientation. The regions of interest were then sectioned at 80 nm, mounted on copper grids, and contrast-stained with 5% uranyl acetate (20 min) followed by 3% lead citrate (7 min). Imaging was performed on a Hitachi HT7800 TEM operated at 100 kV equipped with an RC camera. For thickness measurement, five positions were measured and averaged for each cell (8 cells per strain) at 150,000x magnification using ImageJ. For quantification of inner wall density, images were inverted to reverse black and white contrast, and the mean intensity of the inner-wall region was measured for each strain at 150,000x magnification (8 cells per strain). Statistics were performed by one-way ANOVA with Bonferroni’s multiple comparisons (****, *P* < 0.0001; NS, not significant).

### Phosphoproteome analysis

The strains of interest (wild-type H99, *cac1*Δ, *cac1*Δ::*CAC1*, *cac1*Δ::*CAC1^PP2C^*^Δ^, *cac1*Δ::*CAC1^ACC^*^Δ^) were grown in 50 ml liquid YPD for 20 h at 30°C in triplicate. Following the OD_600_ measurement, 50 OD equivalents of cells for each sample were harvested and washed twice with sterilised purified water. During the second wash, the cell suspension was split into two halves with 25 OD equivalents each into fresh 50 ml conical tubes and pelleted. One set was resuspended in 20 ml of YNB medium (minimal medium), and the other set was resuspended in 20 ml of YNB media containing 2% glucose. Cells were grown for 4 more h at 30°C with vigorous shaking. Cells were harvested by centrifugation at 4°C at 3200 rpm for 10 min and immediately frozen at –80°C. All 30 samples were then used for phosphoproteome analysis at the Duke University Proteomics and Metabolomics Core Facility. Samples were supplemented with 200 μl of 8 M urea in 50 mM ammonium bicarbonate before protein extraction using a bead beater (2 rounds at 10 sec). 175 μl was taken from each sample and were spiked with 1 or 2 pmol bovine casein as an internal quality control standard. Next, the samples were reduced for 30 min at 32°C, alkylated with 20 mM iodoacetamide for 30 min at room temperature, then supplemented with a final concentration of 1.2% phosphoric acid and 1513 μL of S-Trap (Protifi, USA) binding buffer (90% MeOH/100 mM TEAB). Proteins were trapped on the S-Trap micro cartridge, digested using 20 ng/μL sequencing grade trypsin (Promega, USA) for 1 h at 47°C, and eluted using 50 mM TEAB, followed by 0.2% FA, and lastly using 50% ACN/0.2% FA. All samples were then lyophilised. The phosphopeptide samples were resuspended in 80% acetonitrile and 1% TFA prior to TiO_2_ enrichment. Each sample was subjected to complex TiO_2_ enrichment using GL biosciences TiO_2_ tips and manufacturer-recommended protocols. All samples were frozen and lyophilised.

Quantitative LC/MS/MS was performed using an EvoSep One UPLC coupled to a Thermo Orbitrap Astral high-resolution accurate mass tandem mass spectrometer. Briefly, each sample-loaded EvoTip was eluted onto a 1.5 μm EvoSep 150 μm ID x 15 cm performance column using the SPD30 gradient at 55°C. Data collection on the Orbitrap Astral mass spectrometer was performed in a data-independent acquisition (DIA) mode of acquisition with a resolution of 240,000 (@ m/z 200) full MS scan from m/z 380-1080 in the OT with a target AGC value of 4e5 ions. Fixed DIA windows of 5 m/z from m/z 380 to 1080 DIA MS/MS scans were acquired in the Astral with a target AGC value of 5e4 and a max fill time of 8 ms. HCD collision energy setting of 27% was used for all MS2 scans. The total analysis cycle time for each sample injection was approximately 40 min. Following 34 total UPLC-MS/MS analyses (30 experimental samples and 4 internal quality control samples), data were imported into Spectronaut (Biognosis), and individual LCMS data files were aligned based on the accurate mass and retention time of detected precursor and fragment ions. Relative peptide abundance was measured based on MS2 fragment ions of selected ion chromatograms of the aligned features across all runs. The MS/MS data were searched against a *Cryptococcus neoformans* H99 database (downloaded in 2024), a common contaminant/spiked protein database (bovine albumin, bovine casein, yeast ADH, etc.), and an equal number of reversed-sequence “decoys” for false discovery rate determination. A library was created using only LC-MS data files collected within this study. Database search parameters included fixed modification on Cys (carbamidomethyl) and variable modification on Met (oxidation), Protein N-term (acetyl), and Ser/Thr/Tyr (phos). Full trypsin enzyme rules were used along with 10 ppm mass tolerances on precursor ions and 20 ppm on product ions. Spectral annotation was set at a maximum of 1% peptide false discovery rate based on q-value calculations. Note that peptide homology was addressed using razor rules in which a peptide matched to multiple different proteins was exclusively assigned to the protein that has more identified peptides. The output raw peptide intensity values from the Spectronaut detection software were noted. At this stage, any peptide that was not detected a minimum of 2 times across all of the samples and not detected in at least 50% of one of the biological groups was removed from further analysis. For remaining peptides, any missing data values were imputed using the following rules: 1) if less than 50% of signals were missing within a group, a randomized value near the average of the remaining values was calculated, or 2) if >50% of the signals were missing within the group, a randomized intensity within the bottom 1% of the detectable signals was used. Next, non-phosphorylated peptides and phosphopeptides which did not pass a 75% confidence of localization were removed from the analysis. The remaining phosphopeptides were then subjected to a robust mean normalization in which the highest and lowest 10% of the phosphopeptide signals were ignored and the average value of the remaining phosphopeptides was used as the normalized factor across all samples. We then summed all of the same precursor states for the same phosphopeptide (ie. different charge states) into a single modified phosphopeptide value and calculated the percent coefficient of variation across each group. Following database searching and peptide scoring using a library-based Spectronaut search, the data was annotated at a 1% peptide false discovery rate. A total of 103,664 phosphopeptides were identified in the dataset corresponding to 3,434 phosphoproteins.

### RNA sequencing-based transcriptome analysis

RNA extraction was performed from cells grown in the same manner as described for phosphoproteome analysis. Briefly, 10 ml cultures for five strains (wild-type H99, *cac1*Δ, *cac1*Δ::*CAC1*, *cac1*Δ::*CAC1^PP2C^*^Δ^, *cac1*Δ::*CAC1^ACC^*^Δ^) were grown in triplicates in two media conditions, YNB and YNB + 2% glucose. Cells were harvested, snap-frozen, and then lyophilised. The dry cell biomass was weighed and 0.05 g equivalent cells for each sample were used for RNA extraction using the mirVana™ miRNA Isolation Kit (Life Technologies, USA). The total RNA isolation was done following the kit guidelines and the extracted RNA was checked for quality via NanoDrop and gel electrophoresis. The RNA samples were quantified using Qubit and submitted to the Duke Sequencing and Genomic Technologies core facility. The library preparation was performed to select and sequence the PolyA mRNA pool (mRNA-seq). The libraries were then sequenced on Illumina NovaSeq X Plus to obtain 150-bp paired-end reads. Raw sequencing reads were subjected to quality control and adapter trimming using Cutadapt v2.4 in Python 3.7.4, employing the specified adapter sequence^107^. Subsequent read processing followed previously established protocols^108^. Cleaned reads were aligned to the *Cryptococcus neoformans* H99 reference genome using Hisat2 v2.2.1, which utilizes the Hisat and Bowtie2 algorithms. Genome annotation files were retrieved from the NCBI FTP repository. Hisat2 was executed with the parameters “-p 30” and “--dta –1,” along with default settings. The resulting alignment files were converted and sorted using Samtools v0.1.19, applying “-Sb –@ 8” for file conversion and “-@ 20” for sorting operations^109^. Gene-level expression matrices were generated and further analyzed using the R package IsoformSwitchAnalyzeR^110^. Data quality was evaluated with DEBrowser^111^, and DEGs were identified using DESeq2 v1.24^112^. DEGs were defined based on a threshold of > 2-fold change and a P-value < 0.05, and visualized using the EnhancedVolcano package in R v4.1.0^113^. For phosphoproteome analysis, the raw phosphopeptide intensity tables were imported into Perseus v1.6.15.0 for statistical analysis^114^. Intensities were first transformed to log scale and normalized by subtracting the median value of each sample to reduce systematic variation across runs. For pairwise comparisons between strains and media conditions, two-sided Student’s t-tests were performed in Perseus with permutation-based false discovery rate (FDR) control (FDR < 0.05). Significantly altered phosphorylation events were defined using a combined threshold of |log_2_ fold change| > 1 and *P* < 0.05. Volcano plots were generated via EnhancedVolcano package to visualize differentially phosphorylated peptides. Functional enrichment analysis was conducted for the set of significantly altered phosphoproteins or transcriptome. Gene Ontology (GO) term enrichment was performed using the DAVID^115^.

### Statistics and reproducibility

All experiments were performed with at least three independent biological replicates. For qRT-PCR, three biological replicates were analysed with three technical replicates each, which were averaged and not treated as independent measurements. For single-cell measurements (e.g., capsule thickness, titan cell size), the number of cells analysed (n) is indicated in the figure legends, and representative images are shown. Mouse group sizes are also indicated in the figure legends. Data are presented as mean ± SEM, with individual data points shown. Statistical analyses were performed using GraphPad Prism, employing Student’s *t*-test or one-way ANOVA followed by Tukey’s multiple-comparison test, as appropriate. Survival curves were compared using the log-rank (Mantel-Cox) test. For RNA sequencing, differentially expressed genes were defined as those with adjusted P < 0.05 and |log_2_FC| > 1; for phosphoproteomics, significantly altered phosphosites were defined as those with |log_2_FC| > 1 and adjusted P < 0.05. All tests were two-sided, and P < 0.05 was considered statistically significant.

## Data availability

All data are available in the manuscript and the Supporting Information files. The source data underlying the main figures and other supporting raw data files are deposited and stored in Figshare at: doi.org/10.6084/m9.figshare.30782315^116^. The RNA-seq data have been deposited in the Gene Expression Omnibus (GEO) database (accession number GSE312692), and the phosphoproteome data have been deposited with the ProteomeXchange Consortium via the PRIDE partner repository with the dataset identifier PXD069490. All other relevant data are available from the corresponding authors upon reasonable request. Requests should be directed to Yong-Sun Bahn.

## Acknowledgements

We thank the Duke University Proteomics and Metabolomics Core Facility for phosphoproteome data acquisition and the Duke University Sequencing and Genomic Technologies Core Facility for RNA sequencing.

## Contributions

All authors contributed to the manuscript and approved the final version. S.J. Yu, S.R. Yu, J.H. Jin, and Y.S. Bahn conceived and designed the study. J.H. Jin conducted domain analyses and generated mutant strains. M.K. Choi and H.S. Choi performed structural modelling and phosphatase activity assays. H.S. Cho supervised the structural and biochemical analyses and contributed to data interpretation and manuscript revision. S.J. Yu conducted all in vitro experiments with *C. neoformans* and gene expression analyses. V. Yadav prepared samples for, and performed the transcriptomic and phosphoproteomic experiments, and S.R. Yu analysed and visualised these datasets. E.S. Kim, H.W. Kim, J.S. Jeong, D.K. Kim, and K.T. Lee performed murine survival, fungal burden, histochemistry, immune cell, and cytokine analyses. J. Heitman contributed to interpretation of the findings and manuscript revision. S.J. Yu, S.R. Yu, and Y.S. Bahn prepared the initial draft, and all authors reviewed and revised the manuscript.

## Corresponding Authors

Correspondence to Yong-Sun Bahn, Kyung-Tae Lee, Hyun-Soo Cho, Joseph Heitman

