## Supplementary Information for "Protein phosphatase 2C domain contributes to the pathobiological function of adenylyl cyclase in *Cryptococcus neoformans*"

This PDF file includes:

**Supplementary figures 1-13**

**Supplementary tables 1-2**

**Supplementary image 1**

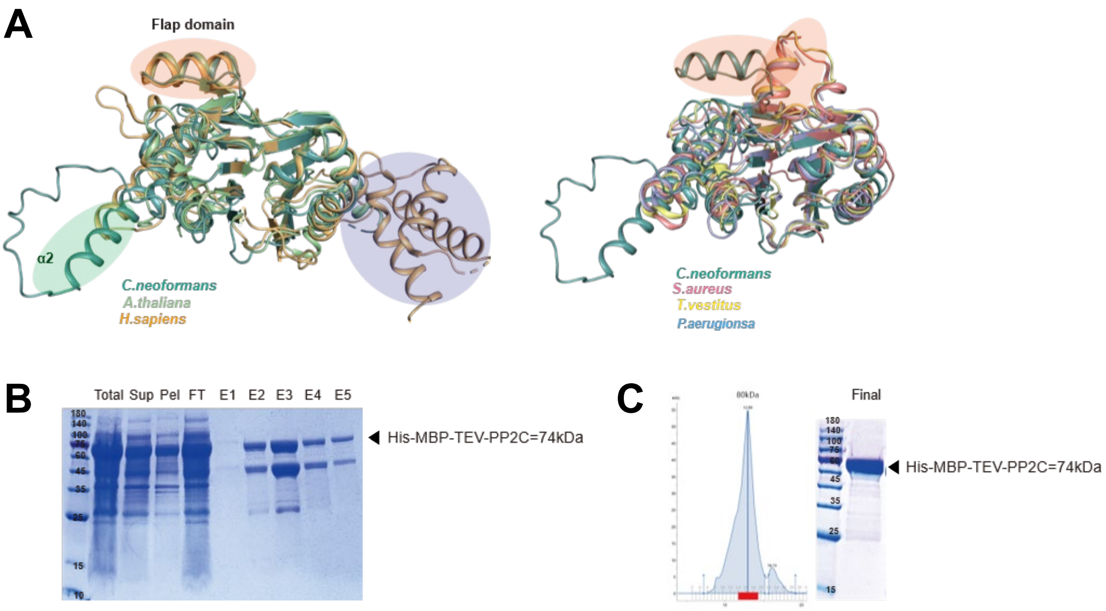


**Supplementary figure 1.** (A) Structural alignment of PP2C like domains. Overall PP2C fold were similar except flap domain (orange circle) and additional α2 helix (green circle) and missing the additional three alpha-helices at the C-terminus (purple circle). (green: *C. neoformans*, light green: *A. thaliana*, yellow: *H. sapiens*, lime: *T. vestitus*, blue: *P. aeruginosa*, salmon: *S.aureus*) (B) Coomassie blue stained SDS-PAGE gel of the Ni-affinity purified Cac1-PP2C domain. (C) Size exclusion chromatography and Coomassie blue stained SDS-PAGE gel of the purified PP2C domain.

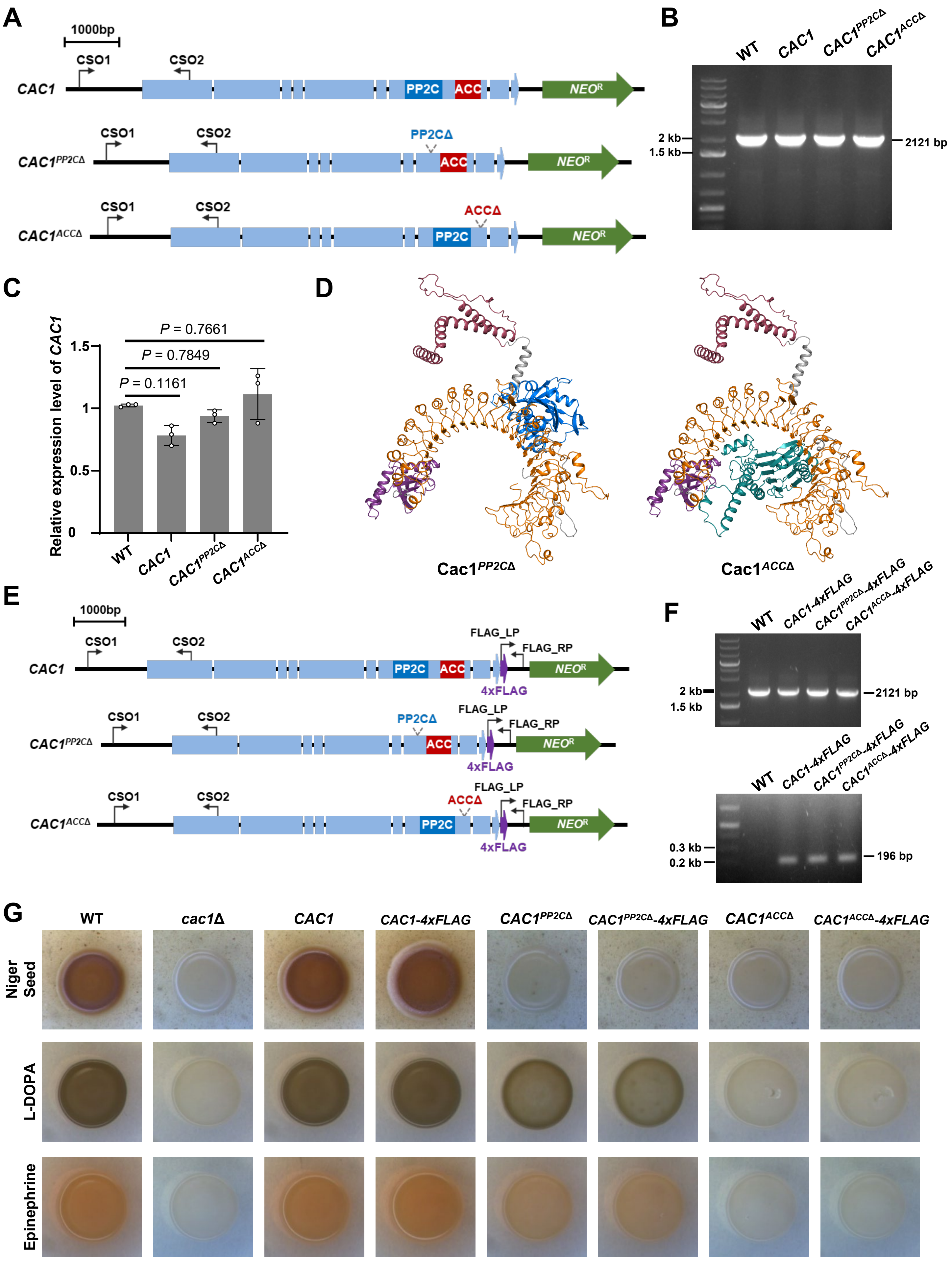
A
1000bp
CSO1
CSO2
CAC1
PP2C
ACC
NEO^R
CAC1^PP2CΔ
PP2CΔ
CAC1^ACCΔ
ACCΔ
B
WT
CAC1
CAC1^PP2CΔ
CAC1^ACCΔ
2 kb
1.5 kb
2121 bp
C
Relative expression level of CAC1
P = 0.7661
P = 0.7849
P = 0.1161
1.5
1
0.5
0
D
Cac1^PP2CΔ
Cac1^ACCΔ
E
FLAG_LP
FLAG_RP
4xFLAG
F
CAC1-4xFLAG
CAC1^PP2CΔ-4xFLAG
CAC1^ACCΔ-4xFLAG
0.3 kb
0.2 kb
196 bp
G
cac1Δ
Niger Seed
L-DOPA
Epinephrine

Continued

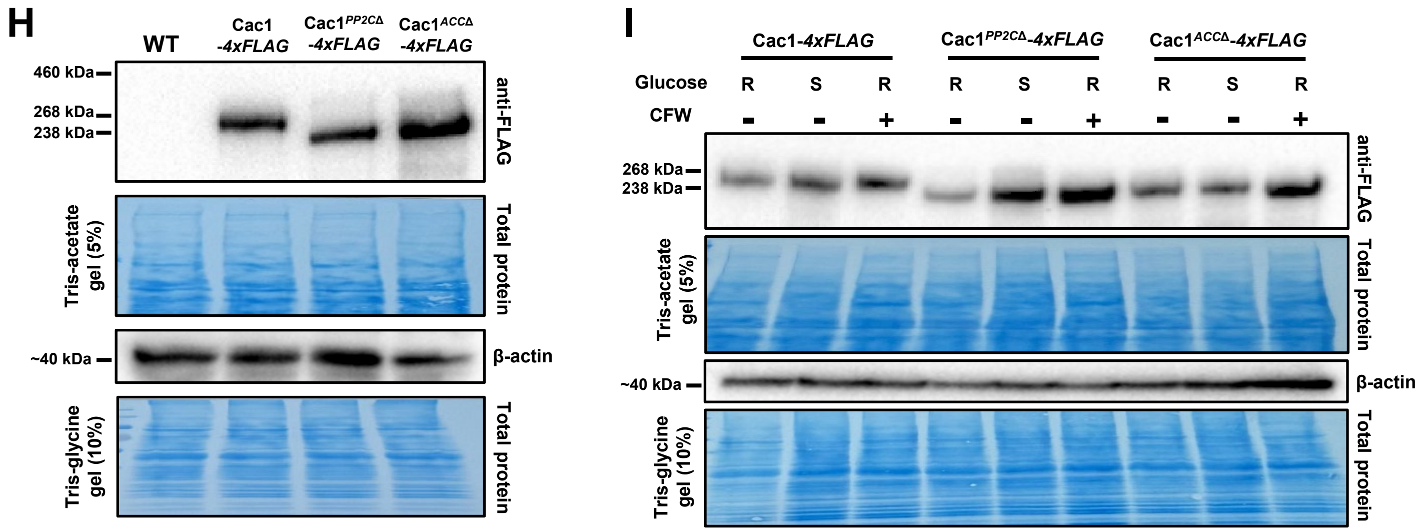


**Supplementary Figure 2. Confirmation of domain deletion, complementation, and FLAG-tagged strains through diagnostic PCR, qRT-PCR, and western blot analyses.** (A) Graphical representation of domains, deletion sites, and primer binding sites for the *CAC1*, $CAC1^{PP2C\Delta}$, and $CAC1^{ACC\Delta}$ alleles. (B) Diagnostic PCR was performed to confirm the targeted integration of *CAC1*, $CAC1^{PP2C\Delta}$, and $CAC1^{ACC\Delta}$ alleles into the native locus of *CAC1*. CSO1 and CSO2 primers were used, and expected band size is 2121 bp. (C) qRT-PCR analysis was performed to measure the relative expression levels of *CAC1* in the WT, *CAC1*, $CAC1^{PP2C\Delta}$, and $CAC1^{ACC\Delta}$ strains after cells were harvested at logarithmic phase ($OD_{600}$ of 0.8), confirming that the domain deletion strains express *CAC1* at levels comparable to the WT (one-way ANOVA with Tukey's multiple comparison test; *P* values indicated on the graph). (D) Structural models of $CAC1^{PP2C\Delta}$ and $CAC1^{ACC\Delta}$ predicted by AlphaFold3. Each domain is differently coloured (purple: Ras binding domain, orange: LRR domain, green: PP2C domain, blue: adenyly cyclase domain and marron: Cap binding domain), and an undefined region is shown in gray. (E) Graphical representation of domains, deletion sites, and primer binding sites for the 4xFLAG-tagged *CAC1*, $CAC1^{PP2C\Delta}$, and $CAC1^{ACC\Delta}$ alleles. (F) Diagnostic PCR was performed to confirm the targeted insertion of the 4xFLAG-tagged *CAC1*, $CAC1^{PP2C\Delta}$, and $CAC1^{ACC\Delta}$ alleles into the native locus of *CAC1*. CSO1 and CSO2 primers were used in the upper panel (expected band size: 2121 bp), and FLAG_LP and FLAG_RP primers were used in the lower panel (expected band size: 195 bp). (G) The indicated strains were spotted on Niger seed, L-DOPA, or epinephrine with 0.1% glucose. The plates were incubated at 30°C for 2 to 3 days and imaged. Representative colony pigmentation is shown. (H) Western blot analysis of the FLAG-tagged strains cultured to an $OD_{600}$ of 0.8 in YPD was performed to confirm that domain deletion does not compromise Cac1 protein stability. Cac1 was detected with an Anti-FLAG antibody, with β-actin and total protein (Coomassie brilliant blue-stained membrane) serving as loading controls. Expected sizes: 257 kDa for Cac1-4xFLAG, 227 kDa for $Cac1^{PP2C\Delta}$-4xFLAG, 236 kDa for $Cac1^{ACC\Delta}$-4xFLAG, and 42 kDA for β-actin. (I) Western blot analysis of the FLAG-tagged strains cultured under glucose-rich (R), nutrient starvation (S), and cell wall stress (250 μg/ml CFW, indicated with +) conditions. Anti-FLAG antibody was used to detect Cac1, with β-actin and total protein serving as loading controls. Uncropped and unprocessed images of gels and blots are provided in Supplementary Image 1.

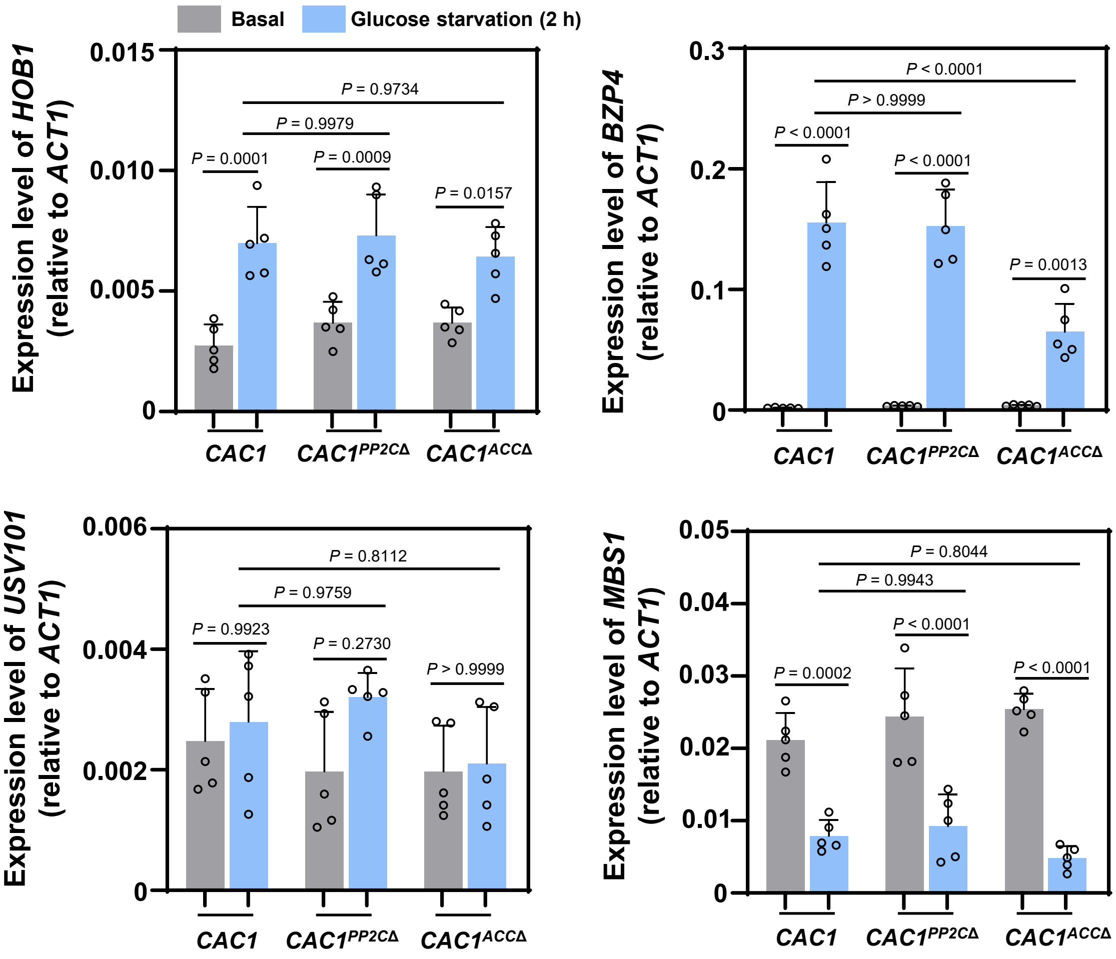


**Supplementary Figure 3. Expression of melanin-regulating genes were measured after glucose starvation.** Quantitative RT-PCR was performed using total RNA of each strain under nutrient rich (basal; YPD; shown in grey) or glucose starvation (YNB without amino acid and glucose; shown in blue) conditions. The expression levels of *HOB1*, *BZP4*, *USV101*, and *MBS1* were measured in *CAC1*, *CAC1*$^{PP2C\Delta}$, and *CAC1*$^{ACC\Delta}$ strains. Three biological replicates were performed with three technical replicates each, and the expression level of each gene was normalized to *ACT1* (ΔCt). Error bars indicate SEM, and statistical analysis was performed by one-way ANOVA with Tukey's multiple comparison test.

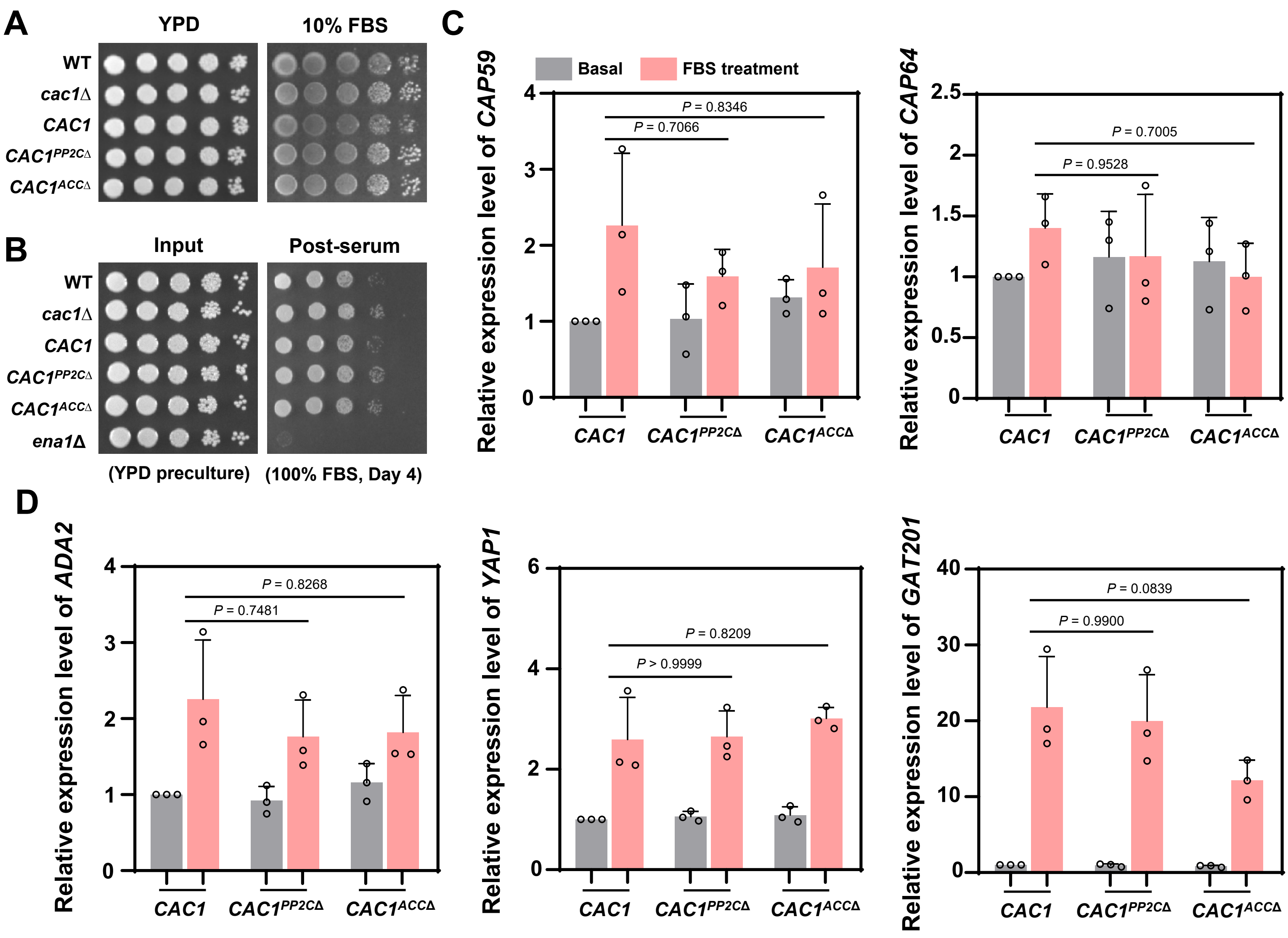


**Supplementary Figure 4. Growth and expression analyses were performed after FBS treatment.** (A) Spot dilution assay of indicated strains on YPD or 10% FBS agar media identical to the media used for FBS capsule analysis. The indicated strains were grown in liquid YPD for 16 h, washed with sterile PBS, serially diluted 10-fold, and spotted onto YPD or 10% FBS agar plates. The plates were incubated at 37°C and imaged after 3 days. (B) Serum survival assay of indicated cells before and after incubation in 100% FBS for 4 days. The indicated strains were grown in liquid YPD for 16 h, washed with sterile PBS, serially diluted 10-fold, and spotted onto YPD agar plates as the input. Next, the washed cells were 1/10 diluted into 100% heat-inactivated FBS. The cells were then incubated at 37°C with shaking for 4 days, serially diluted 10-fold, and spotted onto YPD agar plates. Both input and post-serum plates were incubated at 30°C for 2 days and imaged. (C) Quantitative RT-PCR was performed using total RNA of each strain under nutrient rich (basal; YPD; shown in grey) or 10% FBS induction condition for 6 h (shown in pink). The expression levels of *CAP59* and *CAP64* were measured in *CAC1*, *CAC1*$^{PP2C\Delta}$, and *CAC1*$^{ACC\Delta}$ strains. (D) Quantitative RT-PCR was performed using total RNA of each strain under nutrient rich (basal; YPD; shown in grey) or 10% FBS condition for 2h (shown in pink). The expression levels of *ADA2*, *YAP1*, and *GAT201* were measured in *CAC1*, *CAC1*$^{PP2C\Delta}$, and *CAC1*$^{ACC\Delta}$ strains. Three biological replicates were performed with three technical replicates each, and relative transcript levels were calculated using the $2^{-\Delta\Delta Ct}$ method. Error bars indicate SEM, and statistical analysis was performed by one-way ANOVA with Tukey's multiple comparison test.

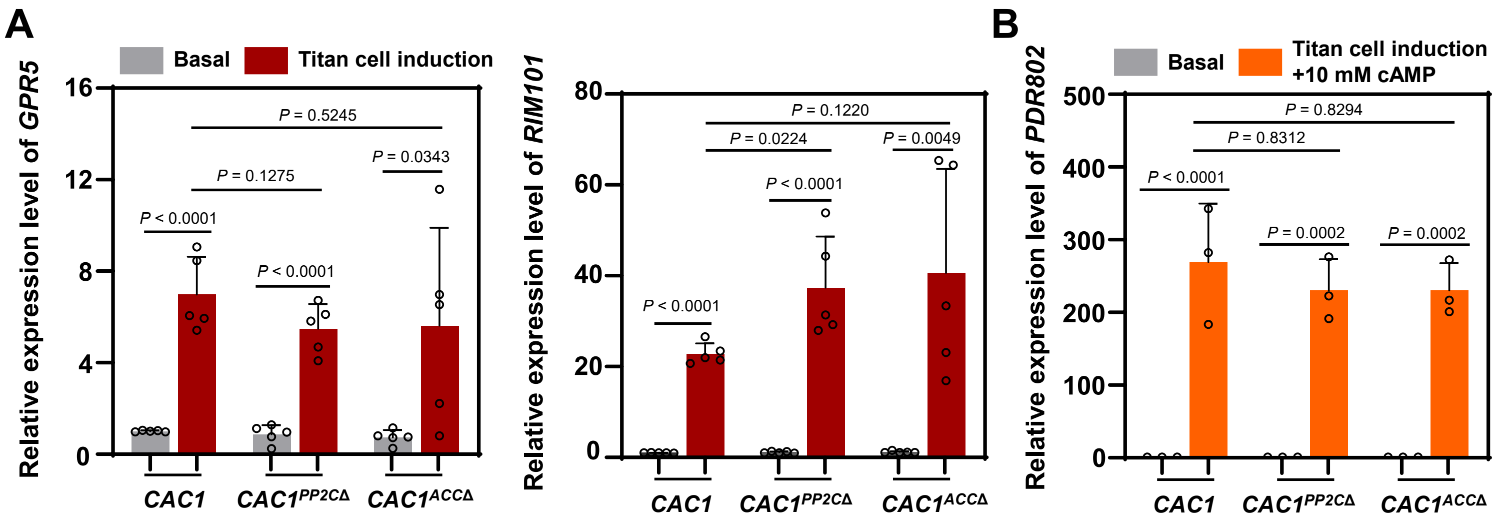


**Supplementary Figure 5. Expression of titan cell-regulating genes were measured after titan cell induction.** Quantitative RT-PCR was used to measure relative transcript levels in strains grown in YPD (basal; grey bard) compared to the titan cell-inducing conditions. (A) Expression of *GPR5* and *RIM101* in *CAC1*, $CAC1^{PP2C\Delta}$, and $CAC1^{ACC\Delta}$ strains following standard titan cell induction (red bars; conditions detailed in Materials and Methods). Three biological replicates were performed with three technical replicates each, and relative transcript levels were calculated using the $2^{-\Delta\Delta Ct}$ method. Error bars indicate SEM, and statistical analysis was performed using an unpaired two-tailed Student's t-test. (B) Expression of *PDR802* following titan cell induction supplemented with 10 mM cAMP (orange bars). Three biological replicates were performed with three technical replicates each, and relative transcript levels were calculated using the $2^{-\Delta\Delta Ct}$ method. Error bars indicate SEM, and statistical analysis was performed using one-way ANOVA with Tukey's multiple comparison test.

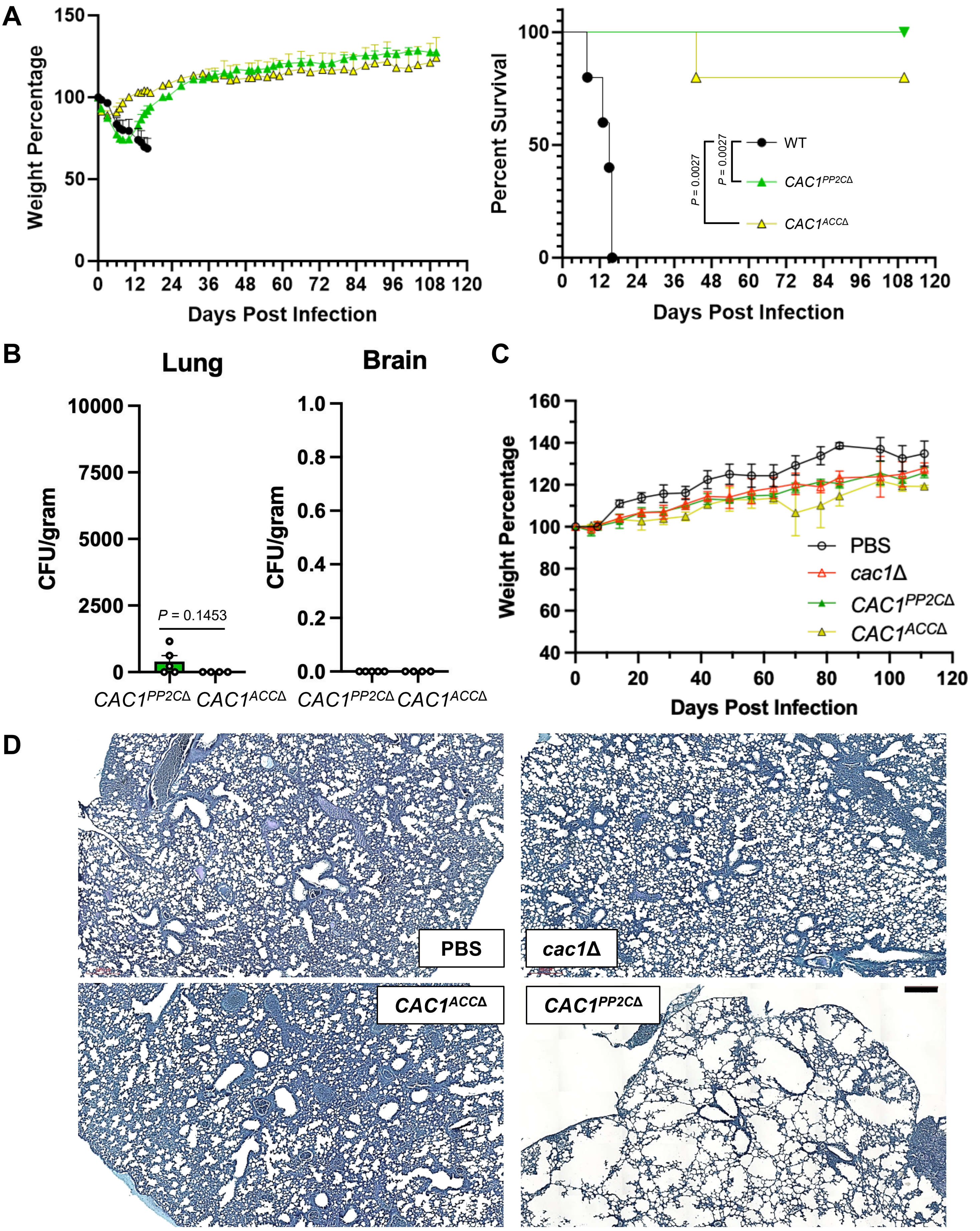
A
Weight Percentage
Days Post Infection
Percent Survival
WT
CAC1PP2CΔ
CAC1ACCΔ
P = 0.0027
P = 0.0027
B
Lung
Brain
CFU/gram
P = 0.1453
CAC1PP2CΔ CAC1ACCΔ
CAC1PP2CΔ CAC1ACCΔ
C
Weight Percentage
Days Post Infection
PBS
cac1Δ
CAC1PP2CΔ
CAC1ACCΔ
D
PBS
cac1Δ
CAC1ACCΔ
CAC1PP2CΔ

Continued

E

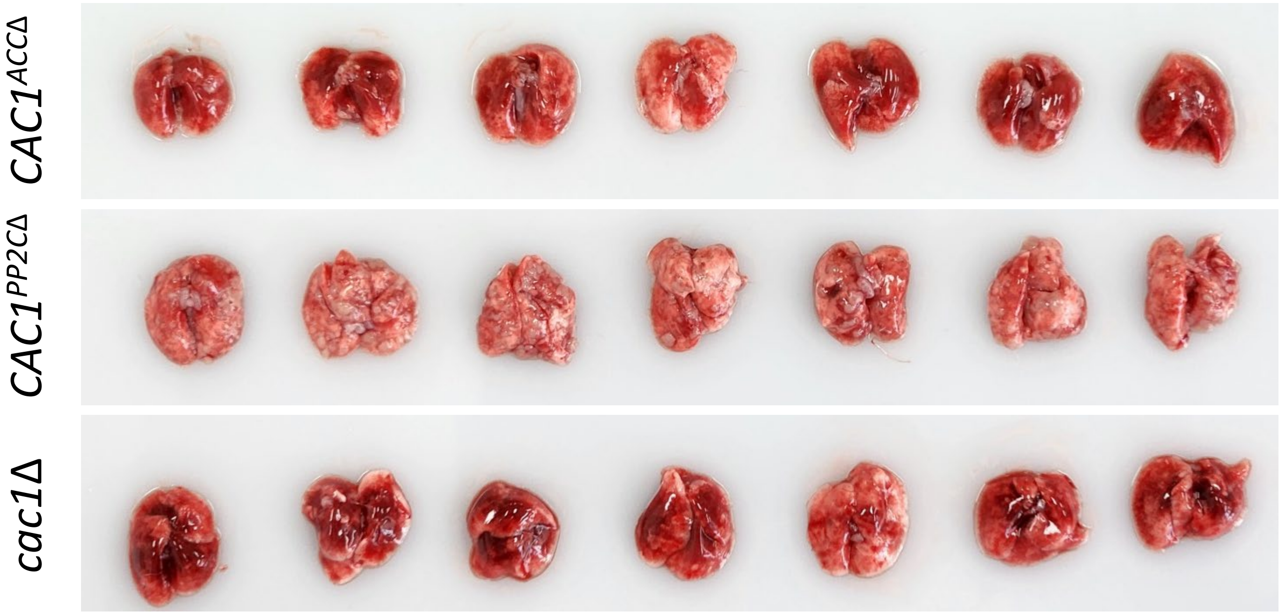


**Supplementary Figure 6. Long-term survival analysis.** Balb/c mice were infected intranasally with $5 \times 10^5$ cells of *Cryptococcus neoformans* strains and sacrificed for each analysis. (A-B) Long-term survival assay (n=5). Body weight changes of infected mice and survival curves analyzed by the log-rank (Mantel–Cox) test, with corresponding *P*-values: WT vs $CAC1^{PP2C\Delta}$ (0.0027); WT vs $CAC1^{ACC\Delta}$ (0.0027). (B) Fungal burden (colony-forming units) in lung and brain tissues was determined at 111 days post-infection. Error bars indicate SEM, and statistical analysis was performed using an unpaired two-tailed Student’s t-test with Welch’s correction. For brain fungal burden, statistical comparison was not performed because all values were zero in both groups. (C-D) Additional cohort (n=2) used for histopathological analysis. Body weight changes of these mice (C) and corresponding lung sections stained with PAS at 111 dpi (D) are shown. Scale bar indicates 200 µm. (E) Lung images obtained from all seven mice in the *cac1*Δ, $CAC1^{PP2C\Delta}$, and $CAC1^{ACC\Delta}$ infection groups are shown to illustrate the reproducibility and range of pulmonary lesions. Lungs from all groups were collected at 26 days post-infection.

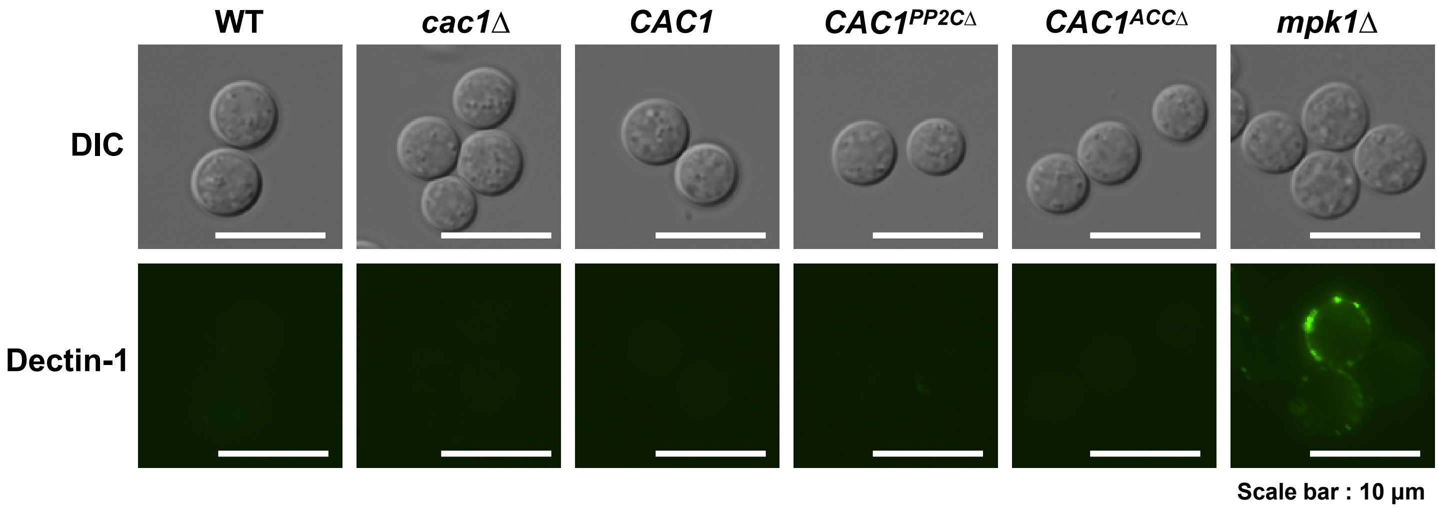


**Supplementary Figure 7. Dectin-1 staining revealed that β-1,3-glucans remained masked and undetectable in the *CAC1* and domain deletion strains.** Dectin-1 staining using soluble human Dectin-1a fused to an IgG1 Fc domain. Indicated strains were cultured for 16 h in liquid YPD at 30°C. The cells were then stained with 15 μg/ml of Fc-hDectin-1a for 1 h, washed twice, and stained with 10 μg/ml of Alexa Fluor 488-conjugated Goat Anti-human IgG in the dark for 1 h. The cells were washed twice and imaged using DIC and fluorescence microscopy. Scale bar, 10 μm.

A

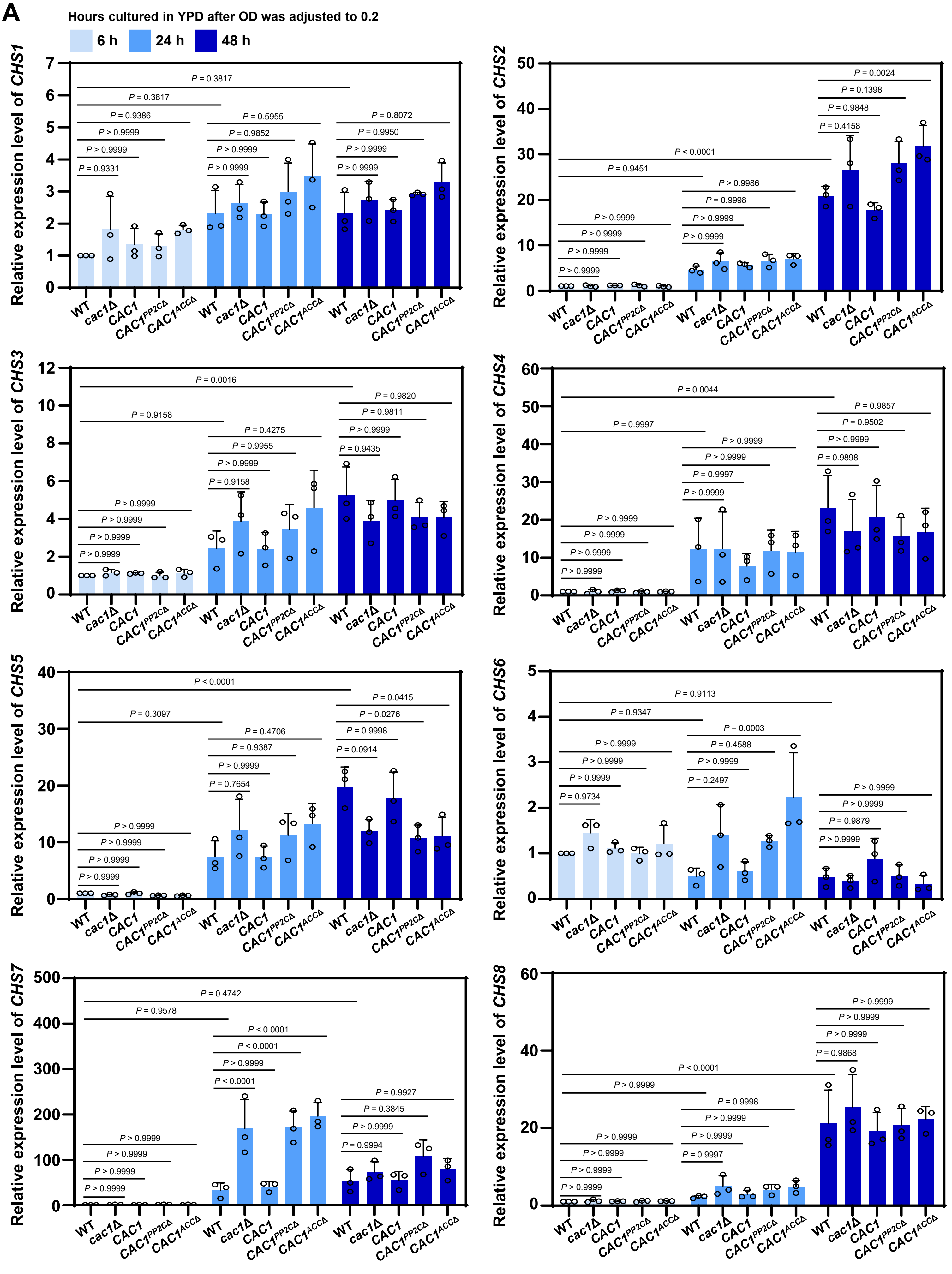

Hours cultured in YPD after OD was adjusted to 0.2
6 h
24 h
48 h
Relative expression level of CHS1
Relative expression level of CHS2
Relative expression level of CHS3
Relative expression level of CHS4
Relative expression level of CHS5
Relative expression level of CHS6
Relative expression level of CHS7
Relative expression level of CHS8
WT
cac1Δ
CAC1
CAC1PP2CΔ
CAC1ACCΔ


Continued

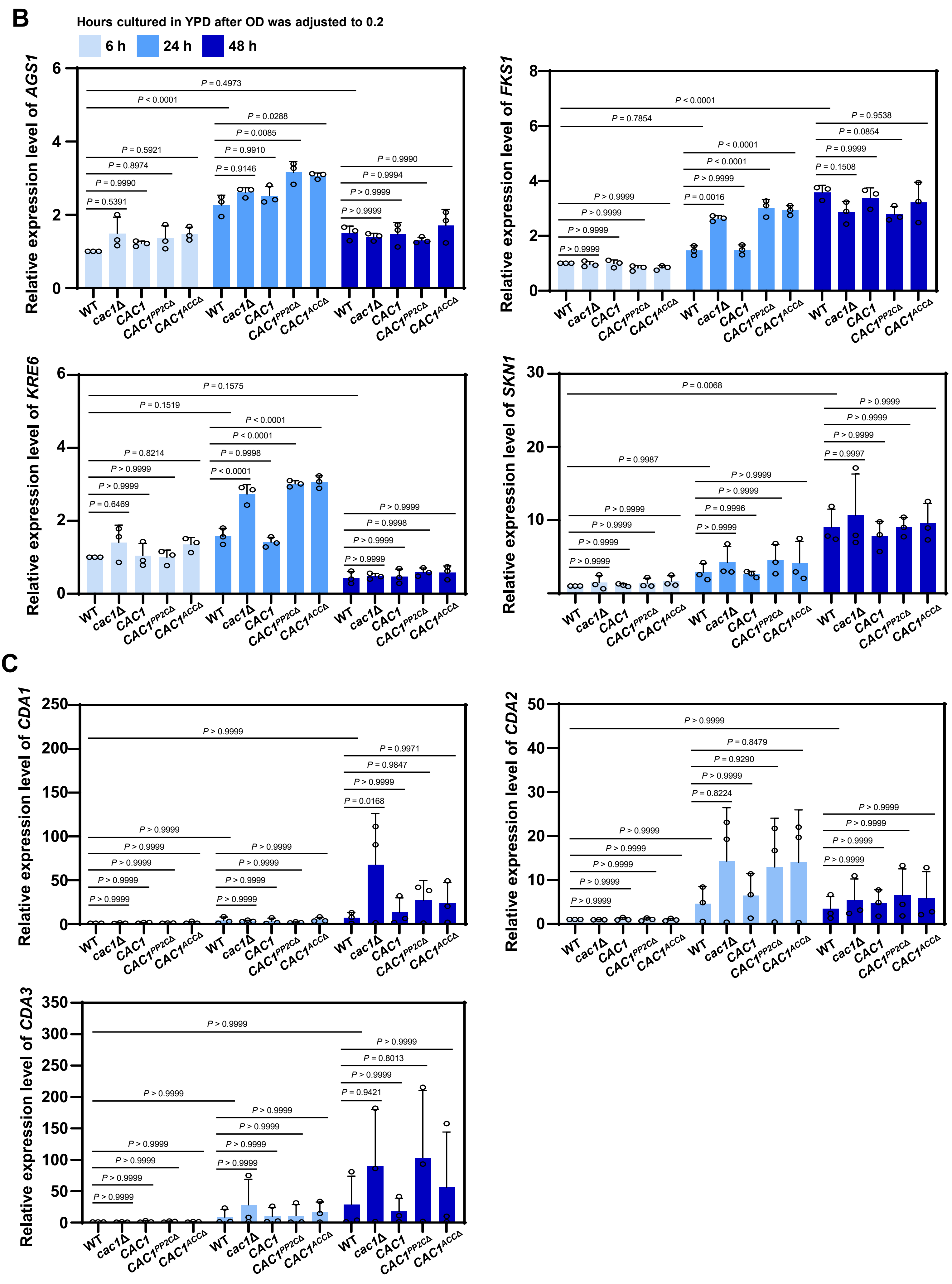
B
Hours cultured in YPD after OD was adjusted to 0.2
6 h
24 h
48 h
Relative expression level of AGS1
P = 0.4973
P < 0.0001
P = 0.0288
P = 0.0085
P = 0.5921
P = 0.9910
P = 0.8974
P = 0.9146
P = 0.9990
P = 0.9990
P = 0.9994
P > 0.9999
P = 0.5391
P > 0.9999
Relative expression level of FKS1
P < 0.0001
P = 0.7854
P = 0.9538
P = 0.0854
P = 0.9999
P = 0.1508
P < 0.0001
P < 0.0001
P > 0.9999
P = 0.0016
P > 0.9999
P > 0.9999
P > 0.9999
P > 0.9999
Relative expression level of KRE6
P = 0.1575
P = 0.1519
P < 0.0001
P < 0.0001
P = 0.8214
P = 0.9998
P > 0.9999
P < 0.0001
P > 0.9999
P = 0.6469
P > 0.9999
P = 0.9998
P > 0.9999
P > 0.9999
Relative expression level of SKN1
P = 0.0068
P > 0.9999
P > 0.9999
P > 0.9999
P = 0.9997
P = 0.9987
P > 0.9999
P > 0.9999
P = 0.9996
P > 0.9999
P > 0.9999
P > 0.9999
P > 0.9999
P > 0.9999
C
Relative expression level of CDA1
P > 0.9999
P = 0.9971
P = 0.9847
P > 0.9999
P = 0.0168
P > 0.9999
P > 0.9999
P > 0.9999
P > 0.9999
P > 0.9999
P > 0.9999
P > 0.9999
P > 0.9999
P > 0.9999
Relative expression level of CDA2
P > 0.9999
P = 0.8479
P = 0.9290
P > 0.9999
P = 0.8224
P > 0.9999
P > 0.9999
P > 0.9999
P > 0.9999
P > 0.9999
P > 0.9999
P > 0.9999
P > 0.9999
P > 0.9999
Relative expression level of CDA3
P > 0.9999
P > 0.9999
P = 0.8013
P > 0.9999
P = 0.9421
P > 0.9999
P > 0.9999
P > 0.9999
P > 0.9999
P > 0.9999
P > 0.9999
P > 0.9999
P > 0.9999
P > 0.9999
WT
cac1Δ
CAC1
CAC1PP2CΔ
CAC1ACCΔ

Continued

**Supplementary Figure 8. Expression of genes involved in cell wall remodeling.** Quantitative RT-PCR was performed using total RNA of each strain under nutrient rich conditions. The cells were cultured for 16 h in liquid YPD at 30°C, synchronized to $OD_{600nm}$ of 0.2, and incubated at 30°C for 6, 24, or 48 h. (A) The expression levels of chitin synthase genes (*CHS1*, *CHS2*, *CHS3*, *CHS4*, *CHS5*, *CHS6*, *CHS7,* and *CHS8*) were measured in WT, *cac1*Δ, *CAC1*, $CAC1^{PP2C\Delta}$, and $CAC1^{ACC\Delta}$ strains. (B) The expression levels of *AGS1*, *FKS1*, *KRE6*, and *SKN1* were measured in the indicated strains. (C) The expression levels of chitin deacetylases (*CDA1*, *CDA2*, and *CDA3*) were measured in the indicated strains. For all genes, three biological replicates were performed with three technical replicates each, and relative transcript levels were calculated using the $2^{-\Delta\Delta Ct}$ method. Error bars indicate SEM, and statistical analysis was performed by one-way ANOVA with Tukey's multiple comparison test (*, $P < 0.05$; **, $P < 0.01$; ***, $P < 0.001$; ****, $P < 0.0001$; NS, not significant).

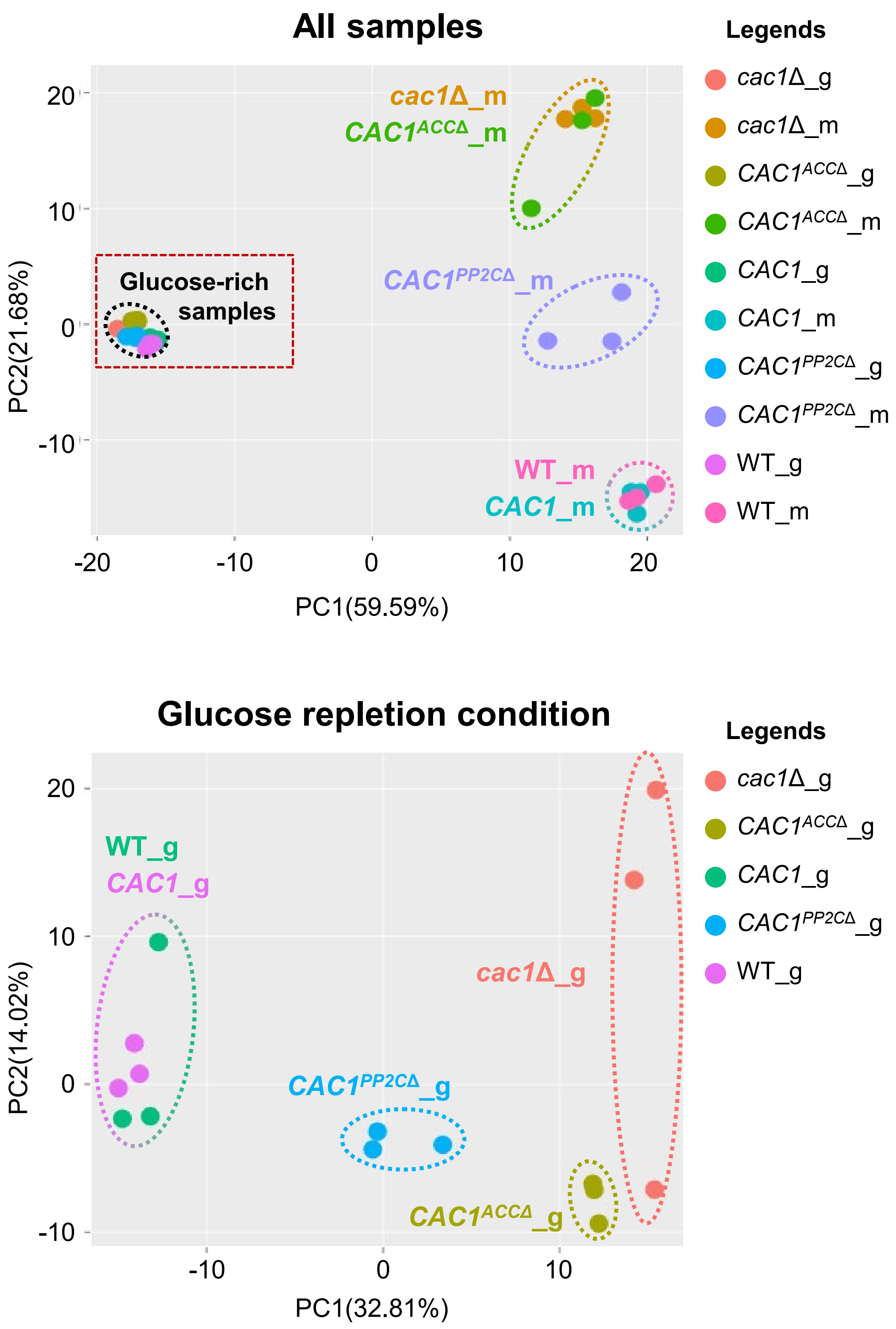


**Supplementary figure 9. Principal component analysis of transcriptome profiles across glucose-rich and minimal media conditions.** PCA was performed using the R package *debrowser* to visualize transcriptomic similarities and differences among *C. neoformans* strains, including wild-type (WT), *cac1*Δ, complemented strain (*CAC1*), and domain-deletion mutants (*CAC1*$^{PP2C\Delta}$ and *CAC1*$^{ACC\Delta}$). Each dot represents an individual biological replicate. Samples grown in minimal medium (m) and glucose-rich medium (g) (bottom) are shown.

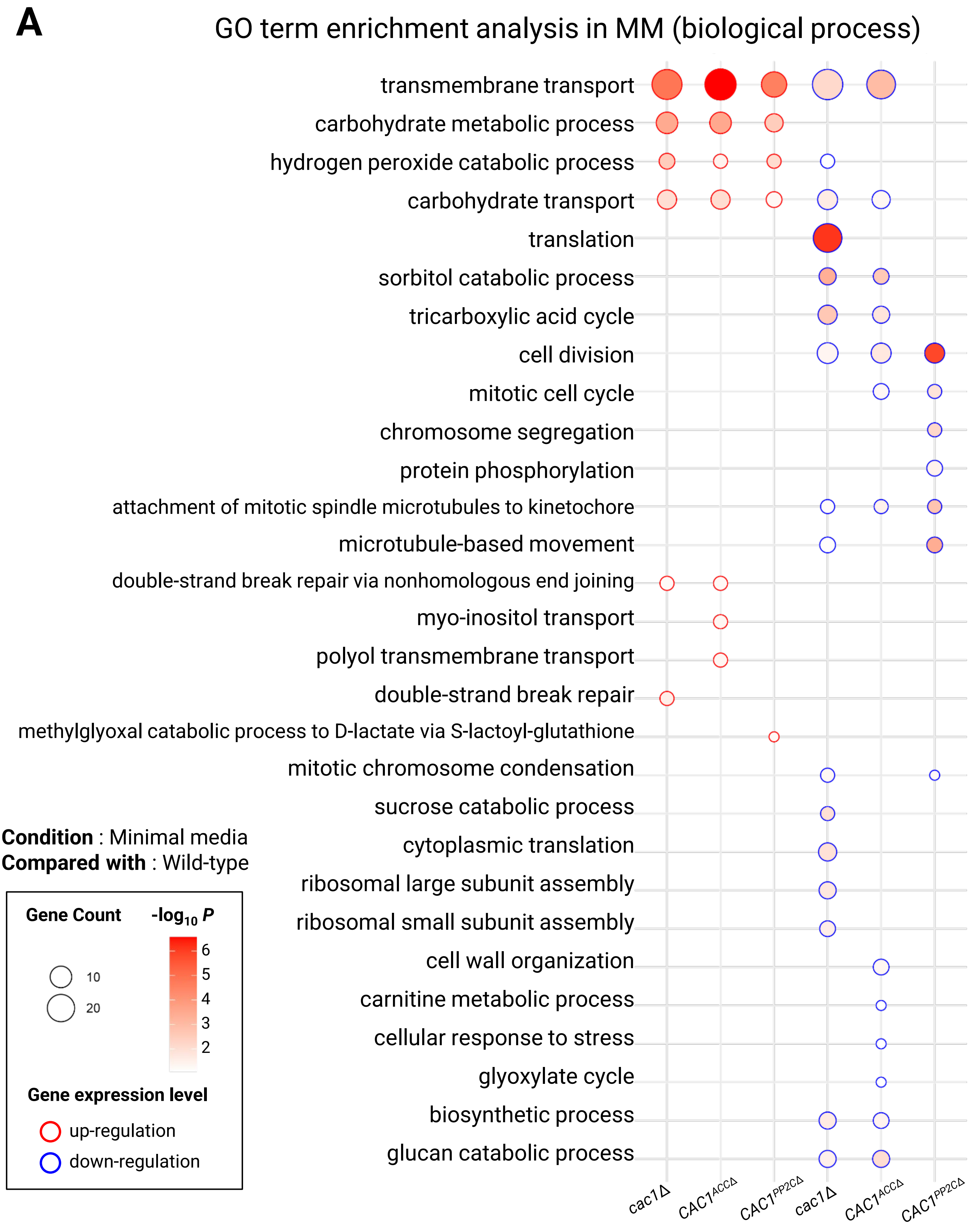
A
GO term enrichment analysis in MM (biological process)
transmembrane transport
carbohydrate metabolic process
hydrogen peroxide catabolic process
carbohydrate transport
translation
sorbitol catabolic process
tricarboxylic acid cycle
cell division
mitotic cell cycle
chromosome segregation
protein phosphorylation
attachment of mitotic spindle microtubules to kinetochore
microtubule-based movement
double-strand break repair via nonhomologous end joining
myo-inositol transport
polyol transmembrane transport
double-strand break repair
methylglyoxal catabolic process to D-lactate via S-lactoyl-glutathione
mitotic chromosome condensation
sucrose catabolic process
cytoplasmic translation
ribosomal large subunit assembly
ribosomal small subunit assembly
cell wall organization
carnitine metabolic process
cellular response to stress
glyoxylate cycle
biosynthetic process
glucan catabolic process
cac1Δ
CAC1ACCΔ
CAC1PP2CΔ
cac1Δ
CAC1ACCΔ
CAC1PP2CΔ
Condition : Minimal media
Compared with : Wild-type
Gene Count
10
20
-log10 P
6
5
4
3
2
Gene expression level
up-regulation
down-regulation

Continued

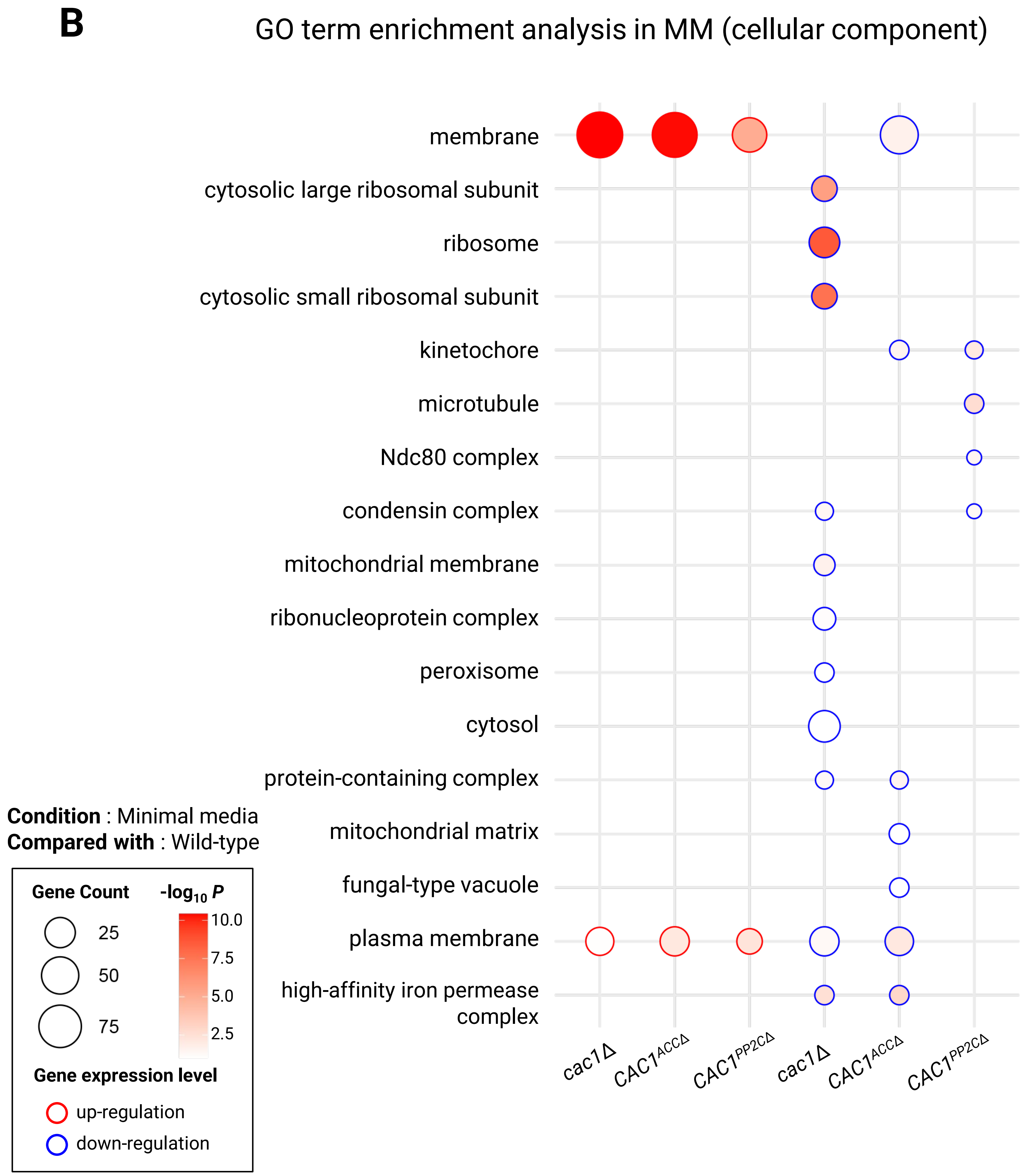

B
GO term enrichment analysis in MM (cellular component)
membrane
cytosolic large ribosomal subunit
ribosome
cytosolic small ribosomal subunit
kinetochore
microtubule
Ndc80 complex
condensin complex
mitochondrial membrane
ribonucleoprotein complex
peroxisome
cytosol
protein-containing complex
mitochondrial matrix
fungal-type vacuole
plasma membrane
high-affinity iron permease complex
cac1Δ
CAC1ACCΔ
CAC1PP2CΔ
cac1Δ
CAC1ACCΔ
CAC1PP2CΔ
Condition : Minimal media
Compared with : Wild-type
Gene Count
25
50
75
-log10 P
10.0
7.5
5.0
2.5
Gene expression level
up-regulation
down-regulation


Continued

C

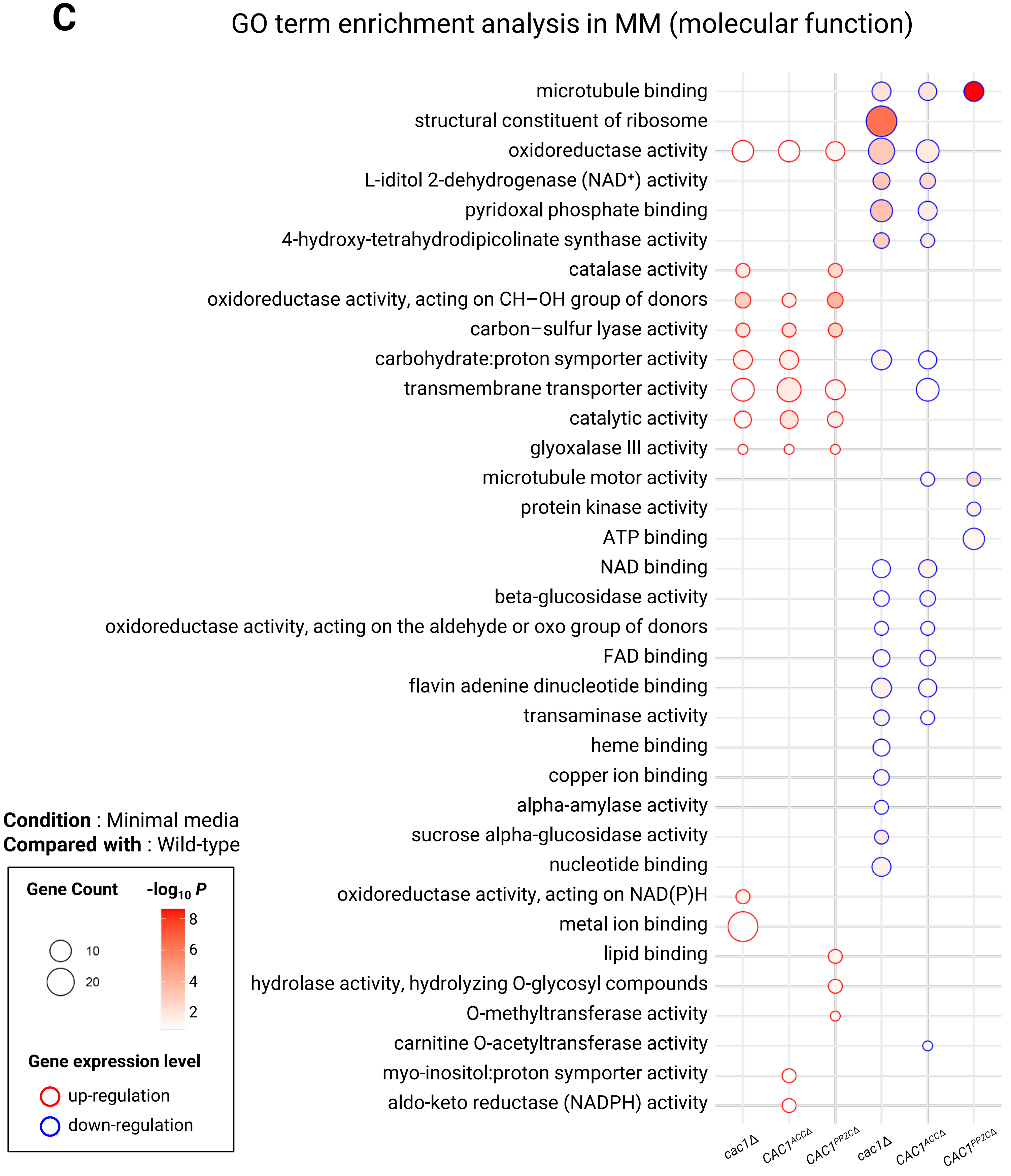
GO term enrichment analysis in MM (molecular function)
microtubule binding
structural constituent of ribosome
oxidoreductase activity
L-iditol 2-dehydrogenase (NAD⁺) activity
pyridoxal phosphate binding
4-hydroxy-tetrahydrodipicolinate synthase activity
catalase activity
oxidoreductase activity, acting on CH−OH group of donors
carbon−sulfur lyase activity
carbohydrate:proton symporter activity
transmembrane transporter activity
catalytic activity
glyoxalase III activity
microtubule motor activity
protein kinase activity
ATP binding
NAD binding
beta-glucosidase activity
oxidoreductase activity, acting on the aldehyde or oxo group of donors
FAD binding
flavin adenine dinucleotide binding
transaminase activity
heme binding
copper ion binding
alpha-amylase activity
sucrose alpha-glucosidase activity
nucleotide binding
oxidoreductase activity, acting on NAD(P)H
metal ion binding
lipid binding
hydrolase activity, hydrolyzing O-glycosyl compounds
O-methyltransferase activity
carnitine O-acetyltransferase activity
myo-inositol:proton symporter activity
aldo-keto reductase (NADPH) activity
cac1Δ
CAC1ACCΔ
CAC1PP2CΔ
cac1Δ
CAC1ACCΔ
CAC1PP2CΔ
Condition : Minimal media
Compared with : Wild-type
Gene Count
10
20
-log10 P
8
6
4
2
Gene expression level
up-regulation
down-regulation

Continued

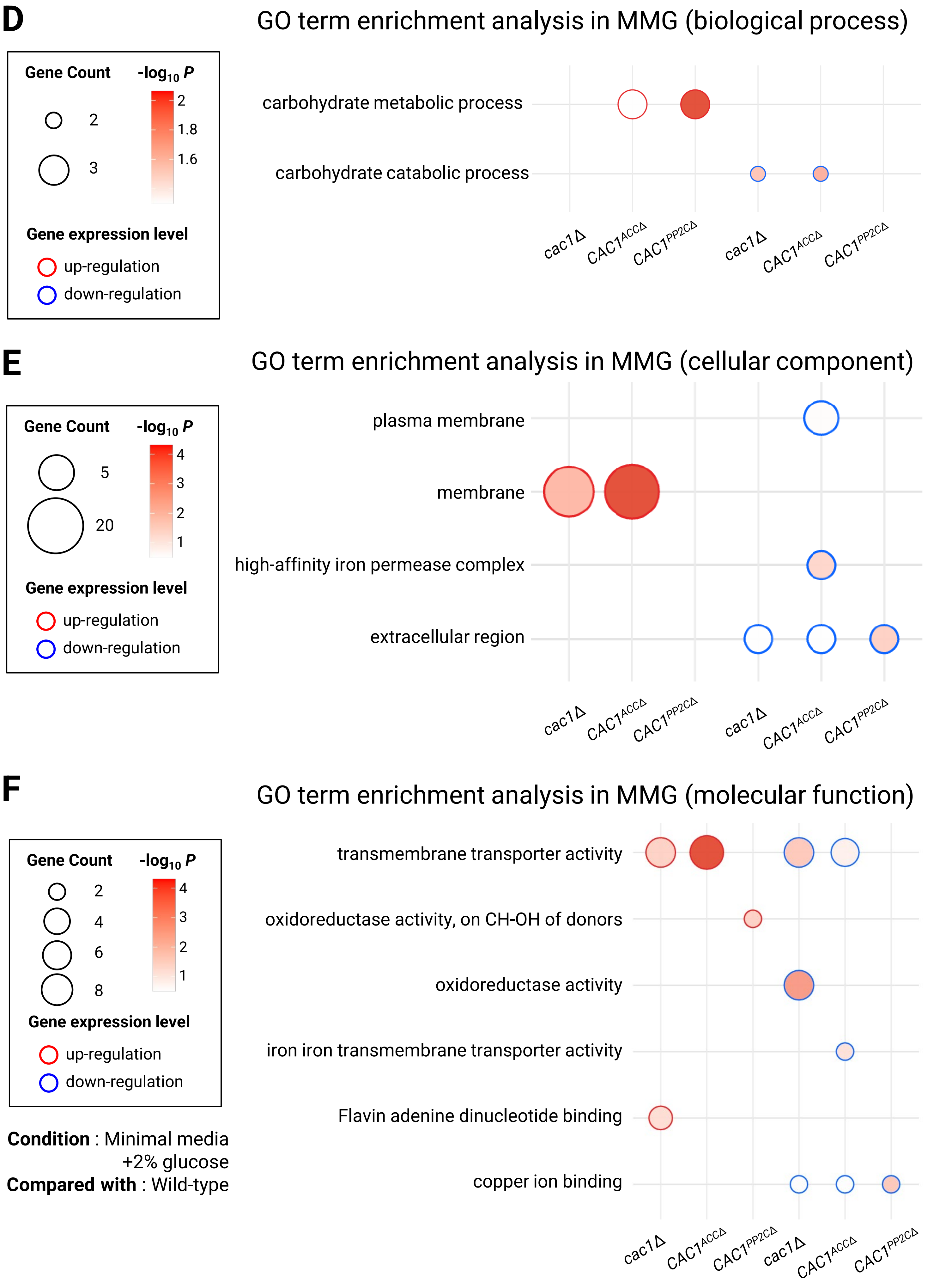

D
GO term enrichment analysis in MMG (biological process)
Gene Count
-log10 P
2
3
2
1.8
1.6
Gene expression level
up-regulation
down-regulation
carbohydrate metabolic process
carbohydrate catabolic process
cac1Δ
CAC1ACCΔ
CAC1PP2CΔ
cac1Δ
CAC1ACCΔ
CAC1PP2CΔ
E
GO term enrichment analysis in MMG (cellular component)
Gene Count
-log10 P
5
20
4
3
2
1
Gene expression level
up-regulation
down-regulation
plasma membrane
membrane
high-affinity iron permease complex
extracellular region
cac1Δ
CAC1ACCΔ
CAC1PP2CΔ
cac1Δ
CAC1ACCΔ
CAC1PP2CΔ
F
GO term enrichment analysis in MMG (molecular function)
Gene Count
-log10 P
2
4
6
8
4
3
2
1
Gene expression level
up-regulation
down-regulation
transmembrane transporter activity
oxidoreductase activity, on CH-OH of donors
oxidoreductase activity
iron iron transmembrane transporter activity
Flavin adenine dinucleotide binding
copper ion binding
cac1Δ
CAC1ACCΔ
CAC1PP2CΔ
cac1Δ
CAC1ACCΔ
CAC1PP2CΔ
Condition : Minimal media +2% glucose
Compared with : Wild-type


Continued

**Supplementary figure 10. GO term enrichment analysis of differentially expressed genes in *CAC1* domain-deletion mutants**. (A,D) Biological process, (B,E) cellular component, (C,F) molecular function terms. Bubble plots show Gene Ontology (GO) enrichment results for upregulated and downregulated genes in *cac1*Δ, $CAC1^{ACC\Delta}$, and $CAC1^{PP2C\Delta}$ strains. (A-C) is for minimal media (MM), and (D-F) is for minimal media + 2% glucose condition (MMG). The color gradient of the bubbles represents the $-\log_{10} P$, while bubble size indicates the number of genes associated with each GO term. Red bubbles denote enriched terms among upregulated genes, and blue bubbles denote those among downregulated genes.

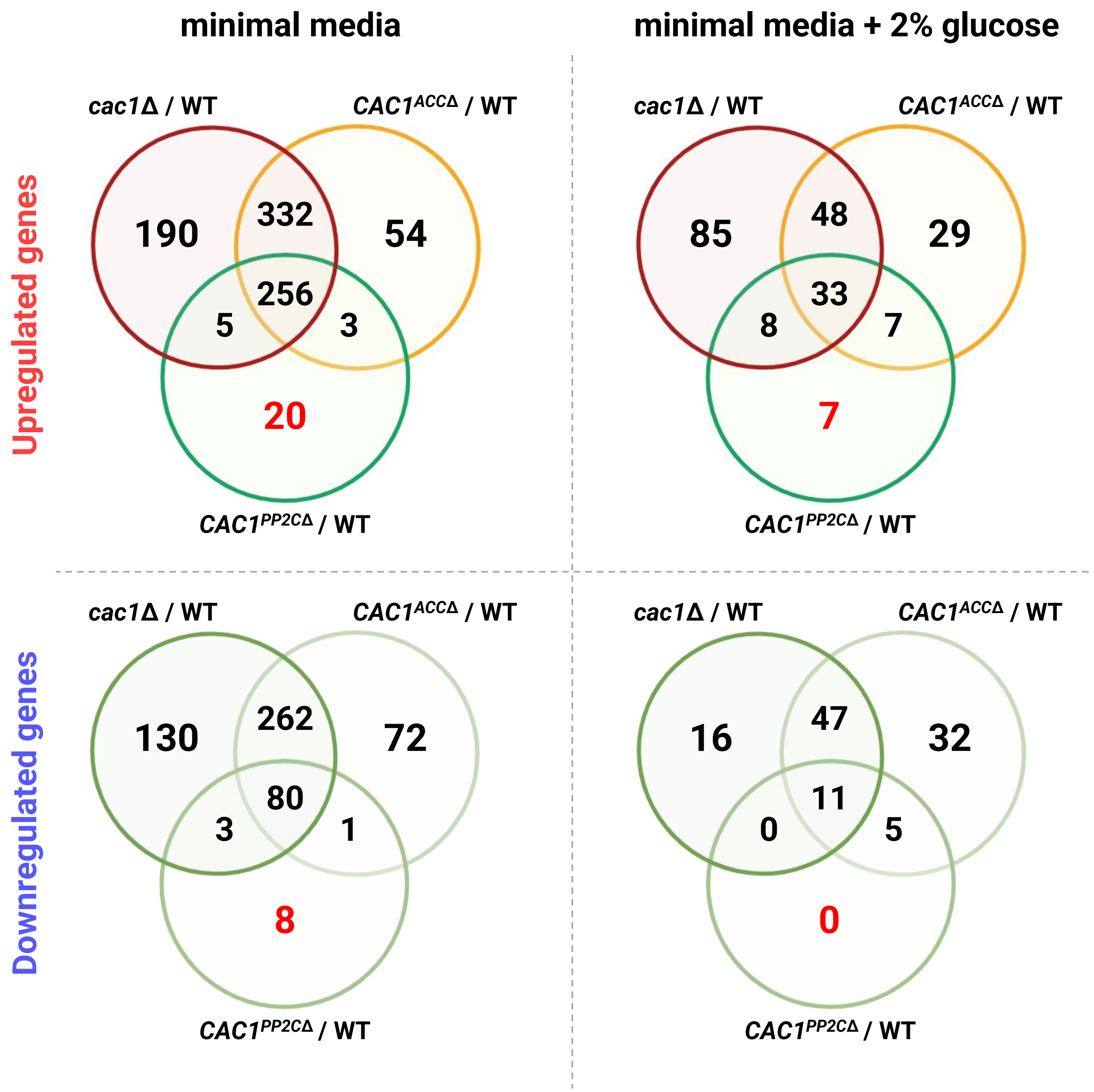


**Supplementary figure 11. Venn diagram analysis of differentially expressed genes in *CAC1* domain-deletion mutants.** Venn diagrams show the overlap of significantly upregulated (top) and downregulated (bottom) genes among the *CAC1* full deletion mutant (*cac1*Δ), the PP2C domain deletion mutant (*CAC1*$^{PP2C\Delta}$), and the adenylyl cyclase catalytic domain deletion mutant (*CAC1*$^{ACC\Delta}$). Analyses were performed separately for cells grown in minimal medium (left) and glucose-rich medium (right). Differential expression was defined by $|\log_2 FC| \geq 1$ and adjusted $P < 0.05$. The numbers indicate the count of shared or unique differentially expressed genes between strains under each condition.

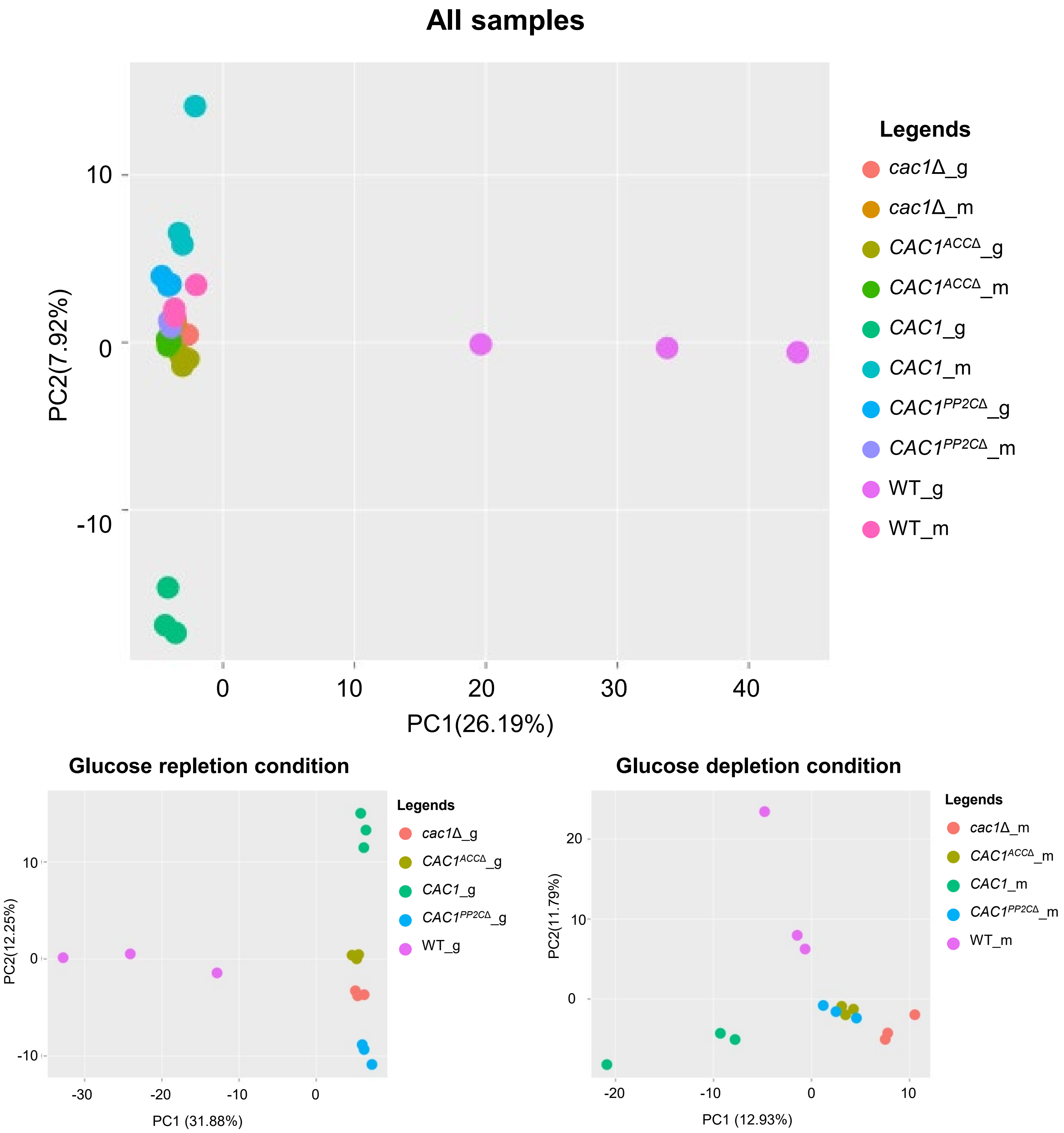


**Supplementary figure 12. Principal component analysis of transcriptome profiles across glucose-rich and minimal media conditions.** PCA was performed using the R package *debrowser* to visualize phosphoproteomic similarities and differences among *C. neoformans* strains, including wild-type (WT), *cac1*Δ, complemented strain (*CAC1*), and domain-deletion mutants (*CAC1*$^{PP2C\Delta}$ and *CAC1*$^{ACC\Delta}$). Each dot represents an individual biological replicate. Samples grown in minimal medium (m) and glucose-rich medium (g) (bottom) are shown.

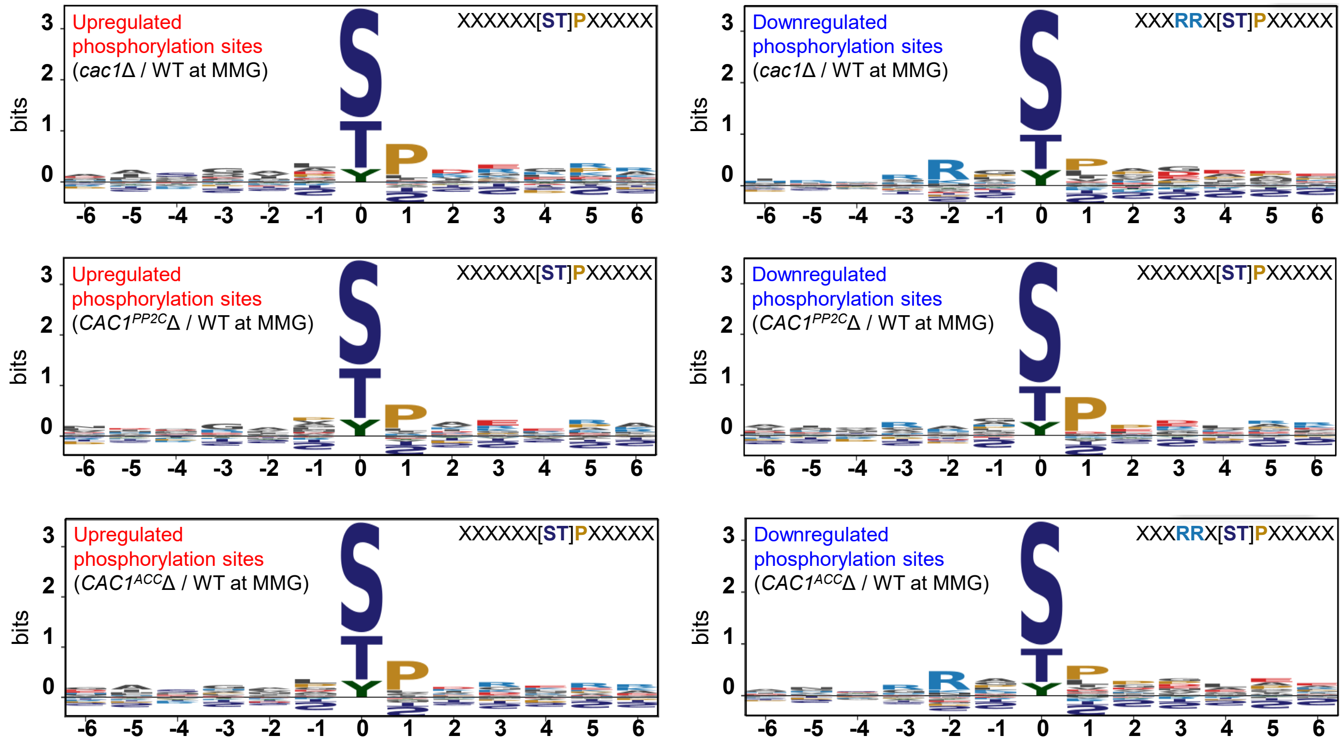


**Supplementary figure 13. Sequence motif analysis of differentially regulated phosphorylation sites in *C. neoformans* Cac1 mutants under MMG condition.** Phosphorylation motifs were analyzed for sites that were upregulated (left, red labels) or downregulated (right, blue labels) in *cac1Δ*, *CAC1*$^{PP2CΔ}$, and *CAC1*$^{ACCΔ}$ strains compared with wild type (WT) under minimal medium supplemented with 2% glucose (MMG). Sequence logos were generated to represent the amino acid frequencies surrounding phosphorylated serine/threonine residues (position 0). Distinct enrichment patterns are observed between upregulated and downregulated sites, including proline-directed motifs ([S/T]P) in upregulated sites and basic residue-enriched motifs (Rxx[S/T]) in downregulated sites.

**Supplementary Table 1. List of strains used in this study.**

| Strain | Genotype | Parent | Reference |
|---|---|---|---|
| H99 | *MATα* | | (1) |
| KN99**a** | *MAT***a** | | (2) |
| YSB3814 | *MAT*α *mpk1*Δ::*NAT*-STM #240 | H99 | (3) |
| AI167 | *MAT*α *ena1*Δ::*NAT* | H99 | (4) |
| YSB5650 | *MAT*α *cac1*Δ::*NAT*-STM #159 | H99 | (5) |
| YSB7885 | *MAT*α *cac1*Δ::$CAC1^{PP2C\Delta}$-*NEO* | YSB5650 | This study |
| YSB8184 | *MAT*α *cac1*Δ::$CAC1^{ACC\Delta}$-*NEO* | YSB5650 | This study |
| YSB8189 | *MAT*α *cac1*Δ::*CAC1*-4xFLAG-*NEO* | YSB5650 | This study |
| YSB8183 | *MAT*α *cac1*Δ::$CAC1^{PP2C\Delta}$-4xFLAG-*NEO* | YSB5650 | This study |
| YSB8192 | *MAT*α *cac1*Δ::$CAC1^{ACC\Delta}$-4xFLAG-*NEO* | YSB5650 | This study |
| *Escherichia coli* BL21 (DE3) | λ(DE3), T7 RNA polymerase | *Escherichia coli B* | (6) |


1. Perfect JR, Ketabchi N, Cox GM, Ingram CW, Beiser CL. 1993. Karyotyping of *Cryptococcus neoformans* as an epidemiological tool. *J Clin Microbiol* 31:3305-9.
2. Nielsen K, Cox GM, Wang P, Toffaletti DL, Perfect JR, Heitman J. 2003. Sexual cycle of *Cryptococcus neoformans* var. grubii and virulence of congenic **a** and alpha isolates. Infect Immun. 71(9):4831-41.
3. Lee KT, So YS, Yang DH, et al. 2016. Systematic functional analysis of kinases in the fungal pathogen *Cryptococcus neoformans*. *Nat Commun* 7, 12766.
4. Idnurm A, Walton FJ, Floyd A, Reedy JL, Heitman J. 2009. Identification of *ENA1* as a virulence gene of the human pathogenic fungus *Cryptococcus neoformans* through signature-tagged insertional mutagenesis. *Eukaryot Cell* 8.
5. Jin JH, Lee KT, Hong J, et al. 2020. Genome-wide functional analysis of phosphatases in the pathogenic fungus *Cryptococcus neoformans. Nat Commun* 11, 4212.
6. Studier, F. W., and Moffatt, B. A. 1986. Use of bacteriophage T7 RNA polymerase to direct selective high-level expression of cloned genes. *J. Mol. Biol*. 189, 113–130.

**Supplementary Table 2. List of primers used in this study.**

| Primer Name | Sequence (5' to 3') | Comment |
|---|---|---|
| **B11623** | CGGCCGCCAGTGTGATGGATAAAAG AGCAGGATGGGAAG | *CAC1* complementation primer LP |
| **B11624** | CAAGGGCGAATTCTGCAGATGTCTG AACTTGAAGGGAATG | *CAC1* complementation primer RP |
| **B11660** | CAGCTGTAATAATCTTCGATGTTGGA TTGTTAAC | *CAC1* split primer MRP |
| **B11661** | ATCGAAGATTATTACAGCTGGGCCA TAC | *CAC1* split primer MLP |
| **B11729** | CATTGACAATCTTGCCATGATGGTCA TGGTGGTG | *CAC1* PP2C domain deletion LP |
| **B11730** | CATTCACCACCATGACCATCATGGC AAGATTGTC | *CAC1* PP2C domain deletion RP |
| **B11731** | CCGACTCCAGCAAGAGTTGAAGAGG GAAGTTATGG | *CAC1* ACC domain deletion LP |
| **B11732** | CCATAACTTCCCTCTTCAACTCTTGC TGGAGTCGG | *CAC1* ACC domain deletion RP |
| **B11771** | GTCATTGGTTCTTTTGCGG | *CAC1* sequencing primer 1 |
| **B11772** | CCTCTAAATCACCACCATTCAC | *CAC1* sequencing primer 2 |
| **B11773** | GCAGGTTTATCGCCCTCTAC | *CAC1* sequencing primer 3 |
| **B11774** | CGGACAACAACAACATCGTC | *CAC1* sequencing primer 4 |
| **B11775** | GACGAATCTGCCATCTGAG | *CAC1* sequencing primer 5 |
| **B11776** | TGCTGTGGTTTACCTCGTG | *CAC1* sequencing primer 6 |
| **B11777** | GTGTCTTTCCAATCTGTTGC | *CAC1* sequencing primer 7 |
| **B11778** | GTGGGGAGGATTGAAAAC | *CAC1* sequencing primer 8 |
| **B11779** | GGACTGTAAAAGATGCCTCG | *CAC1* complementation screening primer LP |
| **B11780** | CTTGCGTTTCCGACAATG | *CAC1* complementation screening primer RP |
| **B6567** | GCATGCGGCGCGCCAGAT | FLAG_LP |
| **B11822** | GCCAACATCTCTTCTTTCG | FLAG_RP |
| **B679** | CGCCCTTGCTCCTTCTTCTATG | *ACT1* qRT primer qLP |
| **B680** | GACTCGTCGTATTCGCTCTTCG | *ACT1* qRT primer qRP |
| **B15938** | CACCCTTTGGAAGTTGTGG | *LAC1* qRT primer qLP |
| **B15939** | TGATAATTGCAGAGTACCG | *LAC1* qRT primer qRP |
| **B8640** | ATTCATTCCCGATTGGCG | *CAP10* qRT primer qLP |
| **B7163** | GAGAACCAAACAGACGACG | *CAP10* qRT primer qRP |
| **B8684** | GCTATTAGAGGCTACAAGCG | *CAP59* qRT primer qLP |
| **B8685** | GGGTGAACAACCTATCGTG | *CAP59* qRT primer qRP |
| **B8643** | ACGCTATGAACGAAGAGGC | *CAP60* qRT primer qLP |
| **B8644** | GGAGTGAAAACAGAGTTGGG | *CAP60* qRT primer qRP |
| **B8645** | CAAGGAAAGGGCATTCAGAG | *CAP64* qRT primer qLP |
| **B8646** | TCAGAAAGCATTGCCTGG | *CAP64* qRT primer qRP |
| **B23102** | CCCACTCCCGGTCTCAAATA | *BZP4* qRT primer qLP |

| | | |
|---|---|---|
| **B23103** | CAGTAGGCGAGACAGGTGAT | *BZP4* qRT primer qRP |
| **B23106** | TGGGTCCGAGAACATTGTGT | *GAT201 qRT primer qLP* |
| **B23107** | CTGCCTCCAATCTCGCTTTC | *GAT201* qRT primer qRP |
| **B23108** | CGCCTAGACCTTCACTCGAT | *PDR802* qRT primer qLP |
| **B23109** | CTTTGGTGGCAGAAAGGGAC | *PDR802* qRT primer qRP |
| **B22028** | AGGGTGAAAGAGAGGGAAC | *ADA2* qRT primer qLP |
| **B22029** | GCTTTTTTGCGTTGCTGG | *ADA2* qRT primer qRP |
| **B15946** | CAGAATGCAAGCCAGAATCA | *YAP1* qRT primer qLP |
| **B15947** | GCGTTCATGCTGTTGTTGTT | *YAP1* qRT primer qRP |
| **B9368** | CCGCCGGTATTGTTAAGATG | *CHS1* qRT primer qLP |
| **B9369** | CCCAAAAGTACCGCTATCCA | *CHS1* qRT primer qRP |
| **B9370** | GCAACTTGGTGATGGATGTG | *CHS2* qRT primer qLP |
| **B9371** | AGTACCGCATGCTGTCCATT | *CHS2* qRT primer qRP |
| **B9372** | CCAAGGGTTTTCGGACTACA | *CHS3* qRT primer qLP |
| **B9373** | TCACGATAATGCCAGAGACG | *CHS3* qRT primer qRP |
| **B9374** | ACGCCCATCATATTCTGCTC | *CHS4* qRT primer qLP |
| **B9375** | TTCTCCTTGACGACCATTCC | *CHS4* qRT primer qRP |
| **B9376** | CAAGGGTGGTGGAAGAAGAA | *CHS5* qRT primer qLP |
| **B9377** | ATAACCTGATCCTGCCCAGTC | *CHS5* qRT primer qRP |
| **B9378** | GGCCCCCTCTTATGACTACC | *CHS6* qRT primer qLP |
| **B9379** | TGATCTTCCCCTTGGAGTTG | *CHS6* qRT primer qRP |
| **B9380** | CGTTCCTTACGACTGATCCA | *CHS7* qRT primer qLP |
| **B9381** | TCTCAAGAAATCGGCTCCTG | *CHS7* qRT primer qRP |
| **B12142** | ACGTTTGCGTCCTTCTTGAC | *CHS8* qRT primer qLP |
| **B12143** | AATGCGACAATCTCACCACA | *CHS8* qRT primer qRP |
| **B14791** | CTGCCGCAGCTATTTTTAGG | *AGS1* qRT primer qLP |
| **B14792** | CGATGATGGGGAAGTAGGAA | *AGS1* qRT primer qRP |
| **B14793** | TCGAATGTCCCCTAACCAAG | *FKS1* qRT primer qLP |
| **B14794** | CAAGTCGAGTTGAGCAGCAA | *FKS1* qRT primer qRP |
| **B14795** | TGGACGTTGGTGTTTTCAGA | *SKN1* qRT primer qLP |
| **B14796** | CCAGTGGGCCAATAGTGAAT | *SKN1* qRT primer qRP |
| **B14799** | GAAGACCATCCAGGTCCAGA | *KRE6* qRT primer qLP |
| **B14800** | GGGAGACTTCTCCTCGGTTC | *KRE6* qRT primer qRP |
| **B11629** | CTGACTCTACTGTGGCCCTC | *CDA1* qRT primer qLP |
| **B11630** | TCACCATCGCCGATTATCCA | *CDA1* qRT primer qRP |
| **B11631** | GTGATTGGACCGGTGTGAAC | *CDA2* qRT primer qLP |
| **B11632** | CACTGGTTTGCAAGCCGATA | *CDA2* qRT primer qRP |
| **B11633** | TGCCAAATGTTCCAGTAGCT | *CDA3* qRT primer qLP |
| **B11634** | TCTTTCACCGATCAATGCCG | *CDA3* qRT primer qRP |

| | | |
|---|---|---|
| **PP2C_5BamHI** | CG GGATCC TTG GGCTCCATTGACAATCTTGCCATG | *CAC1* PP2C pET cloning primer 5' |
| **PP2C_3XhoI** | CCCTCGAGTCAGAACAAATCAGACA CATTCACCAC | *CAC1* PP2C pET cloning primer 3' |
| ***IFNG*_F** | ACAGCAAGGCGAAAAGGATG | *IFNG* qRT primer qLP |
| ***IFNG*_R** | TGGTGGACCACTCGGATGA | *IFNG* qRT primer qRP |
| ***IL4*_F** | ATCATCGGCATTTTGAACGAGG | *IL4* qRT primer qLP |
| ***IL4*_R** | TGCAGCTCCATGAGAACACTA | *IL4* qRT primer qRP |
| ***IL5*_F** | CTCTGTTGACAAGCAATGAGACG | *IL5* qRT primer qLP |
| ***IL5*_R** | TCTTCAGTATGTCTAGCCCCTG | *IL5* qRT primer qRP |
| ***GAPDH*_F** | TGATGGTGTGAACCACGAG | *GAPDH* qRT primer qLP |
| ***GAPDH*_R** | GGCATGGACTGTGGTCATGA | *GAPDH* qRT primer qRP |

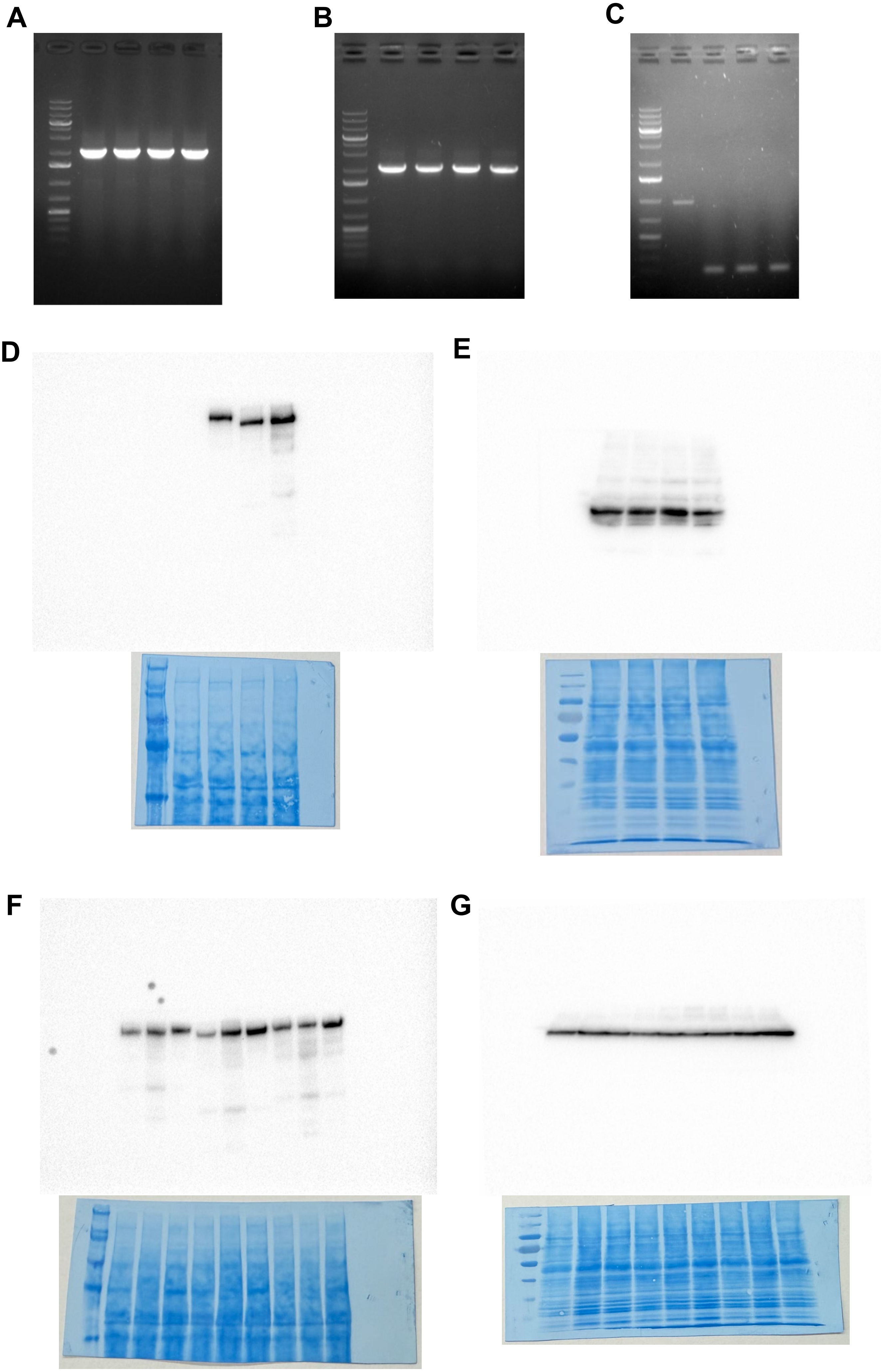


Continued

**Supplementary image 1**. **Uncut and unmodified gel and blot images.** (A) Fig. S2B – Diagnostic PCR for *CAC1*, *CAC1*$^{PP2C\Delta}$, and *CAC1*$^{ACC\Delta}$ strains. (B) Fig. S2F, upper panel – 5' diagnostic PCR of 4xFLAG-tagged *CAC1*, *CAC1*$^{PP2C\Delta}$, and *CAC1*$^{ACC\Delta}$ strains. (C) Fig. S2F, lower panel – 3' diagnostic PCR for 4xFLAG-tagged *CAC1*, *CAC1*$^{PP2C\Delta}$, and *CAC1*$^{ACC\Delta}$ strains. (D, E) Fig. S2H – Immunoblotting of 4xFLAG-tagged *CAC1*, *CAC1*$^{PP2C\Delta}$, and *CAC1*$^{ACC\Delta}$ using anti-FLAG (D) and anti-actin (E) antibodies. Corresponding membranes stained with Coomassie brilliant blue (CBB) are shown below each blot. (F, G) Fig. S2I – Immunoblotting of FLAG-tagged strains cultured under glucose-rich, nutrient starvation, and cell wall stress conditions using anti-FLAG (F) and anti-actin (G) antibodies. Corresponding membranes stained with CBB are shown below each blot. Diagnostic PCR and immunoblot images were exported directly from Bio-Rad Image Lab as TIFF files without further image processing.
